# SPRINT-MS: A high-throughput platform for identifying protein-protein interactions using pooled IP-MS and sparse signal recovery

**DOI:** 10.1101/2025.11.21.689547

**Authors:** Lena A. Street, Moitrish Majumdar, Katherine L. Rothamel, Esin Gogus, Gloria Brar, Gene W. Yeo, Tomas Rube, Marko Jovanovic

**Affiliations:** Department of Biological Sciences, Columbia University, New York, NY, USA; Applied Mathematics Department, University of California Merced, Merced, CA, USA; Department of Cellular and Molecular Medicine, University of California San Diego, La Jolla, CA, USA; Institute for Genomic Medicine, University of California San Diego, La Jolla, CA, USA; Department of Molecular and Cellular Biology, University of California Berkeley, Berkeley, CA, USA

## Abstract

We present SPRINT-MS (SParse Reconstruction of INTeractions by Mass Spectrometry), an integrated experimental and computational platform to accelerate the discovery of protein-protein interactions (PPIs). PPIs, which govern critical cellular and physiological processes such as development and disease, form extensive networks that vary across time, conditions, and cell types, creating a complex, high-dimensionality problem. Thus, there is a pressing need for universally applicable tools capable of mapping and quantifying PPI networks and their context-dependent dynamics with high efficiency. SPRINT-MS combines an innovative antibody (or lysate) pooling scheme, immunopurification-mass spectrometry (IP-MS), and a novel sparse signal reconstruction algorithm to enable pooled PPI capture experiments. This approach increases throughput by an order of magnitude, while reducing sample input requirements. We demonstrate that SPRINT-MS, applied to 30 bait proteins of interest via either antibody or lysate pooling, is comparable to standard individual IP-MS experiments in the identification of PPIs and recapitulation of known interactions.

## Introduction

Interactions between proteins are highly diverse and govern numerous cellular and physiological processes, such as development and disease states. Because protein-protein interaction (PPI) networks change with time, condition, and cell type, mapping them requires repeated measurement across each context, making it an exceptionally demanding experimental problem. Consequently, there is demand for universally applicable experimental tools capable of comprehensively assessing PPIs. In contrast to our ability to globally measure changes in RNA and protein levels at scale, current methods for studying PPIs, including yeast-two-hybrid,^1^ phage display,^2^ proximity labeling assays,^3–7^ protein correlation profiling followed by mass spectrometry,^8–11^ and immunopurification followed by mass spectrometry (IP-MS),^12–16^ are limited in that they cannot offer both high sensitivity and throughput (**Supplemental Figure 1a**). Of these methods, IP-MS is widely used as it provides quantitative results while requiring no modified cell lines (if an antibody against your protein of interest is available) or specialized equipment (beyond access to a mass spectrometry facility). Furthermore, its reliability and high sensitivity, as well as its proven record, makes it the method of choice for most researchers studying PPIs.^17^ However, IP-MS is limited in its throughput as it can only address the interactions of one protein at a time and each single experiment needs a rather large amount of sample input. Thus, IP-MS is often used to capture the PPIs of a few proteins of interest, while only a few groups have performed the tour de force experiments needed to map the interactomes of thousands of proteins, one IP at a time.^13–16^ As exemplified by the pioneering BioPlex dataset,^16^ which mapped the PPIs for thousands of proteins by IP-MS in two cell lines, extensively mapping PPIs even in a single cell line using IP-MS demands thousands of work hours and mass spectrometry runs. Even with such resources, investigating the temporal dynamics of PPIs at scale and at high resolution remains unattainable, despite its crucial relevance, especially in the understanding of physiological changes and disease states.

Here we present SPRINT-MS (SParse Reconstruction of INTeractions by Mass Spectrometry), an integrated experimental and computational method that leverages compressed sensing to drastically boost the throughput of IP-MS experiments (**Supplemental Figure 1b**). SPRINT-MS uses an algorithmically optimized pooling scheme whereby antibodies are mixed in different combinations and concentrations to form a set of pools. By conducting IP-MS using these antibody pools, each experiment measures a different linear combination of the lysate, thus implementing a compressed measurement (**Figure 1a**). The core strength of SPRINT-MS is that these pooled measurements, when combined with a novel algorithm that reconstructs PPIs, require substantially fewer IP-MS runs and less sample input compared to standard one-protein-at-a-time IP-MS experiments, increasing the throughput by about an order of magnitude. We demonstrate that the PPIs identified by SPRINT-MS are comparable to those identified by standard individual IP-MS experiments. We then further extend SPRINT-MS by demonstrating that pooling lysates containing different tagged proteins of interest and performing IP-MS on the lysate pools also identifies PPIs comparably to individual IP-MS experiments. Taken together, these results benchmark our novel approach for antibody- and lysate-based pooling and find that both methods produce results that are comparable to standard IP-MS, but at several fold higher throughput.

**Figure 1:**
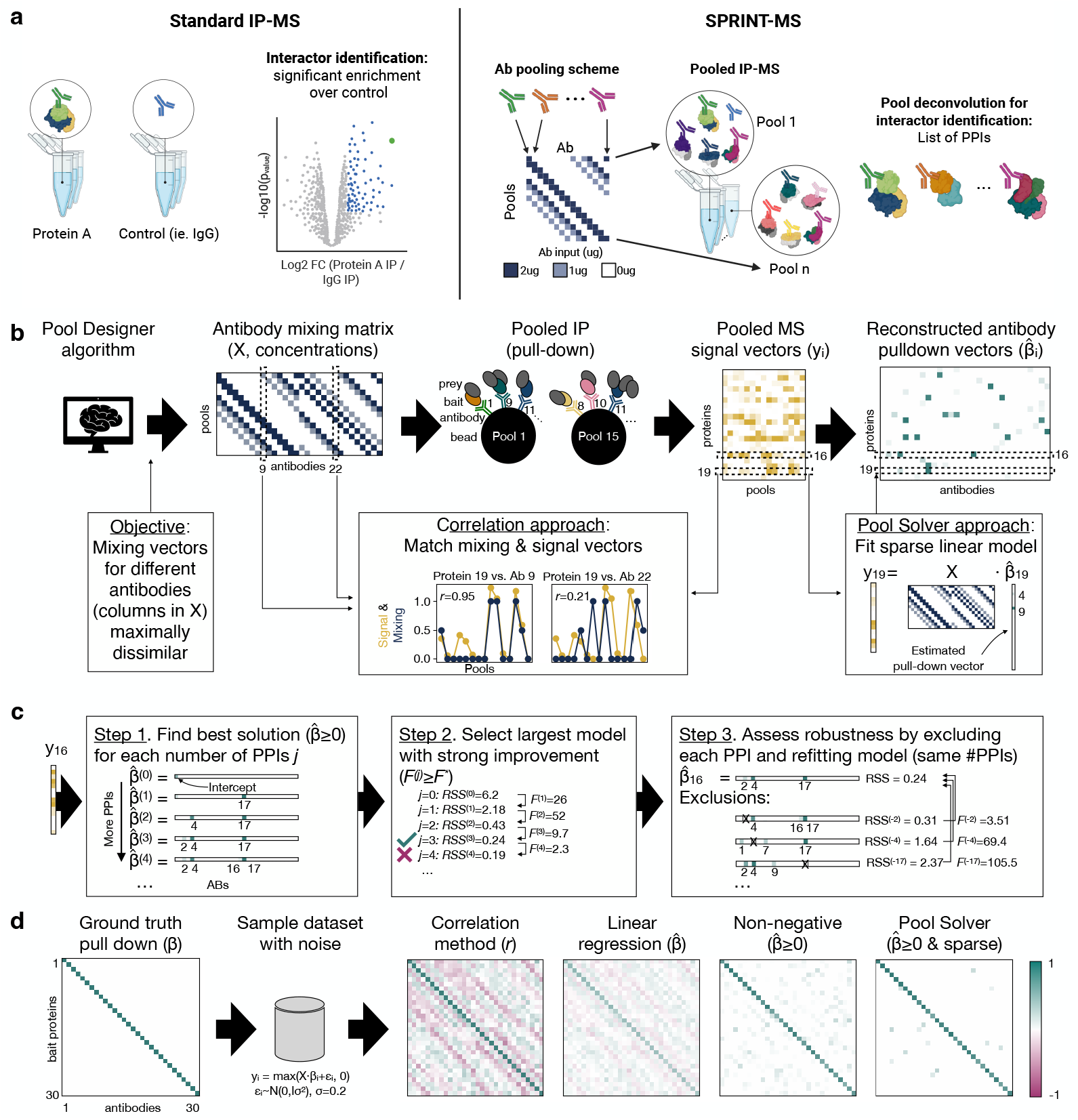
Overview of the SPRINT-MS platform and theoretical assessment using simulations. **(a)** Comparison between standard and pooled IP-MS. In standard IP-MS, single target IP-MS experiments as well as replicate negative control IP-MS experiments (typically 2 to 5 replicates each) are performed and statistically enriched interactors (by combined p-value and log_2_ enrichment value cutoffs) are identified. In contrast, in SPRINT-MS antibodies are combined into pools for pooled IP-MS. SPRINT-MS then uses the *Pool Solver* to recover the PPIs. (**b**) Schematic of the SPRINT-MS workflow: the *Pool Designer* (left box) generates a mixing matrix *X* specifying the concentration of each antibody in each pool. Following pooled IP and MS quantification, PPIs can be reconstructed from the pooled signal vector *y*_*i*_ (*i* indicating protein) using two approaches: by evaluating the correlation between the MS signal and the antibody concentrations (left box, showing a match between protein 18 and antibody 9 but not antibody 22), or by fitting a ‘pulldown’ vector 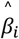 explaining the MS signal in terms of a sparse set of antibodies (right box, illustrating how antibody 9 alone explains the signal for protein 18). (**c**) Workflow whereby the *Pool Solver* reconstructs PPIs (exemplified by the reconstruction of protein 16 from panel (b)). The *Pool Solver* first regresses the MS signal onto the concentrations of different sets of antibodies, testing all combinations with *N*_*max*_ or fewer antibodies and constraining the pulldown coefficients (*β*) to be non-negative. For each number of antibodies *a*, the solution with the smallest residual sum of squares (RSS) is selected as 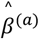 (horizontal bars in the left box, with the best single-antibody model containing antibody 17, etc.). Out of these solutions, the final pulldown vector 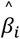 is selected to be the largest model with F-statistic (quantifying the decrease in RSS from adding one antibody to the model, normalized by the estimated noise) exceeding a threshold *F*^∗^ = 5 (check mark). To assess the robustness of each non-zero MS signal to antibody association in 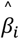, another F-statistic is finally computed by holding out the corresponding antibody from the analysis (crosses) and evaluating the RSS of the best model (of same size). (**d**) Simulation of bait proteins and comparison of signal reconstruction methods. In the simulation, each bait was assumed to only interact with its cognate antibody (diagonal band in left matrix plot, showing the stacked true pulldown vectors) and the pooled MS signal was sampled using additive normal noise and clipped at zero. The signal reconstruction (four right boxes) was performed using the correlation method, linear regression, non-negative least squares, and the final *Pool Solver*. The color bar indicates both the Pearson correlation coefficient and pulldown coefficient values.

## Results

### Overview of the Method

SPRINT-MS multiplexes traditional IP-MS by mixing combinations of antibodies into pools, conducting IP-MS using each antibody pool, and computationally reconstructing the set of antibodies that pulled down each protein (**Figure 1a**). The platform is more efficient compared to traditional IP-MS since it is possible to recover the set of interactions specific to an antibody with a smaller number of (pooled) measurements. The number of pools can be significantly smaller than the number of profiled antibodies, as long as each protein is only pulled down by a small fraction of those antibodies. Moreover, because each antibody is present in multiple pools, these pools act as *de facto* replicates, further increasing the efficiency compared to traditional IP-MS, for which 2-5 replicates per antibody and a negative control IP-MS with the same replicate number are needed.

A striking feature of PPI networks is that each protein typically interacts with a limited number of partners, and SPRINT-MS exploits this sparsity using ideas from compressed sensing. From this perspective, the set of interactors of a protein *i* can be represented using a sparse *pulldown vector β*_*i*_ that quantifies what antibodies enrich the protein. While such vectors were traditionally expensive to measure (one IP per target replicate), compressed sensing makes it possible to drastically reduce the number of required measurements. Put simply, compressed sensing maps the components of a sparse signal vector by first measuring several linear combinations of the components and then reconstructing the non-zero components computationally. Building on these ideas, SPRINT-MS efficiently constructs a pulldown vector *β*_*i*_ for each measured protein *i* (thus providing a list of the antibodies that enrich the protein) by considering linear combinations specified by the *mixing matrix X* (specifying how the antibodies are pooled) and measuring the resulting *pooled signal vector y*_*i*_ = *X* · *β*_*i*_ (quantifying the protein levels in each pool) by using IP-MS.

To understand how this compression works, consider a protein that is pulled down by one antibody. In an ideal experiment, an MS signal is only observed for the pools that included the antibody, and the signal is proportional to that antibody’s concentration in each pool. The antibody-protein pulldown can then be inferred by comparing the pooled MS signal to the concentration profile for each antibody and selecting the best match (line-scatter plots in **Figure 1b**). More formally, if *x*_*a*_ is the *mixing vector* specifying the concentrations of antibody *a* in each of the pools, then the pulldown of a protein by this antibody can be identified based on Pearson correlation coefficients *r*(*y*_*i*_, *x*_*a*_) being sufficiently large. While this method is simple and can be effective, it does not work for proteins that are pulled down by multiple antibodies, in which case the signal vector will be a linear combination of the corresponding antibody mixing vectors (and therefore correlate poorly with any one antibody mixing vector). Moreover, because all mixing vectors cannot be mutually orthogonal, the correlation coefficient will be non-zero for some antibodies that do not enrich the protein, but have mixing vectors similar to those of the target antibody.

To overcome these limitations, we developed a *Pool Solver* algorithm that reconstructs the pulldown vector by modeling the pooled signal and finding a sparse non-negative solution (see **Figure 1c** and Methods for details). For each putative protein-antibody pair (non-zero entry of *β*_*i*_), the algorithm also computes an F-statistic that quantifies the uniqueness and robustness of the solutions. Thus, the *Pool Solver* returns two values for each possible antibody-protein pair in the experiment: the ‘pulldown coefficient’ *β*, which essentially corresponds to the amount of measured protein signal that can be explained by that antibody, and the ‘F-statistic’, that indicates the uniqueness and robustness of that explanation.

### Theoretical Evaluation of the Feasibility of Pooled IP-MS - an *in-silico* stress test

While pooled IP-MS theoretically works for both large- and small-scale experiments (see Discussion), we expect that many users may be interested in the medium-scale use (targeting a few dozens of proteins of interest). We therefore designed a mixing scheme for multiplexing 30 antibodies using 15 pools, an experimental target protein scale that would normally necessitate over 100 individual IP-MS experiments. To take advantage of MS signal changes due to different antibody concentrations between pools and to guarantee multiple measurements for each antibody over all pools, each antibody was added three times at a “normal concentration” (standard individual IP-MS input) and twice at “half concentration”, meaning each antibody was present in 5 out of the 15 pools and each single pool contained 10 different antibodies. We next developed a *Pool Designer* algorithm that optimizes antibody-pool assignments in a way that minimizes the correlation between any pairs of mixing vectors while constraining the mixing matrix to have a cyclic structure that is practical to pipette (see Methods).

To establish the validity of the resulting mixing scheme (left matrix in **Figure 1b**), we next simulated pooled IP-MS experiments wherein each protein is pulled down by one or more antibodies. To test how the *Pool Solver* performs in the presence of noise, a normal error term with variance *σ*^2^ was added, and the signal was clipped at zero (MS intensities cannot be negative). These simulations allowed us to systematically test how well the ground-truth *β*_*i*_ used in the simulation can be reconstructed based on the noisy intensities *y*_*i*_.

In a first set of simulations, we considered the direct antibody target proteins (baits), each of which should be pulled down by its respective antibody. In terms of pulldown vectors, the baits should exhibit a strong signal only for their specific antibody (see diagonal band in ‘Ground truth’ in **Figure 1d**), and verifying if these specific antibody-bait interactions are recovered correctly provides a strong positive control for the platform (for the simulations we assumed that the antibody indeed efficiently enriches for its target protein/bait, which is not necessarily true for all of the antibodies used in experiments - see below). We first tested the simple Pearson correlation method and found that it recovered the diagonal band correctly, although weaker correlations were also observed for non-causal antibodies (‘Correlation Method’ in **Figure 1d**). We next found that while simply fitting a linear model using least squares did not recover the ground truth, adding the non-negativity constraint improved the reconstruction substantially (‘Linear regression’ vs. ‘Non-negative’ in **Figure 1d**). We finally applied the *Pool Solver* to the simulated data and found clear agreement with the ground truth (*Pool Solver* in **Figure 1d**).

We next simulated proteins pulled down by between one and five antibodies. In practical terms, this situation would arise if a protein interacts with multiple bait proteins (meaning it will be immunopurified by the corresponding antibodies), or if the bait of one antibody was also immunopurified by another antibody because it is a protein interactor of that bait, or if the same protein was targeted by different antibodies (e.g. to cross-validate results if antibody specificity might be an issue). As expected, simulating pulldown by more antibodies led to decreased correlation between the signal and the mixing vectors, meaning the correlation method became unlikely to identify the correct antibodies (**Supplemental Figure 2a**). While using a lenient detection threshold (in terms of *r* values) increased the sensitivity of the method, this also led to false detections in simulations where no true pulldown was included (**Supplemental Figure 2b**). In contrast, the *Pool Solver* with the cutoffs *β* ≥ 0.5 and *F* ≥ 5 reconstructed the ground truth with high probability (**Supplemental Figure 2a,b**), and these cutoffs were therefore used in the analysis below. To further assess the robustness, we simulated data with varied relative noise and found that the ground-truth is recovered when the ratio of signal-to-noise (*β*/*σ*) exceeded a value of about 3 (**Supplemental Figure 2c**). Altogether these results demonstrated the theoretical validity of the pooling scheme and the ability of the *Pool Solver* to correctly reconstruct PPIs even for proteins pulled down by several antibodies.

### SPRINT-MS generates experimental results that validate predicted bait protein behavior

We next turned to testing SPRINT-MS experimentally. As this experiment was intended as a proof-of-concept for our pooling method, we selected antibodies, that we had already tested in a previous IP-MS-based publication^18^ or that target stable complexes that we had identified across cell-lines by SEC-MS.^11^ In addition, we added as negative controls some antibodies that we had previously found to fail to enrich their expected baits - SMG1, DYNC1H1, and CPSF4 - and other ‘classical’ negative control antibodies, such as the isotype control IgG antibody and an antibody against V5. Finally, to test the ability of SPRINT-MS to simultaneously handle multiple antibodies against members of the same complex, we included pairs of antibodies, such as EXOSC2 and EXOSC10, that target different subunits of the same protein complex. After a calibration experiment to optimize the antibody concentrations (see Methods), the final pooling scheme (**Figure 2a**) used 2µg and 1µg of antibody as the ‘normal’ and ‘half’ concentrations, respectively, maintaining a constant total of 16µg of antibody in each pool. Using the mixing scheme output by the *Pool Designer* we performed 15 pooled IP-MS experiments targeting 30 baits on lysate from HEK293XT cells.

**Figure 2:**
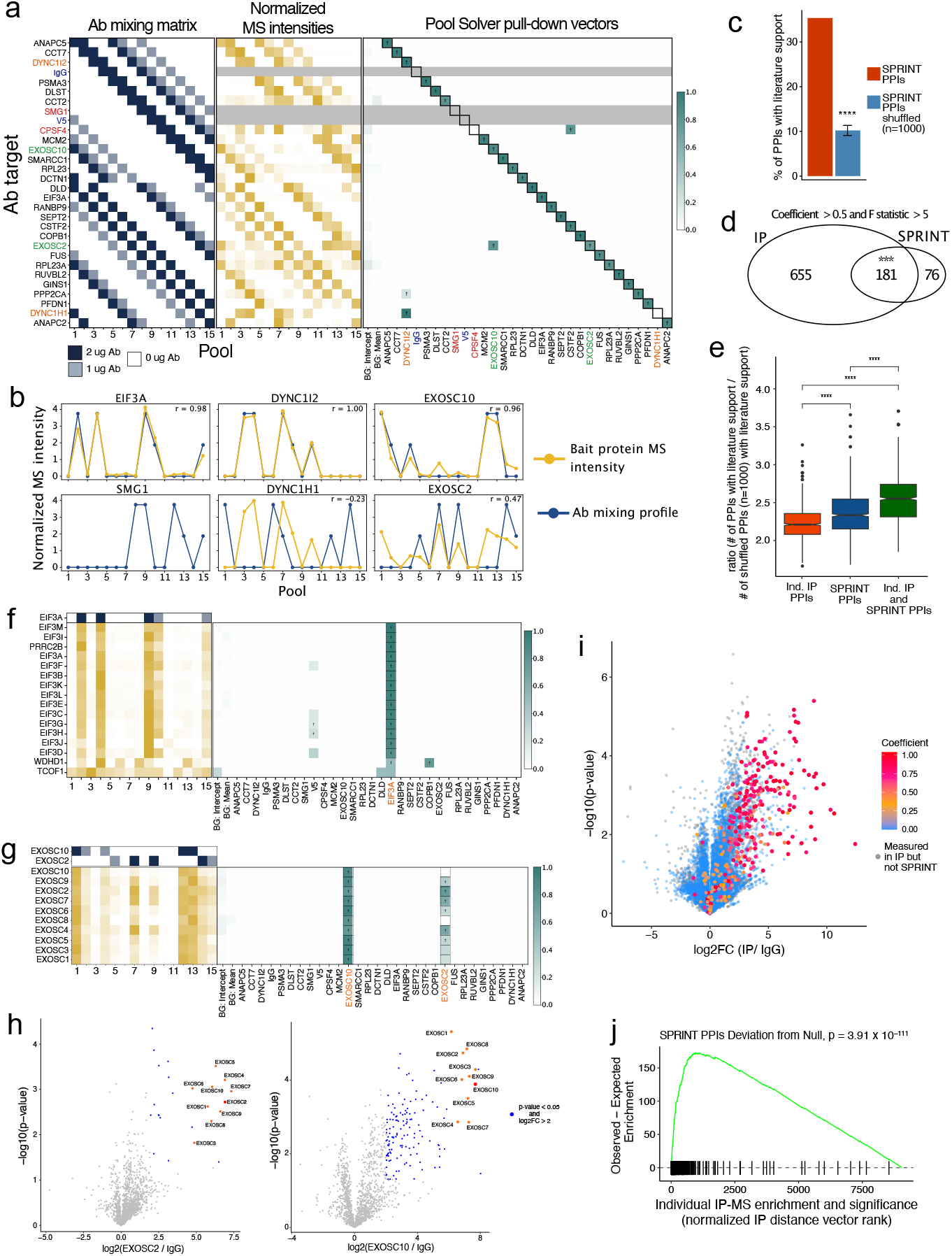
PPIs identified by SPRINT-MS are enriched for known PPIs and comparable to standard individual IP-MS experiments. (**a**) The antibody mixing scheme designed by the *Pool Designer* (left), normalized pooled MS intensities for the bait proteins (middle), and reconstructed pulldown vectors for the bait proteins (right). Antibodies with expected interactions are indicated by colored text. Daggers indicate F > 5, solid boxes indicate cognate antibody-bait pairs, and grey indicates antibodies without measured target protein MS signals. The first two columns of the pulldown heatmap reflect the portion of the MS signal attributed to background (intercept or mean pool intensity, see Methods) rather than antibody-specific enrichment. The scale of the pulldown vector coefficient values is shown (right). (**b**) Normalized MS intensities (gold) and antibody mixing profiles (blue) for select bait proteins, normalized to have a mean of 1. (**c**) Percent of literature supported PPIs compared with randomly shuffled bait-prey pairs from the PPI list for n=1000 tests, one-sample two-sided t-test p-value < 0.0001. (**d**) Venn diagram comparing the sets of PPIs identified by individual IPs and SPRINT-MS. Only shared antibodies were considered (***, p-value = 7.2 × 10^−128^, Fisher’s exact test). (**e**) Ratio of the percent of identified PPIs over the percent of randomly shuffled bait-prey pairs with literature support for individual IP PPIs, SPRINT-MS PPIs (for those baits overlapping with the individual IPs), and PPIs identified by both the individual IPs and SPRINT-MS (resampling test n=1000, one-sample two-sided t-test p-value < 0.001 for all comparisons). (**f**) Normalized MS intensities (left) and reconstructed pulldown vectors (right) for all proteins identified as interactors of EIF3A. The top heatmap (blue) indicates the anti-EIF3A antibody mixing vector. Daggers indicate F > 5, dashed boxes indicate literature supported interactions. Coefficient scale bar is shown on the right. (**g**) *Pool Solver* output (as in **f**) for all subunits of the RNA exosome complex. (**h**) Volcano plots of the individual standard IPs for EXOSC2 and EXOSC10. Members of the exosome complex are indicated in orange. (**i**) Volcano plot for all individual standard IPs with point colors indicating the *Pool Solver*-inferred pulldown coefficient scores (*ß*) (scale bar right) for each PPI also measured in the IPs. (**j**) Gene set enrichment analysis (GSEA) of the SPRINT-MS PPIs across a ranked list of the normalized distance vectors for all individual IP measured proteins. Lower ranks correspond to IPed proteins with larger vectors, indicating greater enrichment in the IPs (p-value = 3.9 × 10^−111^, K-S test). Ticks indicate SPRINT-MS PPIs and green line summarizes the distribution of the ratio of observed over expected SPRINT-MS PPIs.

After the appropriate normalization between the protein quantifications from the 15 pooled IP-MS runs (see Methods), we used the bait proteins (defined as the proteins that each antibody should bind as described by the vendor) as positive controls. First, we checked whether the signal vectors of the bait proteins matched the correct antibody mixing vectors (**Figure 2a**). Across most of the 30 antibodies, the bait protein signal overlapped well with the antibody mixing scheme **(Supplemental Figure 3a**). More specifically, we observed four general types of behavior which matched our expectations based on our a priori knowledge of the antibodies (**Figure 2b**): (**1**) for antibodies like anti-EIF3A that were previously shown to enrich the bait protein well and were the only antibody targeting the complex, as expected the bait protein signal overlapped cleanly with the antibody mixing scheme; (**2**) for antibodies like anti-SMG1, which we expected to fail to enrich for its bait, we indeed found that SMG1 was not even identified by the mass spectrometer; (**3**) for antibodies that targeted two members of the same complex, like the dynein complex antibodies - DYNC1|2 and DYNC1H1 - but where one antibody was not working well for IPs (we expected anti-DYNC1H1 to fail to enrich its bait); as expected, we found that both bait proteins were enriched by anti-DYNC1|2 and that their behavior followed the DYNC1|2 mixing scheme; (**4**) Finally, for antibodies targeting two members of the same complex, but where both enrich their respective bait, we expected both antibodies - for example anti-EXOSC2 and anti-EXOSC10 - to enrich their bait proteins and the rest of their protein complex; indeed, we found that while anti-EXOSC10 seemed to enrich the complex more strongly, the bait protein behavior is best explained when taking both antibody mixing schemes into account.

We next analyzed our pooled IP-MS data using the *Pool Solver* and found that the resulting pulldown vectors looked close to ideal for the targeted baits (**Figure 2c**). The few baits that were associated with several antibodies corresponded to expected antibody crosstalk, such as where two antibodies target different subunits of the same protein complex. As expected, our negative control antibodies showed no signal for their respective baits. Interestingly, the *Pool Solver* indicated that CPSF4 (for which the antibody failed as expected) is pulled down by another antibody in our pool, specifically the antibody targeting CSTF2, which is a known CPSF4 protein interactor. Altogether these results indicate that SPRINT-MS generates experimental data that behaves as expected based on our simulations.

### PPIs identified by SPRINT-MS are enriched for known PPIs and comparable to standard individual IP-MS experiments

Having established that SPRINT-MS reconstructs the bait proteins correctly, we next turned to the full dataset and asked whether the PPIs identified by SPRINT-MS are consistent with previously reported PPIs. We compared the SPRINT-MS PPIs to nine previously published datasets ^1,5,13,16,18–22^ and analyzed how the number of supporting datasets varied with the pulldown coefficient (*β*) and F-statistic reported by the *Pool Solver* (see Methods for detail). Consistent with the simulations above, this analysis indicated PPIs with *β* ≥ 0.5 and *F* ≥ 5 had the most robust literature support (**Supplemental Figure 3b**). Applying these cutoffs left us with 530 high-confidence PPIs for the 25 bait proteins that were successfully enriched. We next asked what percent of these PPIs were supported by the literature, finding that 35% of our PPIs had literature support (**Supplemental Figure 3c**, Supplemental Table 2), which is towards the top of the range reported by recent large-scale interactomes in human cells (BioPlex 3.0: 9.3% (Huttlin et al. 2021),^16^ BioPlex 2.0: 13% (Huttlin et al. 2017),^15^ Human Interactome: 16% (Hein et al. 2015),^20^ OpenCell: 38% (Cho et al. 2022)^13^ of all interactions have been previously reported) and also by our own previous IP-MS study (34%, Street et al. 2024)^18^. In addition, we showed that our PPIs did significantly better than expected by chance (**Figure 2d**; resampling test, n=1000, p-value < 1 × 10^−4^; see Methods for details). To directly test how our method compares to standard IP-MS experiments, we decided to perform our own standard IP-MS experiments using about a third of the antibodies (against 11 baits) used in the pooled experiment in triplicate in HEK293XT cells to enable direct comparison. Such individual IP-MS experiments are often regarded as the current gold standard in the field of PPIs. To select for the most enriched PPIs, we used cutoffs of a log_2_ fold change of IP over IgG control of 2 and a p-value cutoff of smaller than 0.05 to determine the individual IP PPIs, generating a list of 836 PPIs for the 11 bait proteins. When we subset the SPRINT-MS data to the PPIs of those 11 baits, we found that 181 of the 257 high-confidence SPRINT-MS PPIs were also identified in the individual IPs, significantly more than the 5.2 overlapping PPIs expected by chance (Odds Ratio: 34.8; p-value = 7.2 × 10^−128^, Fisher’s exact test) (**Figure 2d**). We also repeated the random PPI pair resampling test on the individual IP PPIs and the subset of SPRINT-MS-determined PPIs for the overlapping antibodies, finding that the percent of literature supported PPIs identified by both methods was very similar between the two approaches (greater than 2 times more than expected for random PPIs, resampling test, n=1000, p-value < 1 × 10^−4^; **Supplemental Figure 3d**). Perhaps unsurprisingly, PPIs identified by both SPRINT-MS and individual IP-MS showed the greatest enrichment compared to random, with more than 2.5 times as many PPIs having literature support (**Figure 2e**).

We next turned to specific antibodies to see how well the Pool Solver was able to identify the expected PPIs (Supplemental Table 3). We first looked at eIF3A, for which we have previously performed IP-MS experiments^18^ that demonstrated that the antibody cleanly enriches for eIF3A and the other components (eIF3B-M) of the thirteen-member eukaryotic translation initiation factor 3 (eIF3) complex. SPRINT-MS clearly identified all 13 components of the complex (**Figure 2f**). This high overlap between individual IP-MS and SPRINT-MS PPIs was also observed for antibodies that target subunits of the same complex, such as EXOSC2 and EXOSC10 - members of the 10-subunit RNA exosome. When we looked at all ten components of the exosome, we found that they were all identified as PPIs of EXOSC10 (**Figure 2g**). As expected, most of the subunits were also identified as PPIs of EXOSC2 (**Figure 2g**). We next compared these SPRINT-MS results with the individual IPs we performed against EXOSC10 and 2, which also identified the exosome components that we identified with SPRINT-MS as the most enriched (**Figure 2h**).

We next sought to compare the SPRINT-MS PPIs to our standard individual IPs more generally to better understand how the methods compare with each other. We therefore overlaid the pulldown coefficient (**Figure 2i**) and F-statistic (**Supplemental Figure 3e**) values over a combined volcano plot of all the individual IPs and observed a striking skew towards the most enriched individual IP PPIs, with higher coefficient (*β*) and F-statistic scores overlapping with the strongest and most significantly enriched IP proteins. To quantify this skew for our set of pool PPIs, we collapsed the log_2_FCs and p-values from the individual IPs into one normalized distance vector value (distance from the origin minus the difference in the legs (x and y, see Methods for details) and performed a gene set enrichment analysis (GSEA). This analysis also confirmed that the SPRINT-MS PPIs were highly significantly skewed towards the longest distance vectors, meaning the most enriched proteins in the individual IPs (**Figure 2j**, K-S test, p-value = 3.9 × 10^−111^).

Finally, we compared both the SPRINT-MS and individual IP PPIs to a completely orthogonal method - size exclusion chromatography-mass spectrometry (SEC-MS) - which identifies PPIs by co-elution rather than affinity purification. Using our previously-published data from HEK293XT cells,^11^ we asked what percent of PPIs had proteins with overlapping elution peak apices in the SEC data; 72% of PPIs identified in both SPRINT-MS and the individual IPs had overlapping apices, while 56% of SPRINT-MS and 48% of individual IP PPIs had overlapping apices, compared with only ~28% for negative control random PPI pairs of proteins in the SPRINT-MS or individual IPs (**Supplemental Figure 3g**).

Taken together, these data demonstrate that SPRINT-MS meets several important benchmarks when compared against the literature, individual standard IP-MS experiments, and an orthogonal method - SEC-MS. Across all metrics evaluated above, the SPRINT-MS method performed at least as well as the standard individual IP-MS approach.

### SPRINT-MS also works for lysate pooling

Building on the successful antibody pooling experiments above, we next turned to an alternative application of SPRINT-MS, namely pooling cell lysates rather than antibodies. This would have the advantage of also making SPRINT-MS accessible in settings and model organisms for which antibodies against endogenous proteins are not widely available, but for example, transgenic lines expressing tagged proteins of interest are. Theoretically, SPRINT-MS does not need to be adjusted for lysate pooling and the same *Pool Designer* and *Solver* algorithms can be used, with the only difference being that instead of pooling different antibodies according to the mixing matrix before performing pooled IP-MS from one cell lysate, here the cell lysates from different samples are pooled and IP-MS is done on those pooled lysates using a single antibody - in this case V5 (**Figure 3a**). For our proof-of-concept experiment for the lysate pooling, we used 30 *Saccharomyces cerevisiae* lysates, with a different protein V5-tagged in each lysate and mixed them into 15 pools using the same design we had used for the antibodies. After a pilot experiment to optimize single sample lysate input amounts (see Methods for details), we decided on 0.3 mg and 0.15 mg total protein as our “normal” and “half” lysate input amounts. The 30 lysates that we collected also included pairs of tagged bait proteins that are known members of the same complex (**Supplemental Figure 4a**, Supplemental Table 1). We followed the same experimental and analytical procedures as for the antibody pooling, and set out to evaluate the data across similar metrics.

**Figure 3:**
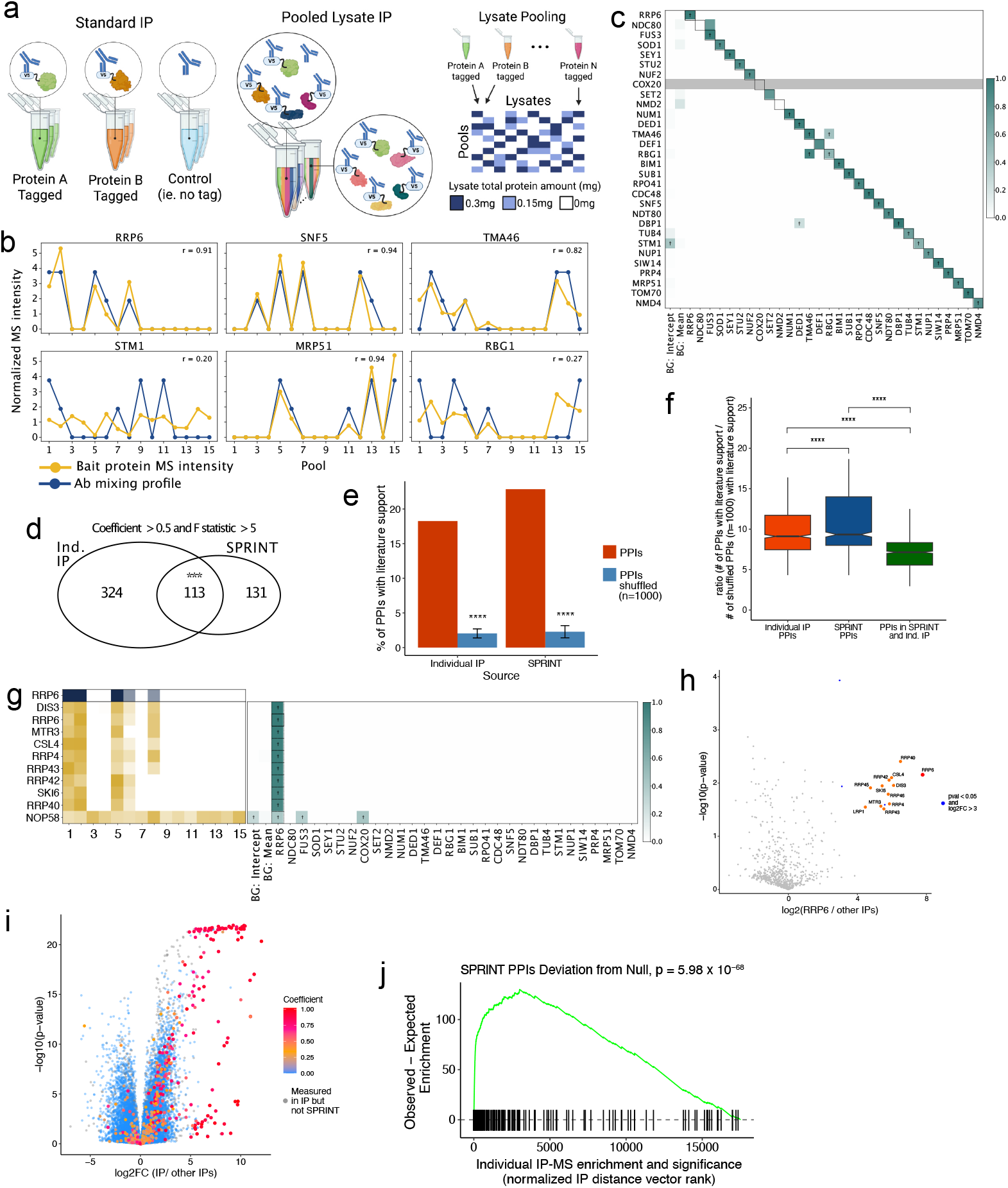
SPRINT-MS on pooled lysates performs comparably to individual IP-MS on the same lysate samples. (**a**) Overview of the lysate pooling approach. Lysates with V5-tagged proteins are mixed and IP-MS is performed with an anti-V5 antibody. (**b**) Examples of normalized bait protein signals across all 15 pools (gold). The amount of specific lysate added to each pool is indicated in blue. (**c**) Heatmap showing the reconstructed pulldown vectors (coefficient scores) for the bait proteins, shown as in Figure 2a. (**d**) Overlap of the individual standard IP PPIs with the SPRINT-MS PPIs (***, p-value = 1.4 × 10^−148^, Fisher’s exact test). (**e**) Percent of literature supported PPIs compared with randomly shuffled bait-prey pairs from the PPI list for n=1000 tests, one-sample two-sided t-test p-value < 0.001. (**f**) Ratio of the percent of identified PPIs over the percent of randomly shuffled bait-prey pairs with literature support for individual IP PPIs, SPRINT-MS PPIs, and PPIs identified in both the individual and pooled IPs (resampling test n=1000, one-sample two-sided t-test p-value < 0.0001 for all comparisons). (**g**) Output of the *Pool Solver* for all proteins identified as interactors of the bait protein RRP6, shown as in Figure 2f. (**h**) Volcano plots of the individual standard IPs for RRP6. Members of the exosome complex are indicated in orange. (**i**) Volcano plot for all individual standard IPs with point colors indicating the Pool Solver coefficient scores *ß* (scale bar right) for each PPI also measured in the IPs. (**j**) Gene set enrichment analysis (GSEA) of the SPRINT-MS PPIs across a ranked list of the normalized distance vectors for all individual IP measured proteins. Lower ranks correspond to IPed proteins with larger vectors, indicating greater enrichment in the IPs (p-value = 6 × 10^−68^, K-S test). Ticks indicate SPRINT-MS PPIs and green line summarizes the distribution of the ratio of observed over expected SPRINT-MS PPIs.

To assess the quality of the pooled signal, we first compared the signal vectors for the tagged bait proteins to the corresponding lysate mixing vectors and found that the two overlapped well (**Figure 3b, Supplemental Figure 4b**). As with the pooled antibody experiment, we observed different patterns of bait-mixing overlap: (**1**) IPs of baits such as RRP6, SNF5, and MRP51, which are all the sole baits in their respective complexes, followed the lysate mixing scheme cleanly; (**2**) Failed IPs - STM1 was not enriched above background signal and was considered a failed IP; (**3**) IPs against baits in the same complex - e.g. the protein signals of TMA46 and RBP1, which form a conserved heterodimeric GTPase involved in translation,^23^ overlap with the combined mixing patterns of both lysates (**Figure 3b**). The Pool Solver was able to disentangle the expected lysate to tagged bait interactions with high specificity, following closely the ideal lysate-bait pulldown vector pattern, and showing cross-reactivity only for bait proteins that are members of the same protein complex (**Figure 3c**). The expected cross-reacting bait pairs were translation regulators TMA46 and RBP1 and initiation factor paralogs DBP1 and DED1,^24^ while STM1 and NMD2 were explained by background terms in the model, and COX20, which was not measured by the mass spectrometer, was absent. Interestingly, the reconstruction of NDC80 indicated, with low-confidence (*F* = 4.84), pulldown by FUS3. This misattribution was likely due to a secondary contribution to the NDC80 signal from its known interactor NUF2,^25^ that by chance allowed the total signal to be approximated by the FUS3 mixing vector.

As there are fewer high-quality PPI literature sources for yeast compared to human, and we had not previously performed large-scale IP-MS experiments in yeast, we decided to perform individual standard IP-MS experiments on all the lysates used in the pooled experiment to allow for robust comparison between methods. After performing the individual IP-MS experiments for all 30 baits, we ended up with a total of 437 PPIs across all tagged bait proteins (see Methods for details). We next generated a list of high-confidence SPRINT-MS PPIs (using *F* ≥ 5 and *β* ≥ 0.5 as above) and compared these to the PPIs identified by the individual IPs. Of the 244 SPRINT-MS PPIs, 113 were shared with the individual IPs, yielding a highly significant overlap compared to the 3.2 PPIs overlapping expected by chance (**Figure 3d**, Odds Ratio: 35.3, p = 1.4 × 10^−148^, Fisher’s exact test). We next asked what percent of these SPRINT-MS and individual IP PPIs were supported by the literature; 22.5% of SPRINT-MS PPIs have literature support while 18.3% of the individual IP PPIs were supported (**Figure 3e, Supplemental Figure 4c**, Supplemental Table 4). We again performed the resampling test in which we shuffled the PPI protein pairs to see how much literature support would be expected for random interactions, finding that the SPRINT-MS PPIs had ~11 times and the individual IP PPIs ~9 times more support in the literature than expected by chance (**Figure 3e,f**, resampling test, n=1000, p-value < 1 × 10^−3^). This strong recapitulation of known PPIs by SPRINT-MS can also be seen across the full pool data, with almost all previously reported interactions having very high pulldown coefficient (*β*) and F-statistic values (**Supplemental Figure 4d**).

We next assessed the *Pool Solver* at the level of an individual bait protein (see Supplemental Table 5 for all bait-specific results). RRP6, a component of the yeast RNA exosome,^26^ was one of the bait proteins that agreed well with the mixing scheme and was the sole bait in the exosome complex. Every PPI with a high coefficient (*β*) value is an exosome subunit (**Figure 3g**), and the individual standard IPs for the RRP6 lysate, also identified the exosome components as the most enriched (**Figure 3h**).

We finally compared the SPRINT-MS and individual IP PPIs more generally by overlaying the pulldown coefficients (*β*) (**Figure 3i**) and F-statistics (**Supplemental Figure 4e**) from the *Pool Solver* over a combined volcano plot of all the individual IPs. Similar to the trend observed for the pooled antibodies, this analysis revealed that the most enriched individual IP PPIs had the largest pulldown coefficients (*β*) and F-statistics scores. To quantify this enrichment, we again performed a gene set enrichment analysis (GSEA; as above), which confirmed that the SPRINT-MS PPIs were highly enriched amongst the most enriched proteins in the individual IPs (K-S test, p-value = 6 × 10^−68^) (**Figure 3j)**.

Taken together, these results demonstrate that SPRINT-MS works well for both antibody and lysate pooling, increasing the study of PPIs in both cases by about an order of magnitude.

## Discussion

Here we present and benchmark SPRINT-MS, an integrated computational and experimental method of pooling IP-MS experiments, that produces results comparable to gold standard individual IP-MS experiments with an order of magnitude increase in throughput. This substantial increase in throughput extends beyond merely reducing the measurement time in LC-MS/MS; it also proportionally diminishes the required sample input and reduces the hands-on labor in the laboratory. Consequently, this heightened throughput will render the medium-to large-scale investigation of PPIs in rare or challenging samples more attainable. Furthermore, SPRINT-MS imposes no specific sample constraints, rendering it suitable for application to the wide array of samples where individual IP-MS experiments can be conducted.

### Limitations of SPRINT-MS

However, over the course of our proof-of-concept experiments for the antibody and lysate pooling we encountered several points which are important to consider and currently puts some limitations on the applicability of SPRINT-MS.

Firstly, our current method is designed to use all antibodies at the same high and low input amounts. In our preliminary experiments we tested multiple amounts for each antibody and chose the two that seemed to best produce differentiable target protein MS intensities. Our method, however, seems to be quite sensitive to differences in antibody affinity meaning that the high and low antibody amounts might have to be adjusted for certain antibodies. The *Pool Designer* can be modified to take these differences into account.

Secondly, and related to the prior antibody differences, solving the reconstruction problem appears to be more challenging if multiple antibodies that differ in their binding affinities target the same complex. This challenge is illustrated by the IPs of the exosome proteins EXOSC2 and EXOSC10; when performed individually using the standard IP-MS approach, both antibodies enrich their baits and the full exosome complex with high efficiency (**Figure 2h**). However, in the pooled approach, the EXOSC10 antibody captured both baits and the other complex members more strongly than the EXOSC2 antibody (**Figure 2g**). Although SPRINT-MS picked up nearly all the known exosome interactors for both antibodies, we envision that taking differences in antibody strength into account would improve our analysis platform.

Finally, we found that baits in large complexes, such as the cytoplasmic ribosome (**Supplemental Table 3**), did not recapitulate known PPIs as cleanly as baits for smaller complexes. Generally, it is not always easy to enrich for the ribosome by IP-MS since it is a very large complex that forms many interactions with nuclear and cytoplasmic proteins.^18^ In fact, we found previously that RNase treatment - either to partially digest the ribosome itself, or to digest larger RNP complexes - improved our ability to comprehensively IP the ribosome.^18^ Since the ribosome is one of the most abundant complexes in the cell, we expect it to be both highly abundant in the background and to form interactions with many proteins. As the *Pool Solver* leverages the sparsity of PPI networks to reconstruct the pull down vector (meaning it looks for enrichment due to a specific antibody, or set of specific antibodies, compared to all other IPs in the full experiment rather than compared to a control IP such as IgG or V5), highly abundant and connected complexes will be more difficult to capture than by standard individual IP-MS. Interestingly, the mitochondrial ribosome in the lysate pooling was identified well, perhaps because its compartment specific localization reduces its number of interactions and background abundance. This is an avenue for improvement of the method, but can also be seen as a partial advantage as many promiscuous interactions, which sometimes make the interpretations of individual IP-MS results difficult,^27^ will be automatically filtered out.

## Conclusion

Despite these current limitations, SPRINT-MS provides for robust IP-MS PPI capture at an order of magnitude higher throughput and can be directly applied to many current biological questions. For example, SPRINT-MS could be applied to projects like BioPlex,^16^ where a cDNA library was used to transiently express one tagged protein at a time to do thousands of individual IP-MS experiments. However, using the pooling strategy we introduce here, varying pools of cDNA plasmids could be transfected and then IP-MS could be done on these different pools and our Pool Solver could be used to deconvolve the PPI networks. This could increase the throughput of a monumental endeavor such as BioPlex several fold. Indeed, while we decided to use a mid-scale approach (e.g., 30 antibodies) for our proof-of-concept experiments, SPRINT-MS can be scaled up to higher levels of throughput. In our computational modeling of SPRINT-MS, we tested multiplexing 100 antibodies across 20 pools and found that at least for the modeling, this scale worked as well as the 30 antibodies by 15 pools method we applied here (data not shown). Testing this larger scale is an interesting avenue for future work that will further demonstrate the broad applicability and flexibility of our pooling method.

## Figures and Figure Legends

**Supplemental Figure 1:**
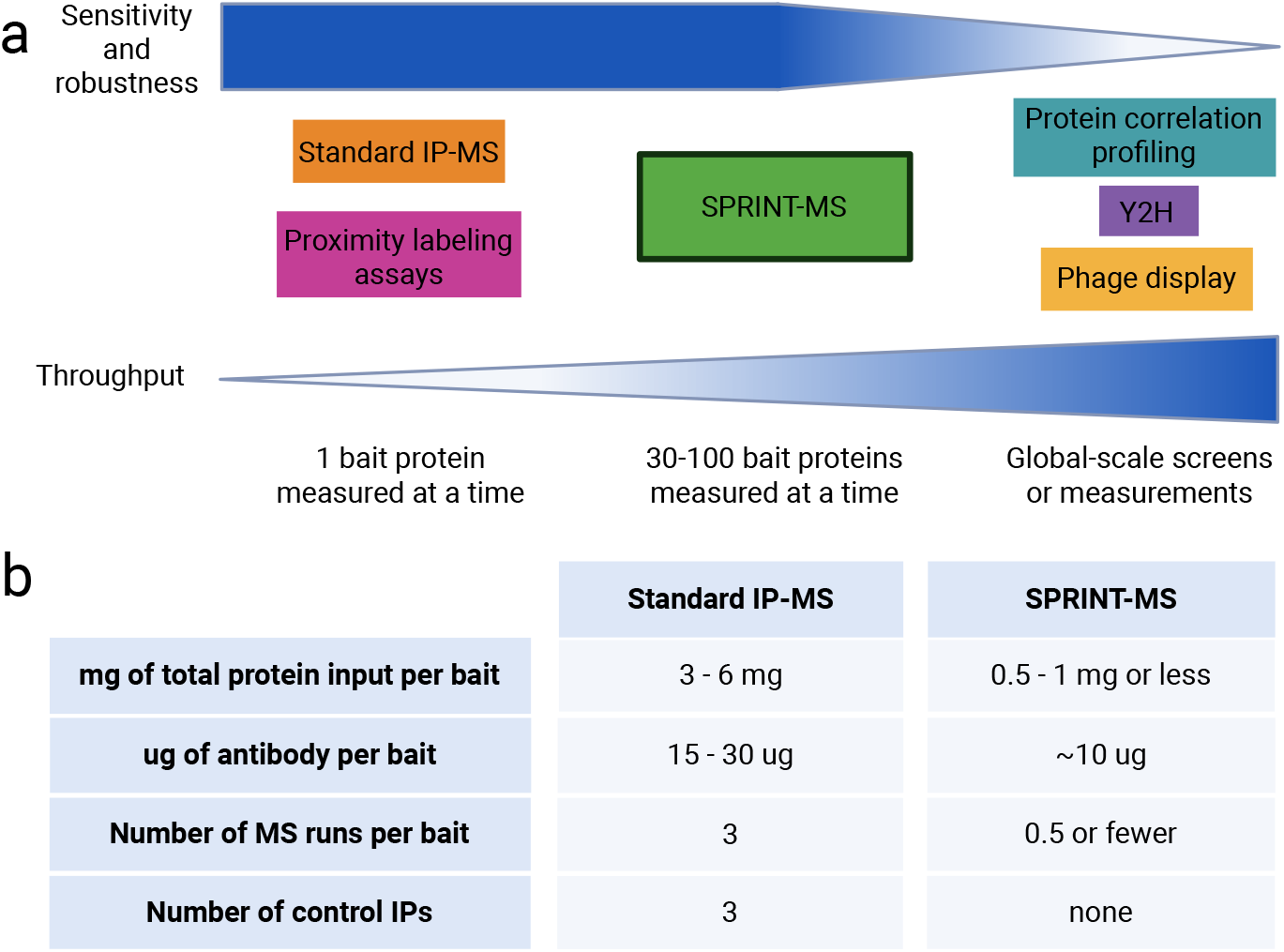
Use case for SPRINT-MS. (**a**) The sensitivity and robustness (on a qualitative scale) versus the throughput for current protein-protein interaction capture methods. (**b**) The per-target protein input and experimental requirements in standard IP-MS versus SPRINT-MS.

**Supplemental Figure 2:**
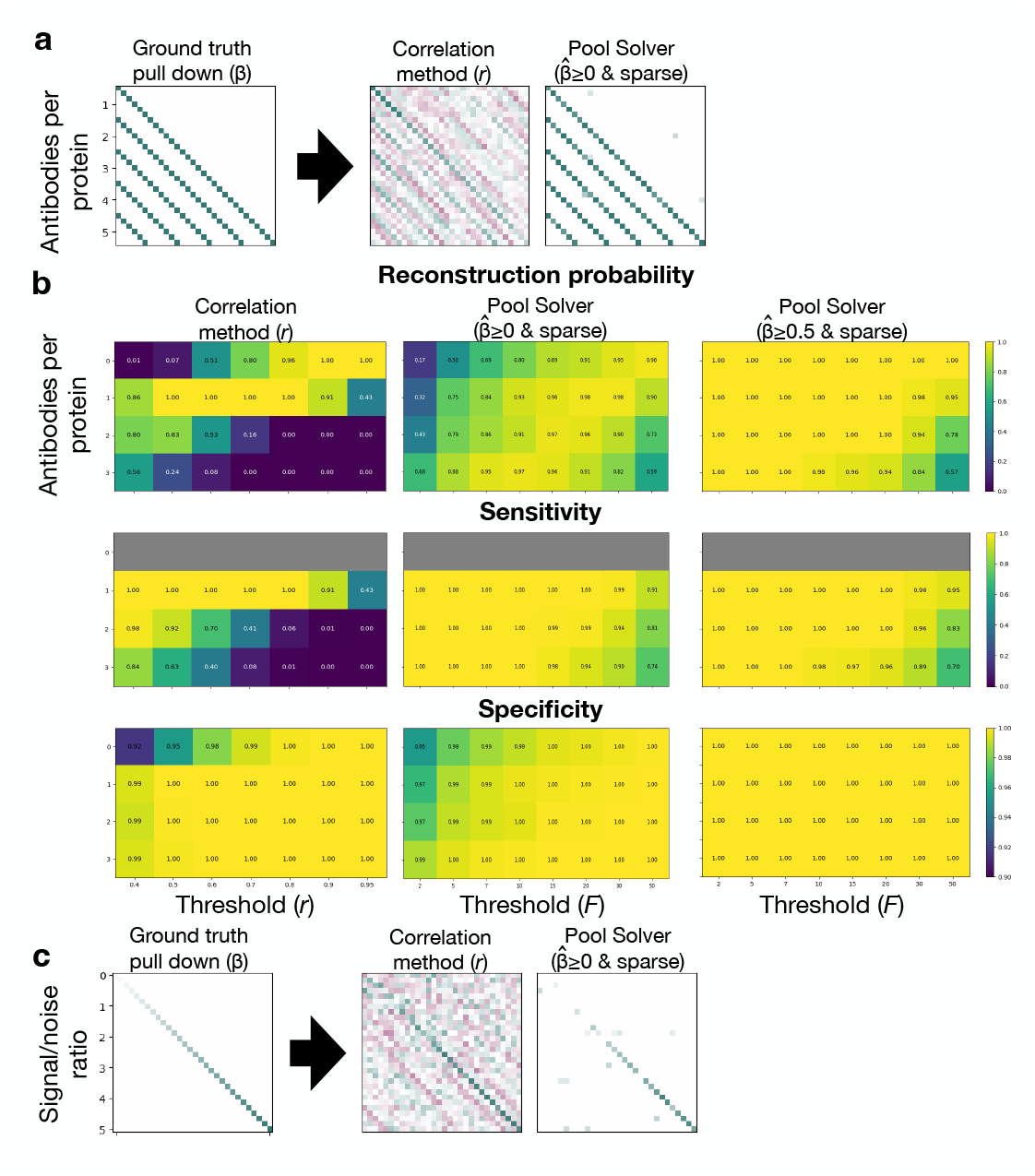
Simulations and signal reconstruction for proteins interacting with multiple baits and with varying signal-to-noise ratio. (**a**) Ground truth and reconstructed interactions for proteins pulled down by between one and five antibodies, displayed as in Figure 1d. A diagonal ground truth pattern was used for ease of comparison. (**b**) Simulation of proteins pulled down by between zero and three randomly selected antibodies. For each number of antibodies, 100 simulations were performed, the signal vectors were normalized (see Methods), and the sets of antibodies were reconstructed using the correlation method (left heatmaps, columns corresponding to applying different r thresholds) and the Pool Solver (middle heatmaps, using F*=5 in step 2 of Figure 1c, columns corresponding to applying different F thresholds on the final PPIs), and using the Pool Solver with the additional detection threshold *β* > 0.5 (right heat maps). In each simulation, the discrete set of reconstructed antibodies was compared to the set in the ground truth and the reconstruction probability was estimated as the fraction simulations with a perfect match (top heatmaps, right scale bar). In addition, the sensitivity (middle heatmaps, right scale bar) and specificity (bottom heatmaps, right scale bar) were computed to further characterize errors. (**c**) Simulations of bait proteins with varying signal-to-noise ratio, displayed as in Figure 1d.

**Supplemental Figure 3:**
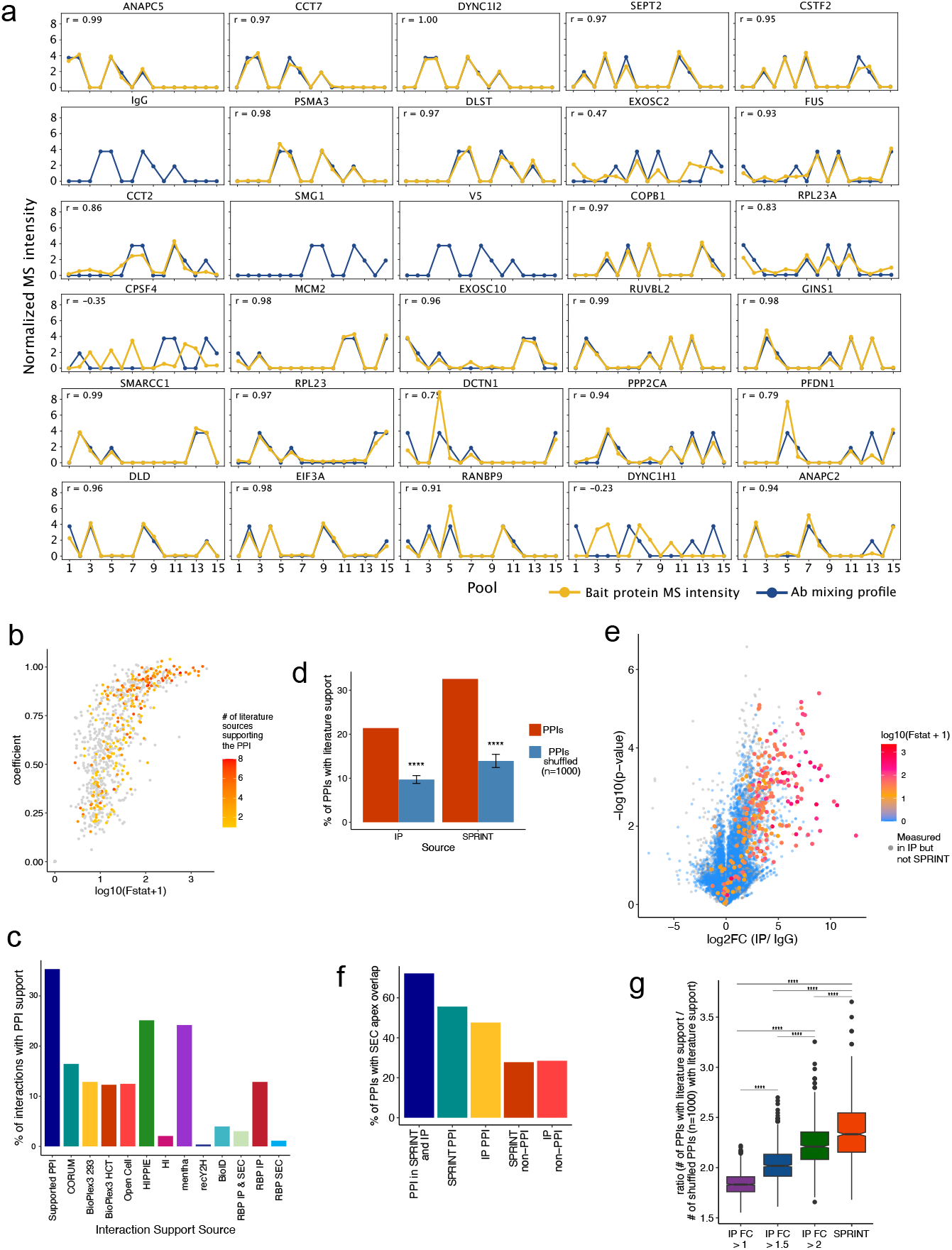
PPIs identified by SPRINT-MS are enriched for known PPIs and comparable to standard individual IP-MS experiments. **(a)** Normalized bait protein signals across all 15 pools (gold) for all antibodies (Ab) in the pooled experiment. The amount of antibody added to each pool is indicated in navy. **(b)** *Pool Solver* coefficient scores plotted against F-statistic values for all PPIs identified by the *Pool Solver*. Points are colored by the number of literature support sources for each PPI (see scalebar). **(c)** Percent of SPRINT-MS PPIs with literature support separated by the source of the support. **(d)** Percent of literature supported PPIs for the individual standard IPs and the overlapping SPRINT-MS IP baits compared with randomly shuffled bait-prey pairs from the respective PPI lists for n=1000 tests, one-sample two-sided t-test p-value < 0.001. **(e)** Volcano plot for all individual standard IPs with point colors indicating the *Pool Solver* F-statistic values (right scale bar) for each PPI also measured in the IPs. **(f)** Percent of PPIs with overlapping SEC-MS peak apices. **(g)** Ratio of the percent of identified PPIs over the percent of randomly shuffled bait-prey pairs with literature support for individual IP PPIs at different log_2_FC cutoffs and SPRINT-MS PPIs for those baits overlapping with the individual IPs (resampling test n=1000, one-sample two-sided t-test p-value < 0.0001 for all comparisons).

**Supplemental Figure 4:**
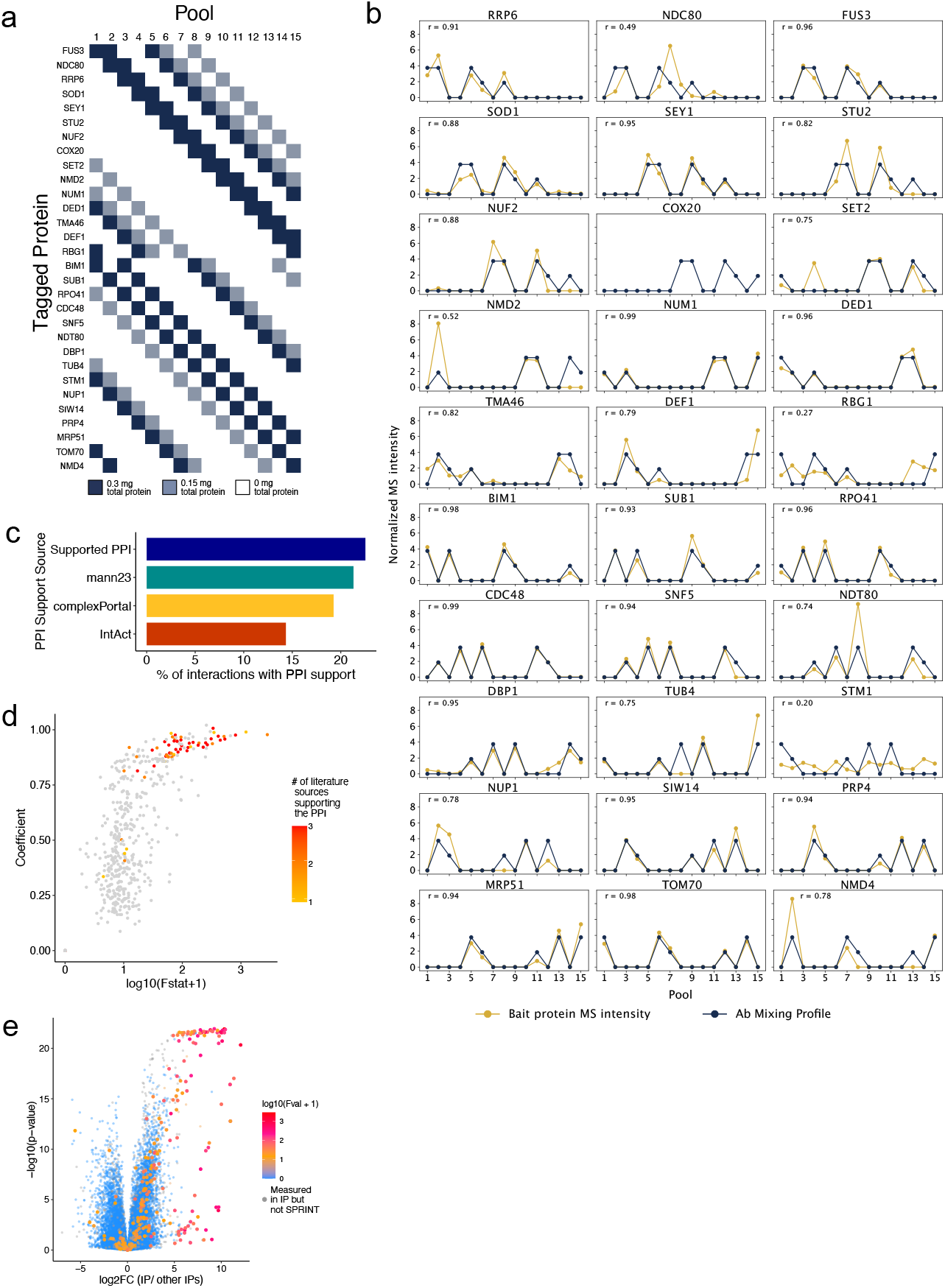
SPRINT-MS on pooled lysates is comparable to standard individual IP-MS. (**a**) The lysate mixing scheme designed by the *Pool Designer*. (**b**) Normalized tagged bait protein signals across all 15 pools (gold) for all tagged protein lysates in the pooled experiment. The amount of lysate added to each pool is indicated in navy. (**c**) Percent of SPRINT-MS PPIs with literature support separated by the source of the support. (**d**) Pulldown coefficients plotted against F-statistic values for all PPIs identified by the *Pool Solver*. Points are colored by the number of literature support sources for each PPI (see scalebar). (**e**) Volcano plot for all individual standard IPs with point colors indicating the *Pool Solver* F-statistic values (scale bar right) for each PPI also measured in the individual IPs.

## Supplemental Tables

Supplemental Table 1: Antibody and lysate information.

Supplemental Table 2: Full dataset for the Pooled Antibody experiment. The table combines the output matrix of the Pool Solver with the enrichment values (log_2_FC and p-values) for all proteins measured in the individual IP experiments and the literature support information.

Supplemental Table 3: The Pool Solver generated PPI heatmaps for all antibody baits.

Supplemental Table 4: Full dataset for the Pooled Lysate experiment. The table combines the output matrix of the Pool Solver with the enrichment values (log_2_FC and p-values) for all proteins measured in the individual IP experiments and the literature support information.

Supplemental Table 5: The Pool Solver generated PPI heatmaps for all lysate baits.

## Code and data availability

SPRINT Pool Designer and Pool Slover available here: https://github.com/RubeGroup/SPRINT-MS All data is uploaded to MassIVE: ftp://massive-ftp.ucsd.edu/v11/MSV000099960/

## Acknowledgements

This study was supported by grants from the NSF (Award # 2516741 to T.R. and M.J.) and the NIH (R01 AG071869 to M.J. and G.B.). G.W.Y. is supported by National Institutes of Health (NIH) grants R01 HG004659 and U24 HG009889.

## Author contributions

M.J. and T.R. conceived the study. L.S., K.R., G.W.Y., and M.J. designed initial experiments. L.S. and K.R. performed initial experiments. L.S., T.R., and M.J., designed subsequent experiments. L.S., E.G., and G.B. performed subsequent experiments. M.M. and T.R. built and optimized the computational platform. L.S., M.M., and T.R. analyzed the results. L.S., M.M., T.R., and M.J. wrote the manuscript. T.R. and M.J. acquired funding and supervised the study.

## Materials and Methods

### The utility of sparse signal reconstruction for deconvolving pooled signals

The central observation underlying SPRINT-MS is that pooling of antibodies (or lysates with different tagged proteins) makes it possible to map PPIs with drastically increased throughput. Analogous pooling strategies were devised long ago to control infectious disease, most famously to identify syphilitic conscripts during World War 2.^28^ To understand how this pooling strategy works, consider the extreme case where exactly one individual in a group of 1,000 has syphilis and where blood has been drawn from everyone. This individual is most efficiently identified using a binary tree search^29^: first form two groups of 500 and test pooled blood from one group. After the test reveals the group containing the sick individual, this group is divided in half and pooled testing is repeated. In the end, only 10 grouped tests are needed to identify the sick individual, much less than the 1,000 tests needed for individual screening. While more tests are needed if multiple individuals are sick, the speedup can still be substantial if the prevalence is low.^30,31^

While the above example assumes that the test is binary and that a single sick individual is sufficient to cause a positive pooled test, pooling can also boost throughput for quantitative linear measurements. Compressed sensing aims to estimate a high-dimensional signal of interest (vector *β*) based on a limited number of measurements. Rather than measuring all signal components directly, compressed sensing measures a limited number of linear combinations of the signal (*y* = *X* ⋅ *β*, where *X* is the sensing matrix defining the linear combinations) and reconstructs the signal computationally. Foundational work showed that efficient reconstruction is possible even when the dimension of the signal greatly exceeds the number of measurements so long as the signal is sufficiently sparse (that is, *β* mostly contains zeros) and the sensing matrix has favorable properties.^32–34^ Several studies have applied pooled testing strategies and compressed sensing for biological measurement, for example proposing to identify rare alleles using pooled DNA sequencing^35^ and COVID-19 cases using pooled RT-PCR,^36^ and boosting the throughput of RNA-seq and Perturb-seq^37,38^. In 2009, one study increased the throughput of a yeast two-hybrid experiment by pooling plasmids encoding different interactors.^39^ However, while compressed sensing has been highly successful in boosting throughput across multiple domains, it has not yet been leveraged to map PPIs using IP-MS.

### PPI Reconstruction

#### Setup and linear model

To model the system, we assumed that in an ideal pooled IP-MS experiment the MS signal for a given protein *i* in each pool receives linear contributions from each antibody in that pool, and that this contribution is proportional to the concentration of the antibody. The MS intensity vector *y*_*i*_ for protein *i* (with vector index running over pools) can then be modeled using a system of linear equations:

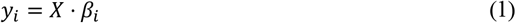

Here the *mixing matrix X* (which is generated by the *Pool Designer*, see below) encodes the concentration of each antibody (columns) in each pool (rows), and the *pulldown vector β*_*i*_ quantifies the association between the MS intensities for protein *i* and the concentration of the antibodies (entries). Note that for equation (1) to be satisfied, the MS intensities for the different pools need to be normalized to the same scale (see discussion below). Because we expect non-zero *β*_*i*_ entries to arise through antibody-dependent pulldown mediated by PPIs, the goal of pooled IP-MS is to identify such entries with high sensitivity and specificity. In order to prevent non-specific contributions to contaminate the pulldown vector, two ‘background’ columns were added to *X*: the first column contains ones and implements a standard intercept term, and the second column contains the mean signal in each pool (after normalization, see below) in order to absorb any proteome-wide bias. Because the number of unknowns in this model equals the number of antibodies plus two, and because the number of equations (one per pool) is less than the number of antibodies, the system is underdetermined, meaning it does not have a unique solution and the true value of *β*_*i*_ be reconstructed. However, a unique solution can exist if it is known that most (but not which) *β*_*i*_ entries are zero, and if the mixing matrix is well designed. The goal of the algorithms below is to accomplish this.

#### Pool Solver Algorithm

The *Pool Solver* constructs an estimated pulldown vector 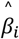 by solving equation 1. Because this is an underdetermined linear system, it does not have a unique solution and system and simply solving it does not recover the ground truth (Figure 1d). However, foundational theoretical results established that a unique solution can exist if it is known *a priori* to be sparse. ^48^ In the case of pooled IP-MS, this requirement is satisfied if only a few baits pull down each protein, meaning most entries in 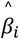 are zero. To find such sparse solutions, we first constrained the entries in *β*_*i*_ to be non-negative (reasoning that antibodies cause, rather than obstruct, pulldown). Such non-negativity constraints are known to promote sparsity^40^ and this alone improved the signal reconstruction (Figure 1d).

To explicitly impose sparsity on the solutions, the *Pool Solver* uses a modified version of best subset selection. In a first step (c.f. Figure 1c), the algorithm loops over the sparsity level *j* = 0, . . ., *N*_*max*_ and constructs a series of candidate solutions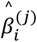, each with exactly *j* non-zero antibody associations. *N* = 5 was used for the synthetic-data analysis (Figure 1), but this was decreased to *N*_*max*_ = 3 for the pilot dataset analyses (Figures 2 and 3) since using a larger value only marginally increased the number of discoveries. For each number *j*, the *Pool Solver* tests all 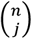 subsets of the *n* antibodies. For each such subset, it tests all 4 combinations of the two ‘background’ columns in *X* (see above), meaning these columns are not counted for the purpose of sparsity and therefore are preferentially used to explain the MS signal. Specifically, the *Pool Solver* constructs a reduced mixing matrix *X*′ by extracting the relevant background/antibody columns (for each of the above combinations) and computes the ordinary least squares estimate (*X*′^*T*^*X*′)^−1^*X*′^*T*^*y*_*i*_. It then rejects all combinations with one or more negative pulldown coefficients and selects combination with the smallest residual sum of squares (RSS) to be the candidate solution 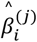 (with associated error 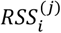).

Having created a series of candidate solutions, the *Pool Solver* next (step 2 in Figure 1c) assess each candidate using a F-statistic that compares the RSS reduction facilitated by including an additional antibody to the estimated variance of the random error:

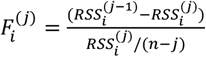

Note that this statistic is not guaranteed to follow a F-distribution, both because of the non-negativity constraints and because it was computed after variable selection. The *Pool Solver* then selects a final 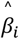 by finding the largest value of *j* for which 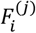 exceeds a pre-set threshold *F*^∗^ (set to 5 by default).

In the final step (step 3 in Figure 1c), the *Pool Solver* assesses the uniqueness and robustness of the solution.

Specifically, given that an antibody *a* has a positive entry in the solution 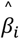, the *Pool Solver* excludes this antibody from the analysis, repeats the first step above to find the best alternative solution 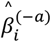 with the same sparsity, and computes the error 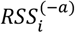 of this reduced mode. It then evaluates a modified F-statistic that compares the error full model (*RSS*_*i*_) to that of the reduced model:

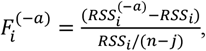

Intuitively, this statistic would be zero if the MS signal can be explained without invoking antibody *a* (meaning the reconstruction is not unique and 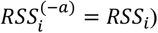, and of order one if the error reduction facilitated by allowing antibody *a* is similar to the estimated noise. The *Pool Solver* repeats this analysis by separately ablating each antibody with a non-zero association. The final output of the method is one matrix containing the pulldown vector for all proteins and one containing the matched F-values.

### Pool Designer Algorithm

The goal of this algorithm is to design the mixing matrix *X*_*ka*_ specifying how much of antibody *a* is added to pool *k*. In designing this matrix, we aimed to accomplish several objectives; (i) the mixing scheme should demonstrate increased efficiency by profiling 2*n* antibodies using *n* pools, meaning the matrix dimension is *n* × 2*n*; (ii) each antibody should be included in *n*_*H*_ pools at high concentration and *n*_*L*_ pools at lower concentrations; (iii) all pools should have the same set of antibody concentrations; (iv) all antibodies should have similar mixing patterns to simplify pipetting; and (v) the columns should be maximally dissimilar in order to make it easy to distinguish the pulldown profiles of the different antibodies;

To accomplish these objectives, we considered mixing matrices composed of two blocks (denoted *A* and *B*), each with dimension *n* × *n*; each block is specified by a generating vector (denoted *v*_*A*_ and *v*_*B*_) and the *j*’th column of the block is defined to be the (*j −* 1)’th right cyclic permutation of the generating vector. The generating vectors were constrained to contain *n*_*H*_ and *n*_*L*_ high and lower entries, respectively, with the remaining entries being zero. It is straightforward to show that this design satisfies objectives (i)-(iii) above and the repeating pattern makes pipetting more straightforward and errors less likely. Altogether these constraints reduce the design problem to the combinatorial problem of finding the optimal placement of the high and low values in *v*_*A*_ and *v*_*B*_.

Because the number of possible placements 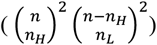 grows quickly with *n*, we devised a greedy two-step algorithm that explores a reduced search space: The first step optimizes the placement of the *n*_*H*_ high values in each of *v*_*A*_ and *v*_*B*_. Specifically, it evaluates a function *L*(*X*) for all 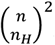possible placements of the high values (each placement implicitly defining a mixing matrix *X*) and records all placements tied to be optimal for the second step. To penalize designs where two antibodies have similar mixing patterns, the loss function was chosen to be

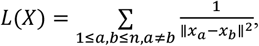

where *x*_*a*_ is the *a*’th column of *X*. The search can be made more efficient by noticing that the loss function is invariant under relabeling of the pools. In particular, applying cyclic permutations to the generator vectors (which does not violate the constraints above) does not change the loss function, meaning there are several degenerate generating vectors. By shifting one of the high values to the first position of the generator vectors, and by shifting the largest gap between high values to the end of the vector, we can eliminate these degenerate vectors. The second step of the algorithm considers all possible placements of the *n*_*L*_ lower values in each of the optimal generator vectors recorded in the first step and selects the placement with the smallest value of the loss function as the final mixing matrix.

### Data normalization, filtering, and fitting models

#### Filtering

Proteins with low signal quality were filtered out using two criteria: First, because a unique reconstruction is impossible if the signal vector contains two or fewer non-zero entries, such proteins were removed from the analysis. Second, proteins with MS signal (quantified as the mean of the top three values) below a threshold (2 × 10^7^ for IP pooling and 1 × 10^7^ lysate pooling) were also removed.

#### Protein normalization

Even after applying the above filtering, the MS intensities varied by four orders of magnitude when comparing different proteins. Importantly, such variations can be entirely unrelated to PPIs and reflect variations in abundance in the initial lysate and differences in detection efficiency. Because the pulldown vector is proportional to the MS signal, it cannot be compared across proteins unless they are normalized to a common scale. To accomplish this, we reasoned that the top three values for each protein likely correspond to the signal from one antibody at the higher concentration (because each antibody is included in three pools at this concentration). We therefore estimated high-concentration signal by computing the mean of these values and normalized the signal vector by this value, thus ensuring that the mean of the top 3 values equals 1.

We note that this normalization in principle could lead to an underestimate of *β*_*i*_ for proteins that are pulled down by multiple antibodies. For example, if two antibodies pull down a protein with equal efficiency, and if the corresponding mixing vectors also have maximum overlap (meaning both occur with the higher concentration in two of the pools), then the final pulldown will be underestimated by 40%. However, repeating the synthetic data analysis with and without the normalization did not change the results appreciably, likely because the normalization cancels out in the F-statistics.

#### Pool normalization

The central assumption underlying our linear model (equation 1) is that adding the same amount of an antibody to two pools should result in the same MS signal intensity. However, because the pools may pull down different amounts of total protein, and because the MS intensities only quantify the relative protein abundance within each pool, the intensities may not be directly comparable. For example, if an antibody that pulls down a large amount of protein is added to a pool, then the relative intensities for proteins pulled down by other antibodies in the pool would decrease. While this potential normalization issue did not appear to have a major impact on our datasets (the raw MS intensities tracked the antibody concentrations relatively well), proper normalization is an important assumption, and we therefore developed a normalization algorithm.

The goal of this algorithm is to learn a global vector δ that contains one multiplicative normalization factor per pool. Using vector notation, pool normalization corresponds to the transformation *y*_*i*_ → *δ* ∘ *y*_*i*_, where o denotes element-wise vector multiplication. Because incorrect normalization should make the model (equation 1) fit the data poorly, we reasoned that *δ* can be estimated by finding the value that minimizes the RSS. However, because repeatedly running the *Pool Solver* on the full dataset is computationally expensive, the algorithm instead focuses on the subset of proteins for which a single antibody explains the signal well. Specifically, it uses the correlation method with threshold *r*^2^ > 0.8 identify putative antibody-protein pairs, keeping at most 20 proteins per antibody to avoid an imbalanced dataset. For each of these pairs, the algorithm next performs linear regression and computes the residual. Specifically, after creating a reduced matrix 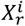 containing the relevant columns of the mixing matrix, the residual is

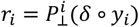

where 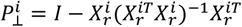 is the residual maker matrix. The total squared error for all the putative pairs is then

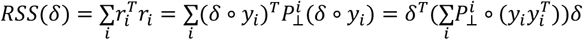

We next minimized this while constraining the normalization vector to have 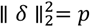 (meaning the mean squared normalization factor entry is one), in which case *δ* is simply given by the eigenvector of 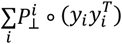 with the smallest eigenvalue. Because applying the pool normalization *δ* to the dataset could impact both the protein normalization (which identifies the top three pools for each protein) and the first step of the pool normalization (which identifies putative protein-antibody pairs), the normalization steps were repeated until convergence (change in *δ* smaller than 10^*-5*^). Finally note that while *δ* is adjusted to increase the agreement between the mixing matrix and the MS intensities, the number of parameters (15) is much smaller than the number of intensity values (3,810 for the IP), meaning the normalization procedure is unlikely to artificially inflate the agreement.

### Cell Culture

HEK293XT (Takara Bio Lenti-X 293T, #632180) cells were cultured in DMEM containing L-glutamine and sodium pyruvate supplemented with 10% Fetal Bovine Serum, Penicillin (100 U/mL), and Streptomycin (100 μg/mL). Cells were grown to 90-100% confluency before passage and were harvested at passages 12-18.

For harvesting, ~90% confluent HEK293XT cells in a 10 cm plate were gently washed in cold 1X PBS and dissociated from the plate with a cell scraper. Cells were then pelleted in cold PBS at 500 g for 5 minutes. Cells were then resuspended in 1 ml of ice-cold PBS and transferred to 1.5 ml Protein Lo-Bind tubes to be pelleted again and flash-frozen in liquid nitrogen and stored at −80 °C.

### Yeast culture

#### Mitotic Culture

Cells (for strains indicated as mitotic in Supplemental Table 1) were inoculated and grown 24h in YEPD (yeast extract peptone dextrose) media and diluted to OD600 0.05 in YEPD. They were grown with shaking at 30C and harvested at OD600 0.6 (4.5 h of growth).

#### Meiotic/BYTA Culture

Cells (for strains indicated as meiotic or spores in Supplemental Table 1) were inoculated and grown 24h in YEPD (yeast extract peptone dextrose) media and diluted to OD600 0.4 in BYTA (buffered yeast extract tryptone acetate). These were then grown overnight and either harvested or washed with sporulation media (0.5% KAc pH7) and resuspended in sporulation media at OD1.9. Cells were either sporulated with shaking at 30C for 4.5h or 24h (for the spore sample).

#### Harvesting and extract preparation

Cells were collected by filtration and rapid freezing in lysis buffer (150 mM NaCl, 50 mM Tris pH 7.5, 1% NP-40, 5% glycerol, 2mM MgCl_2_, 1X PhosSTOP (Sigma), 1X Ultra Complete protease inhibitor (Roche), 0.64 µg/ml AEBSF, and 1 µg/ml Pepstatin A). Lysis was performed by mixermiling in liquid nitrogen, powder was thawed, and subjected to a low-speed spin (3K RCF 5 min at 4C) and supernatant was flash frozen. The total protein concentration of each lysate was measured using a BCA assay.

### Immunopurifications

#### HEK293XT lysis

Cell pellets were thawed on ice and lysed in 400 µl of lysis buffer (150 mM NaCl, 50 mM Tris pH 7.5, 1% IGPAL-CA-630, 5% glycerol, and protease inhibitors), and incubated on ice for 20 minutes. Lysates were centrifuged at 4°C for 10 minutes at 14,000 g. The total protein concentration of each lysate was measured using a BCA assay.

#### Pooled Ab IP set-up

To select “normal” and “half” Ab input amounts, we tested three µg amounts (0.5, 1, and 2.5µg) for each Ab, multiplexed into 4 pools in total that we used to perform IP-MS. We then compared the enrichment of the Ab bait proteins against control V5 Ab IP-MS data to determine which µg amounts captured a linear difference between the low and high concentrations with good enrichment of the bait intensity above V5 control and background signal. While the input Ab amounts can be adjusted for each Ab, we found that generally all Ab selected for the experiment behaved well between the range of 1 to 2.5µg, meaning that we saw bait enrichment, and stronger enrichment with the higher antibody amount. In order to keep the total µg of Ab per pool consistent, we chose to use standard 1µg for low and 2µg for high input for all Ab in the experiment (Supplemental Table 1, CST Abs added volumetrically, 2uL for high and 1uL for low).

Antibodies (Supplemental Table 1) were mixed into pools following the mixing scheme designed by the Pool Designer (Figure 2a). 100 µl of Dynabeads Protein G (Invitrogen:01200616) magnetic beads per pool were washed 3 times in 5 ml lysis buffer and then mixed with the cell lysate (~1 mg total protein per IP). Bead-lysate mix was then added to each Ab pool and incubated overnight at 4°C on a rotator.

#### Individual standard IP set-up (Ab)

50 µl of Dynabeads Protein G (Invitrogen:01200616) magnetic beads were washed 3 times in 1 ml lysis buffer and then combined with the cell lysate (0.5 mg total protein) and 2.5 µg of Ab and incubated overnight at 4°C on a rotator. Three replicates were done per Ab.

#### Pooled lysates IP set-up

Lysates (Supplemental Table 1) were mixed into pools following the mixing scheme designed by the Pool Designer (Supplemental Fig 4A), resulting in 2.4mg total protein in 480µl per pool. 100 µl of Dynabeads Protein G (Invitrogen:01200616) magnetic beads per pool were washed 3 times in 5 ml lysis buffer and then added to the lysate pools along with 10 µg of V5 monoclonal Ab (Invitrogen 46-0705). The IPs were incubated overnight at 4°C on a rotator.

#### Individual standard IP set-up (lysate)

50 µl of Dynabeads Protein G (Invitrogen:01200616) magnetic beads were washed 3 times in 1 ml lysis buffer and then combined with the cell lysate (0.5 mg total protein) and 2.5 µg of V5 Ab (Invitrogen 46-0705) and incubated overnight at 4°C on a rotator. Two replicates were done per cell lysate.

#### General IP protocol

The following day, samples were placed on a magnetic bead separator, the supernatant was removed, and samples were washed 2 times with wash buffer (150 mM NaCl, 50 mM Tris pH 7.5, 5% glycerol) containing 0.05% IGEPAL CA-630 and 2 times with wash buffer without IGEPAL. Beads were then incubated in 80 µl of on-bead buffer (2 M urea, 50 mM Tris pH 7.5, 1 mM DTT, and 5 μg/mL Trypsin (Promega:487603)) for 1 hour at 25°C on a shaker (1000 rpm). After one hour, beads were placed on a magnetic bead separator and 80 µl of supernatant was transferred to a new tube. The beads were then washed twice with 60 μL of HPLC buffer (2 M urea and 50 mM Tris pH 7.5). The supernatant from each wash was combined with the on-bead digest for a total of 200 µl per sample. Samples were then spun at 5000 g, transferred to a new tube, and stored at −80°C.

#### Ab selection

15 RBP Abs we had previously used and validated^18^ and that produced a range of PPI interactome sizes were selected. Additionally, we identified 15 well-described non-RBP complexes from our published SEC-MS data in HEK293XT cells^11^ and ordered immunopurification (IP)-grade Abs for a subunit of each of those complexes. After initial testing of the unvalidated Abs we replaced any that had failed to enrich their expected bait protein. The V5 monoclonal Ab used for the lysate pooling had been previously validated to enrich the tagged proteins across most of the 30 strains used.

### LC-MS/MS

#### IP sample preparation

100 μL of partially digested proteins were used per IP sample. Disulfide bonds were reduced with 5 mM dithiothreitol (DTT) for 45 minutes at 600 rpm and 25°C. Cysteines were subsequently alkylated with 10 mM iodoacetamide (IAA) for 45 minutes in the dark at 600 rpm and 25°C. Samples were then further digested by adding 0.5 μg sequencing grade modified trypsin (Promega) for 16 hours at 600 rpm and 25°C. After digestion, samples were acidified with a final concentration of 1% formic acid. Tryptic peptides were desalted on C18 StageTips according to (Rappsilber, Mann, and Ishihama 2007)^41^ dried in a vacuum concentrator and reconstituted in 15 μL of 3% acetonitrile / 0.2% formic acid for mass spectrometry.

#### LC-MS/MS

5 μL of total peptides were analyzed on a Waters M-Class UPLC using a 15cm Ion-Optics column (1.7um, C18, 75um × 15cm) or 25cm Ion-Optics column (1.7um, C18, 75um × 25cm) for the pooled IP samples and individual IP samples, respectively, coupled to a benchtop ThermoFisher Scientific Orbitrap Q Exactive HF mass spectrometer. Peptides were separated at a flow rate of 400 nL/min with a 90 min gradient, including sample loading and column equilibration times. Data was acquired in data-dependent mode. MS1 spectra were measured with a resolution of 120,000, an AGC target of 3e^6^ and a mass range from 300 to 1800 m/z. MS2 spectra were measured with a resolution of 15,000, an AGC target of 1e^5^ and a mass range from 200 to 2000 m/z. MS2 isolation windows of 1.6 m/z were measured with a normalized collision energy of 25.

### IP-MS Analysis

#### Searches

Raw data were searched with MaxQuant v2.0.3.0 (Cox, J., Mann, M. 2008)^42^ using a UniProt database (Homo sapiens, UP000005640)^43^ under default LFQ quantification settings. Protein group data were exported for subsequent analysis.

#### Normalization and determination of interacting proteins

Normalization and subsequent analysis were performed in R v4.5.0. Contaminants, immunoglobulin proteins and proteins only identified by site or reverse were removed. For the pooled samples, the raw data was then run through the *Pool Solver* with F=10 and the F statistic = 5. Interacting proteins were identified as those that had coefficient values > 0.5 and F statistic values > 5. For the individual IPs, missing values were imputed randomly from the bottom of the signal distribution of the LFQ intensity values and then the LFQ intensity values were log2 transformed. Proteins with a mean MS/MS count value for every IP bait below 3 were removed from subsequent analysis. Interacting proteins were identified as those that for Ab: passed a log2FC sample IP over IgG IP cutoff of 2 and a p-value of 0.05, or lysate: passed a log2FC sample IP over all other IPs (measured at the same time) cutoff of 3 and a p-value of 0.05.

### Resampling test for the PPIs

We firstly calculate the percentage of PPI pairs (bait-prey) that are reported by published PPI datasets (Ab: CORUM,^44^ BioPlex,^16^ OpenCell,^13^ Mentha,^19^ HuRI,^20^ HIPPIE,^21^ rec-Y2H,^1^ BioID,^5^ RBP interactome^18^; Lysate: IntAct,^45^ Mann 23,^46^ compexPortal^47^) out of all our PPI pairs, calling this *Fexp*. We next randomly shuffled the bait-prey pairs in our PPI lists so that the baits were matched with random preys and calculated the percent of random PPIs supported by the published PPI datasets, calling this *F*. We repeated the random sampling steps 1000 times and got the resampled distribution {*F*_*i*_}, where i ranges from 1 to 1000. The one-sample t-test p-value is calculated as the proportion of the resampled distribution where *F*_*i*_ was greater than or equal to *F*_*exp*_.

### Standard IP normalized distance vector

To collapse the log2FC and −log10(p-value) values for all proteins measured by IP-MS onto a single value that, when ranked, generally captures the separation between proteins that would have passed the dual log2FC and p-value cutoffs used to determine enriched interacting proteins. To do this we generated a new metric which is essentially a normalized distance vector (*n*_*dy*_) for each protein from the origin of the volcano plot, where

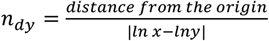

### SPRINT-MS implementation considerations

The application of cutoffs to identify significantly enriched proteins is an inherent, if subjective, part of standard IP-MS analysis. Perhaps unsurprisingly, increasing cutoffs increases the proportion of interactions with literature support (Supplemental Figure 3h), which is often used as a benchmark to assess the reliability of the identified PPIs. In previous publications we have applied cutoffs individually for every antibody as the combination of different protein abundances as well as antibody affinities and mass spectrometry measurement biases combine to generate, at times, large differences in measured enrichment values between IP-MS experiments. While we applied additional cutoffs in the analysis of our SPRINT-MS data, using all PPIs identified by the *Pool Solver* with a coefficient value greater than zero already achieved most of the specific enrichment of high confidence interactors that we achieved through the use of relatively stringent cutoffs in the individual standard IPs (Figures 2j, 3i, Supplemental Figures 3e, 4e). This indicates that depending on individual experimental considerations, SPRINT-MS could possibly be implemented without the need for cutoffs.

