## Supplemental Table 3 for "SPRINT-MS: A high-throughput platform for identifying protein-protein interactions using pooled IP-MS and sparse signal recovery"

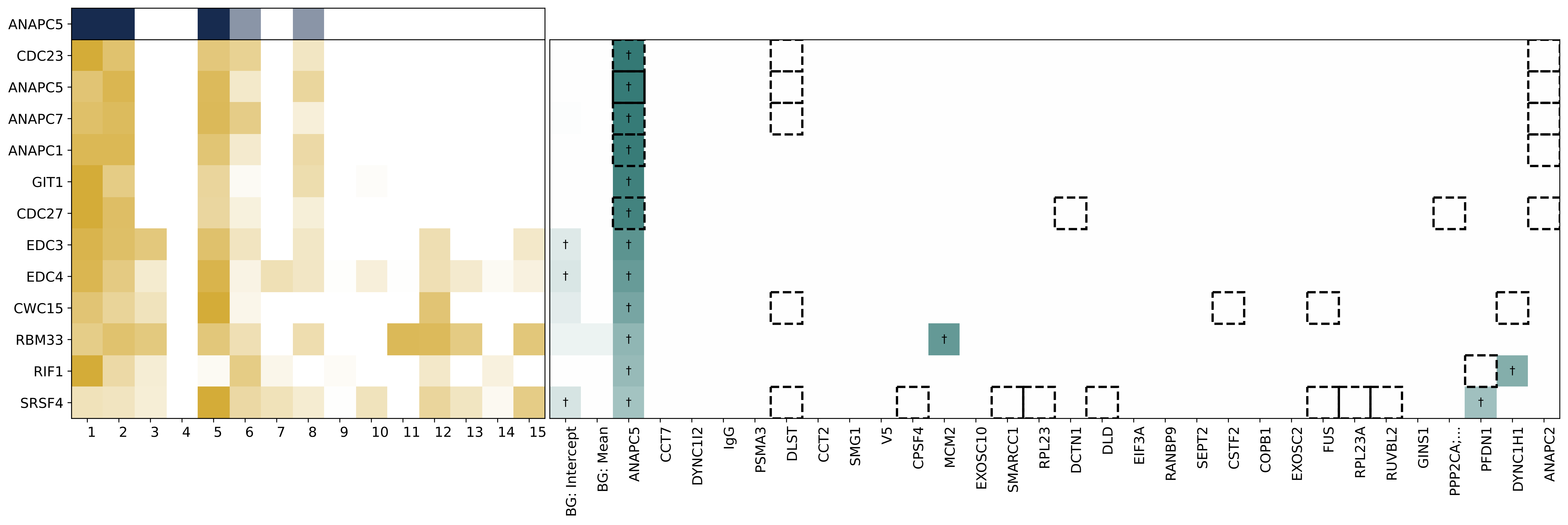

ANAPC5
CDC23
ANAPC5
ANAPC7
ANAPC1
GIT1
CDC27
EDC3
EDC4
CWC15
RBM33
RIF1
SRSF4
1 2 3 4 5 6 7 8 9 10 11 12 13 14 15
BG: Intercept
BG: Mean
ANAPC5
CCT7
DYNC1I2
IgG
PSMA3
DLST
CCT2
SMG1
V5
CPSF4
MCM2
EXOSC10
SMARCC1
RPL23
DCTN1
DLD
EIF3A
RANBP9
SEPT2
CSTF2
COPB1
EXOSC2
FUS
RPL23A
RUVBL2
GINS1
PPP2CA;...
PFDN1
DYNC1H1
ANAPC2

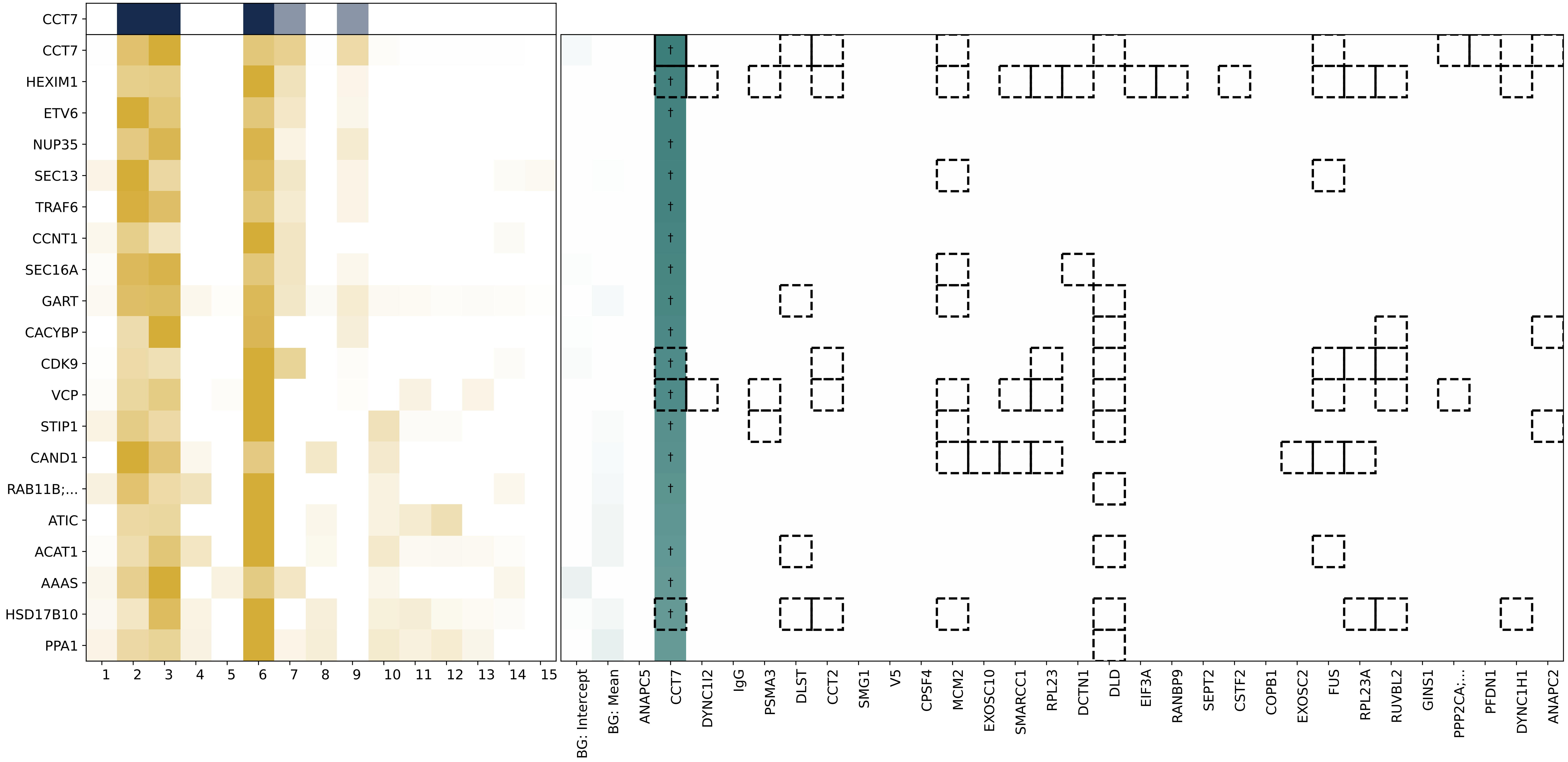

CCT7
CCT7
HEXIM1
ETV6
NUP35
SEC13
TRAF6
CCNT1
SEC16A
GART
CACYBP
CDK9
VCP
STIP1
CAND1
RAB11B;...
ATIC
ACAT1
AAAS
HSD17B10
PPA1
BG: Intercept
BG: Mean
ANAPC5
CCT7
DYNC1I2
IgG
PSMA3
DLST
CCT2
SMG1
V5
CPSF4
MCM2
EXOSC10
SMARCC1
RPL23
DCTN1
DLD
EIF3A
RANBP9
SEPT2
CSTF2
COPB1
EXOSC2
FUS
RPL23A
RUVBL2
GINS1
PPP2CA;...
PFDN1
DYNC1H1
ANAPC2

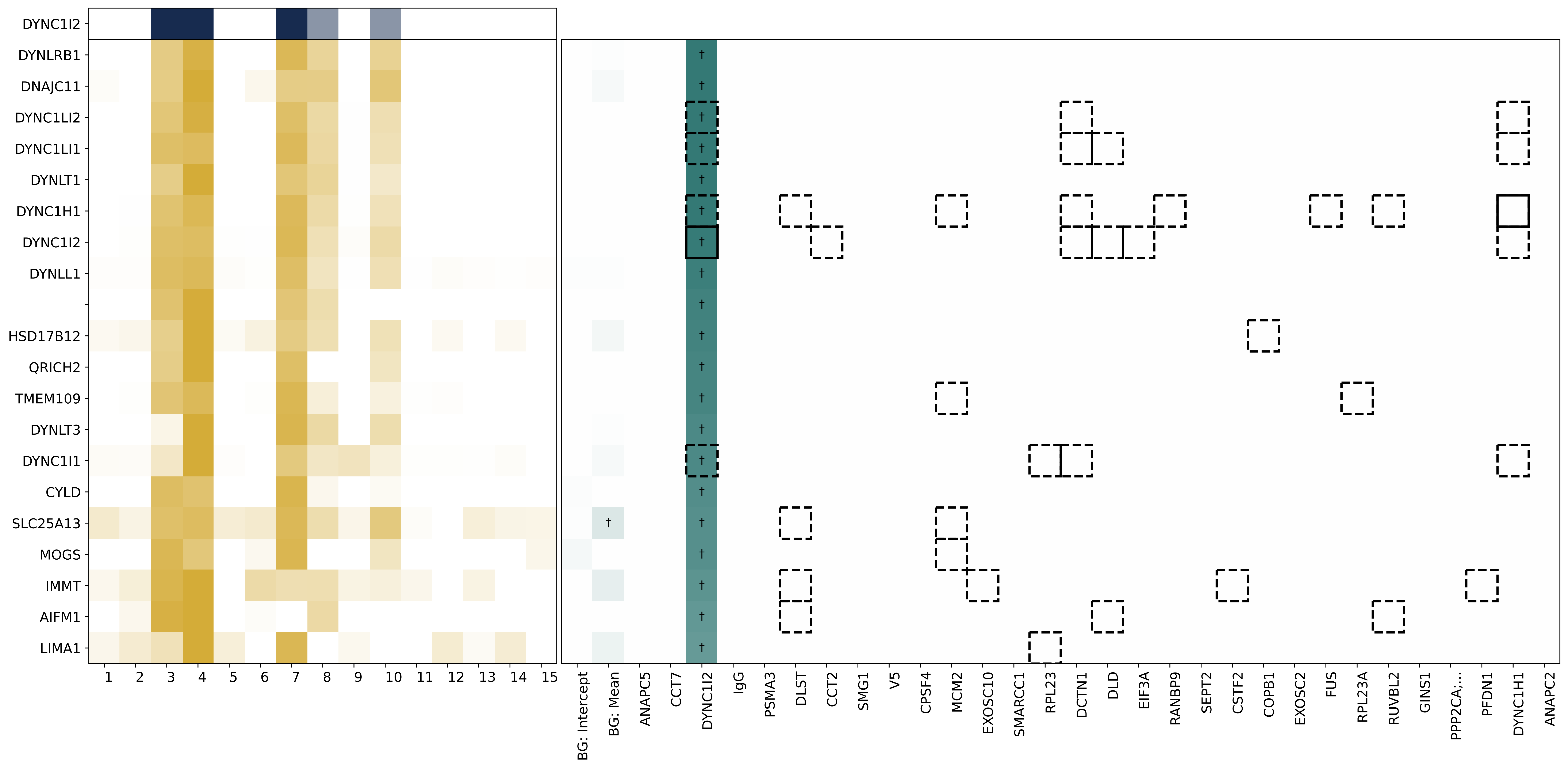

DYNC1I2
DYNLRB1
DNAJC11
DYNC1LI2
DYNC1LI1
DYNLT1
DYNC1H1
DYNC1I2
DYNLL1
HSD17B12
QRICH2
TMEM109
DYNLT3
DYNC1I1
CYLD
SLC25A13
MOGS
IMMT
AIFM1
LIMA1
1 2 3 4 5 6 7 8 9 10 11 12 13 14 15
BG: Intercept
BG: Mean
ANAPC5
CCT7
DYNC1I2
IgG
PSMA3
DLST
CCT2
SMG1
V5
CPSF4
MCM2
EXOSC10
SMARCC1
RPL23
DCTN1
DLD
EIF3A
RANBP9
SEPT2
CSTF2
COPB1
EXOSC2
FUS
RPL23A
RUVBL2
GINS1
PPP2CA;...
PFDN1
DYNC1H1
ANAPC2

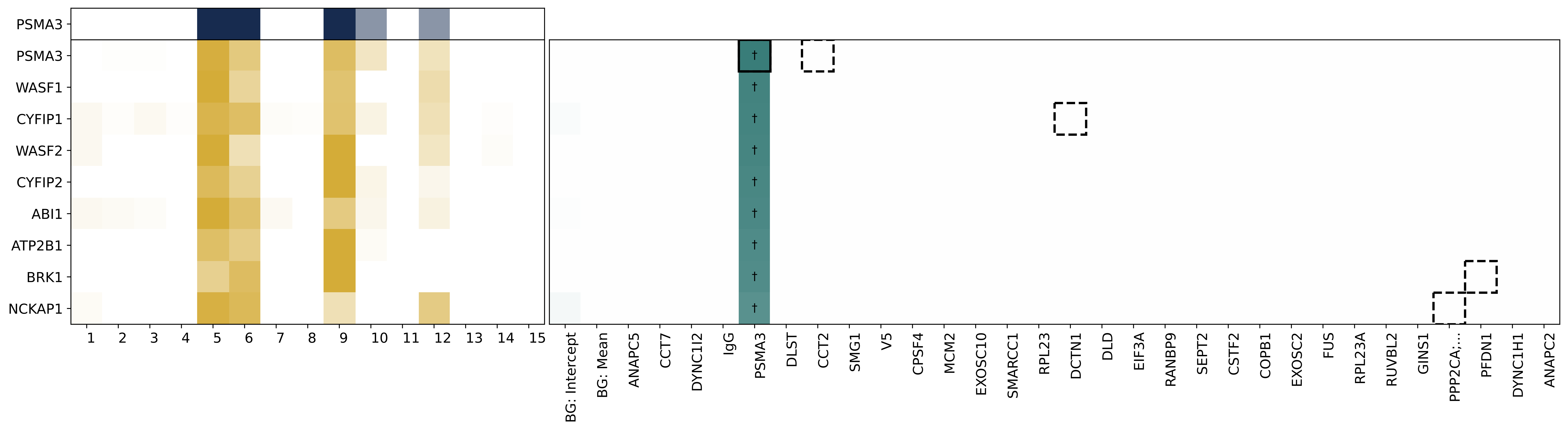
PSMA3
PSMA3
WASF1
CYFIP1
WASF2
CYFIP2
ABI1
ATP2B1
BRK1
NCKAP1
1 2 3 4 5 6 7 8 9 10 11 12 13 14 15
†
BG: Intercept
BG: Mean
ANAPC5
CCT7
DYNC1I2
IgG
PSMA3
DLST
CCT2
SMG1
V5
CPSF4
MCM2
EXOSC10
SMARCC1
RPL23
DCTN1
DLD
EIF3A
RANBP9
SEPT2
CSTF2
COPB1
EXOSC2
FUS
RPL23A
RUVBL2
GINS1
PPP2CA;...
PFDN1
DYNC1H1
ANAPC2

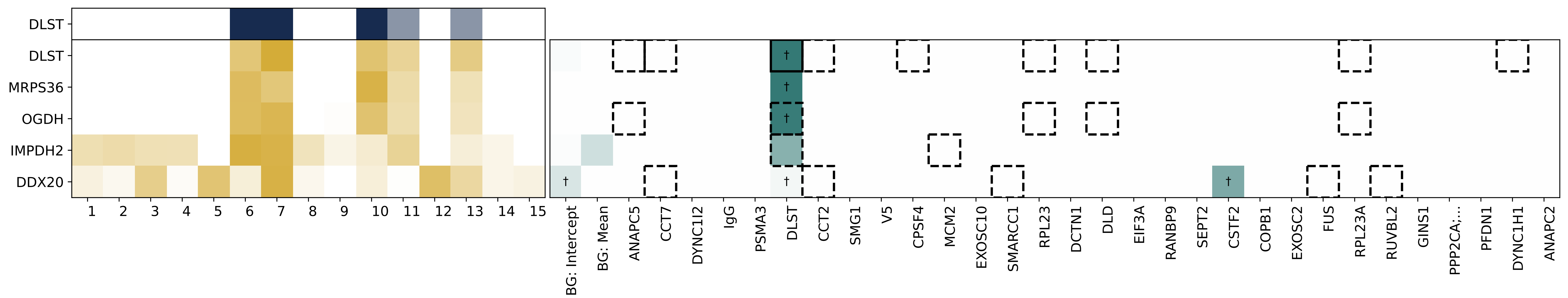
DLST
DLST
MRPS36
OGDH
IMPDH2
DDX20
1 2 3 4 5 6 7 8 9 10 11 12 13 14 15
†
BG: Intercept
BG: Mean
ANAPC5
CCT7
DYNC1I2
IgG
PSMA3
DLST
CCT2
SMG1
V5
CPSF4
MCM2
EXOSC10
SMARCC1
RPL23
DCTN1
DLD
EIF3A
RANBP9
SEPT2
CSTF2
COPB1
EXOSC2
FUS
RPL23A
RUVBL2
GINS1
PPP2CA;...
PFDN1
DYNC1H1
ANAPC2

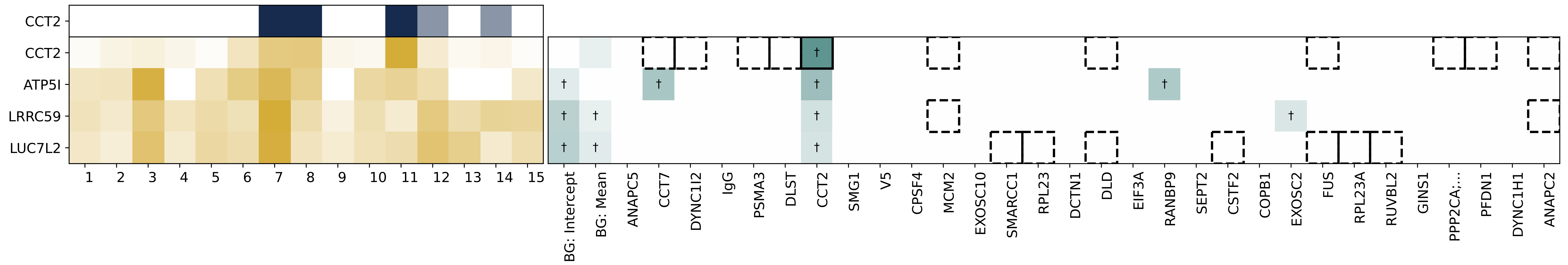
CCT2
CCT2
ATP5I
LRRC59
LUC7L2
BG: Intercept
BG: Mean
ANAPC5
CCT7
DYNC1I2
IgG
PSMA3
DLST
CCT2
SMG1
V5
CPSF4
MCM2
EXOSC10
SMARCC1
RPL23
DCTN1
DLD
EIF3A
RANBP9
SEPT2
CSTF2
COPB1
EXOSC2
FUS
RPL23A
RUVBL2
GINS1
PPP2CA;...
PFDN1
DYNC1H1
ANAPC2

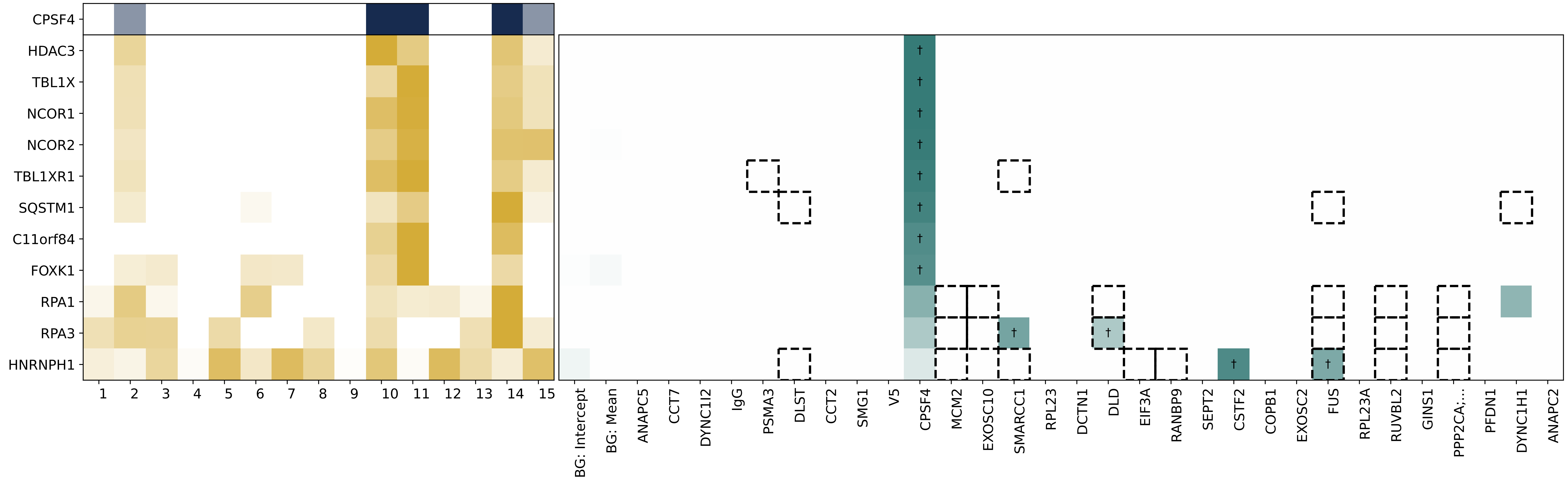
CPSF4
HDAC3
TBL1X
NCOR1
NCOR2
TBL1XR1
SQSTM1
C11orf84
FOXK1
RPA1
RPA3
HNRNPH1
1 2 3 4 5 6 7 8 9 10 11 12 13 14 15
BG: Intercept
BG: Mean
ANAPC5
CCT7
DYNC1I2
IgG
PSMA3
DLST
CCT2
SMG1
V5
CPSF4
MCM2
EXOSC10
SMARCC1
RPL23
DCTN1
DLD
EIF3A
RANBP9
SEPT2
CSTF2
COPB1
EXOSC2
FUS
RPL23A
RUVBL2
GINS1
PPP2CA;...
PFDN1
DYNC1H1
ANAPC2

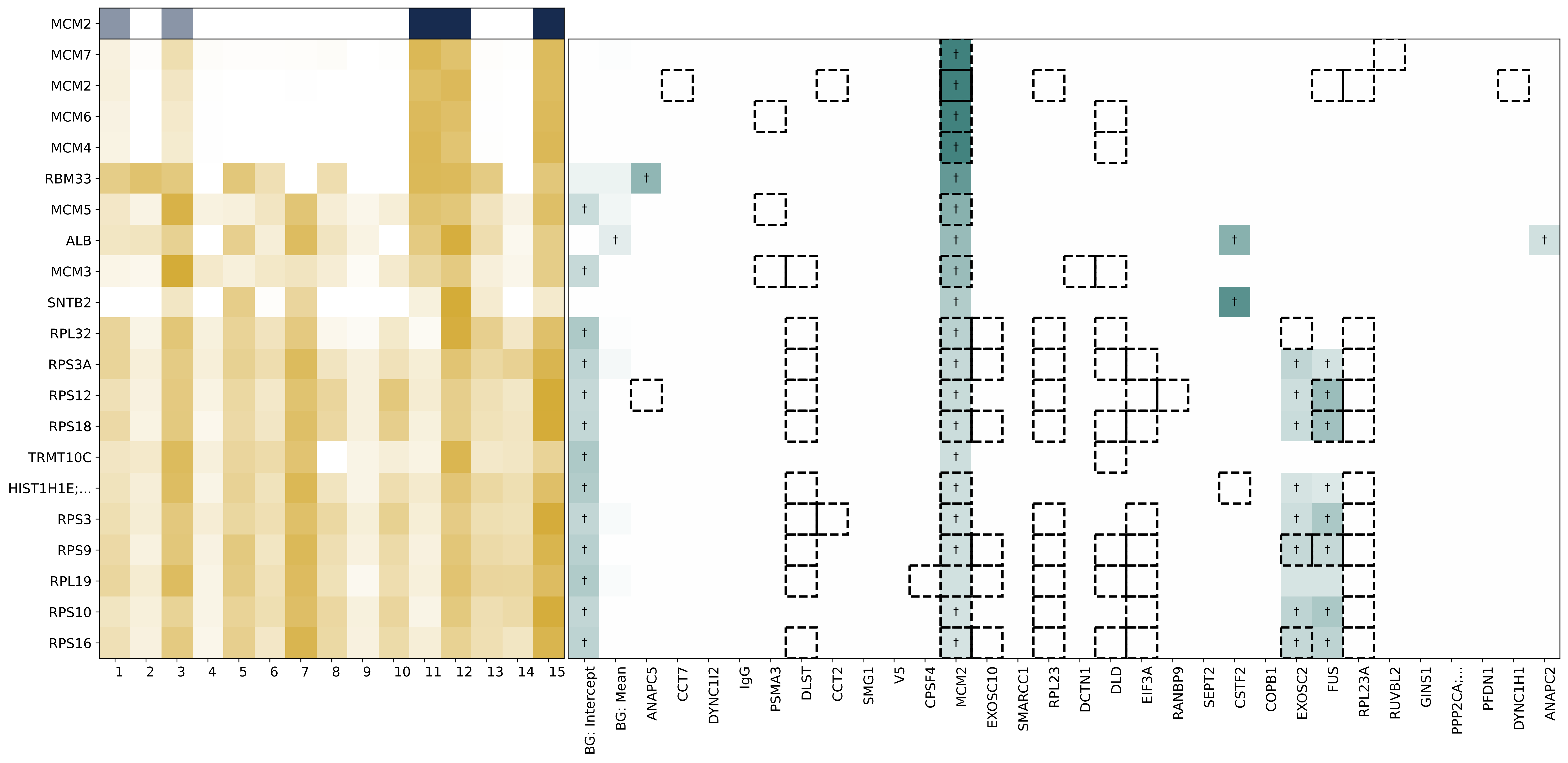
MCM2
MCM7
MCM2
MCM6
MCM4
RBM33
MCM5
ALB
MCM3
SNTB2
RPL32
RPS3A
RPS12
RPS18
TRMT10C
HIST1H1E;...
RPS3
RPS9
RPL19
RPS10
RPS16
1 2 3 4 5 6 7 8 9 10 11 12 13 14 15
BG: Intercept
BG: Mean
ANAPC5
CCT7
DYNC1I2
IgG
PSMA3
DLST
CCT2
SMG1
V5
CPSF4
MCM2
EXOSC10
SMARCC1
RPL23
DCTN1
DLD
EIF3A
RANBP9
SEPT2
CSTF2
COPB1
EXOSC2
FUS
RPL23A
RUVBL2
GINS1
PPP2CA;...
PFDN1
DYNC1H1
ANAPC2

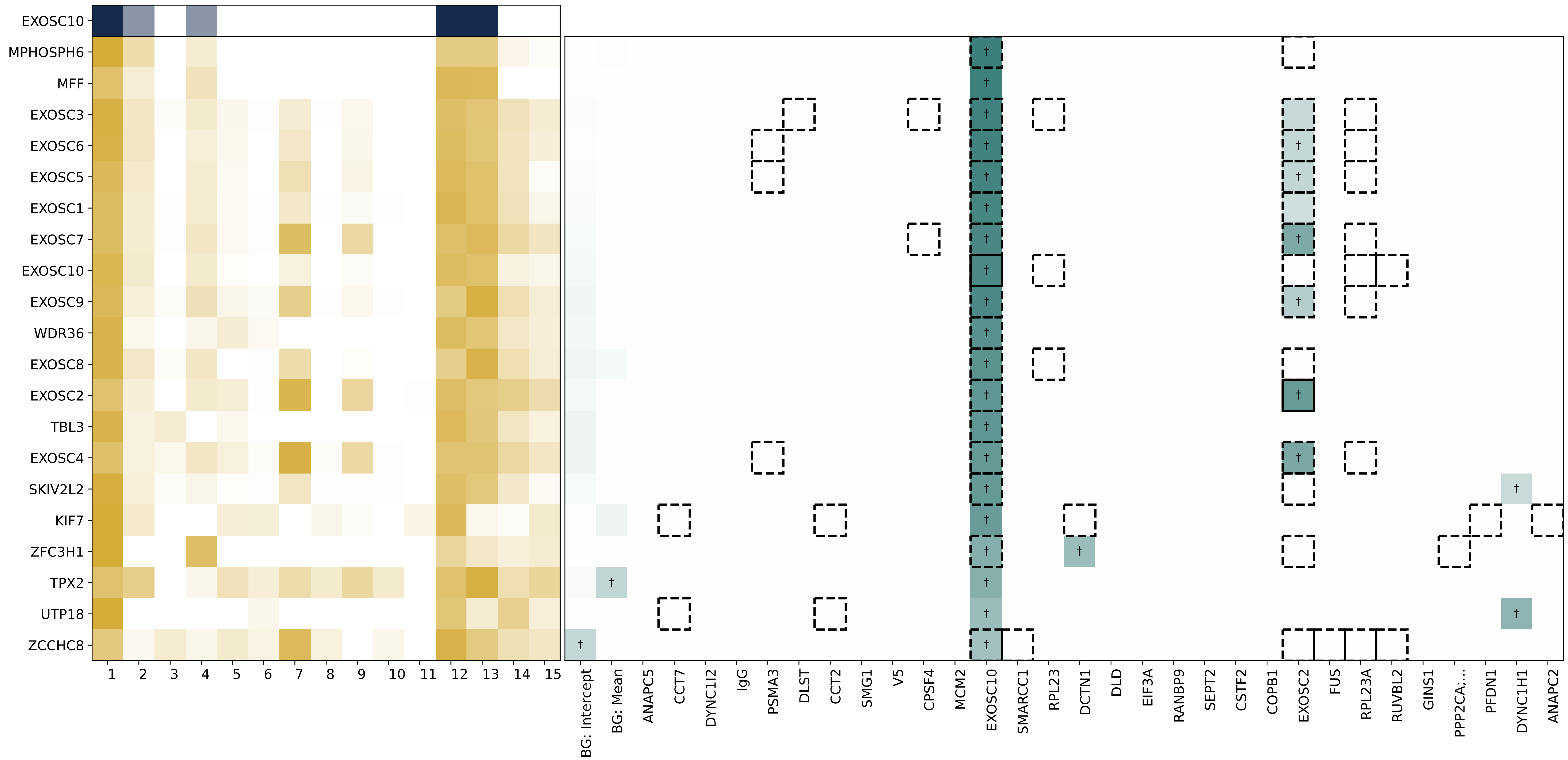

EXOSC10
MPHOSPH6
MFF
EXOSC3
EXOSC6
EXOSC5
EXOSC1
EXOSC7
EXOSC10
EXOSC9
WDR36
EXOSC8
EXOSC2
TBL3
EXOSC4
SKIV2L2
KIF7
ZFC3H1
TPX2
UTP18
ZCCHC8
1 2 3 4 5 6 7 8 9 10 11 12 13 14 15
BG: Intercept
BG: Mean
ANAPC5
CCT7
DYNC1I2
IgG
PSMA3
DLST
CCT2
SMG1
V5
CPSF4
MCM2
EXOSC10
SMARCC1
RPL23
DCTN1
DLD
EIF3A
RANBP9
SEPT2
CSTF2
COPB1
EXOSC2
FUS
RPL23A
RUVBL2
GINS1
PPP2CA;...
PFDN1
DYNC1H1
ANAPC2

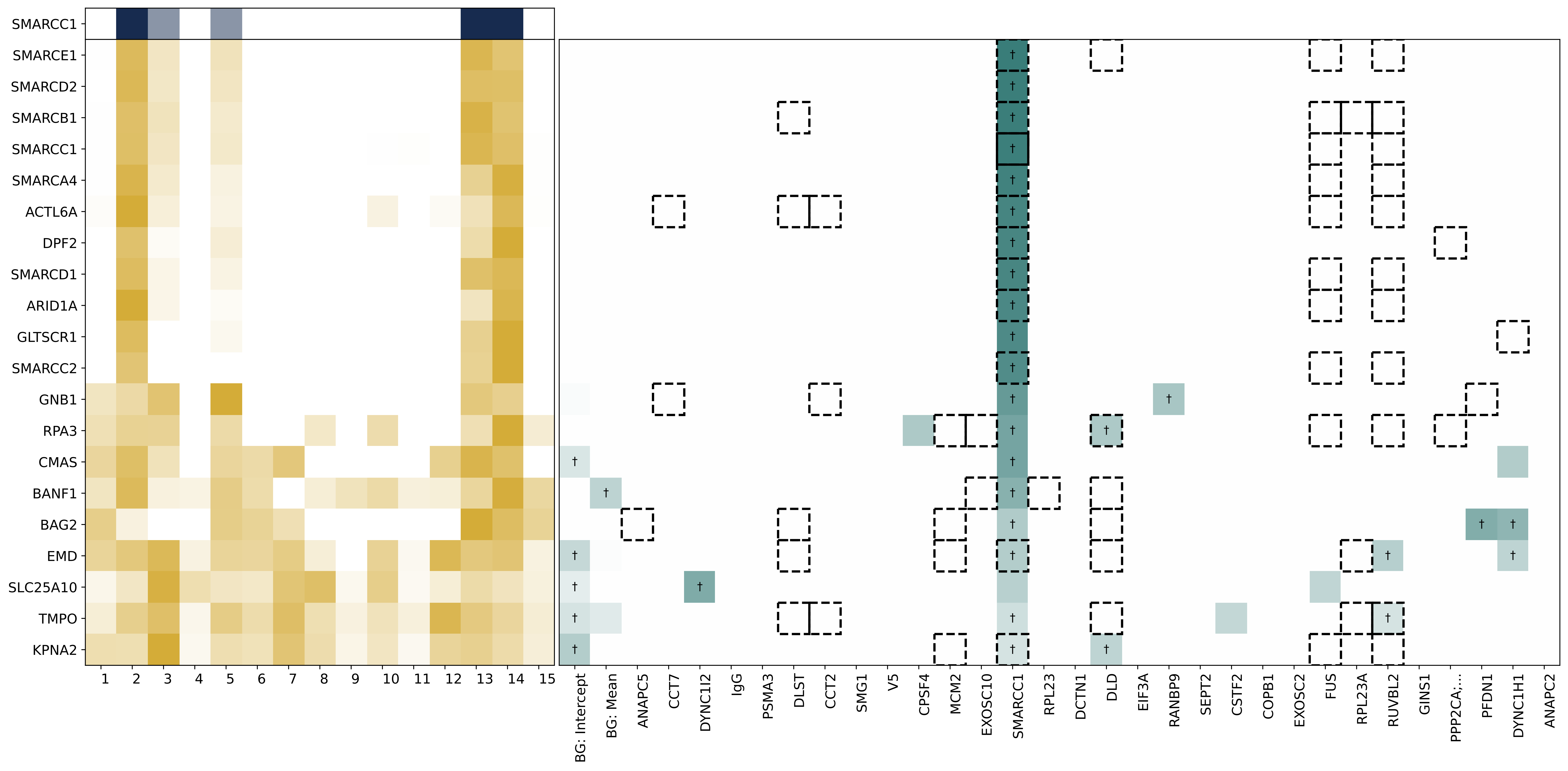

SMARCC1
SMARCE1
SMARCD2
SMARCB1
SMARCC1
SMARCA4
ACTL6A
DPF2
SMARCD1
ARID1A
GLTSCR1
SMARCC2
GNB1
RPA3
CMAS
BANF1
BAG2
EMD
SLC25A10
TMPO
KPNA2
1 2 3 4 5 6 7 8 9 10 11 12 13 14 15
BG: Intercept
BG: Mean
ANAPC5
CCT7
DYNC1I2
IgG
PSMA3
DLST
CCT2
SMG1
V5
CPSF4
MCM2
EXOSC10
SMARCC1
RPL23
DCTN1
DLD
EIF3A
RANBP9
SEPT2
CSTF2
COPB1
EXOSC2
FUS
RPL23A
RUVBL2
GINS1
PPP2CA;...
PFDN1
DYNC1H1
ANAPC2

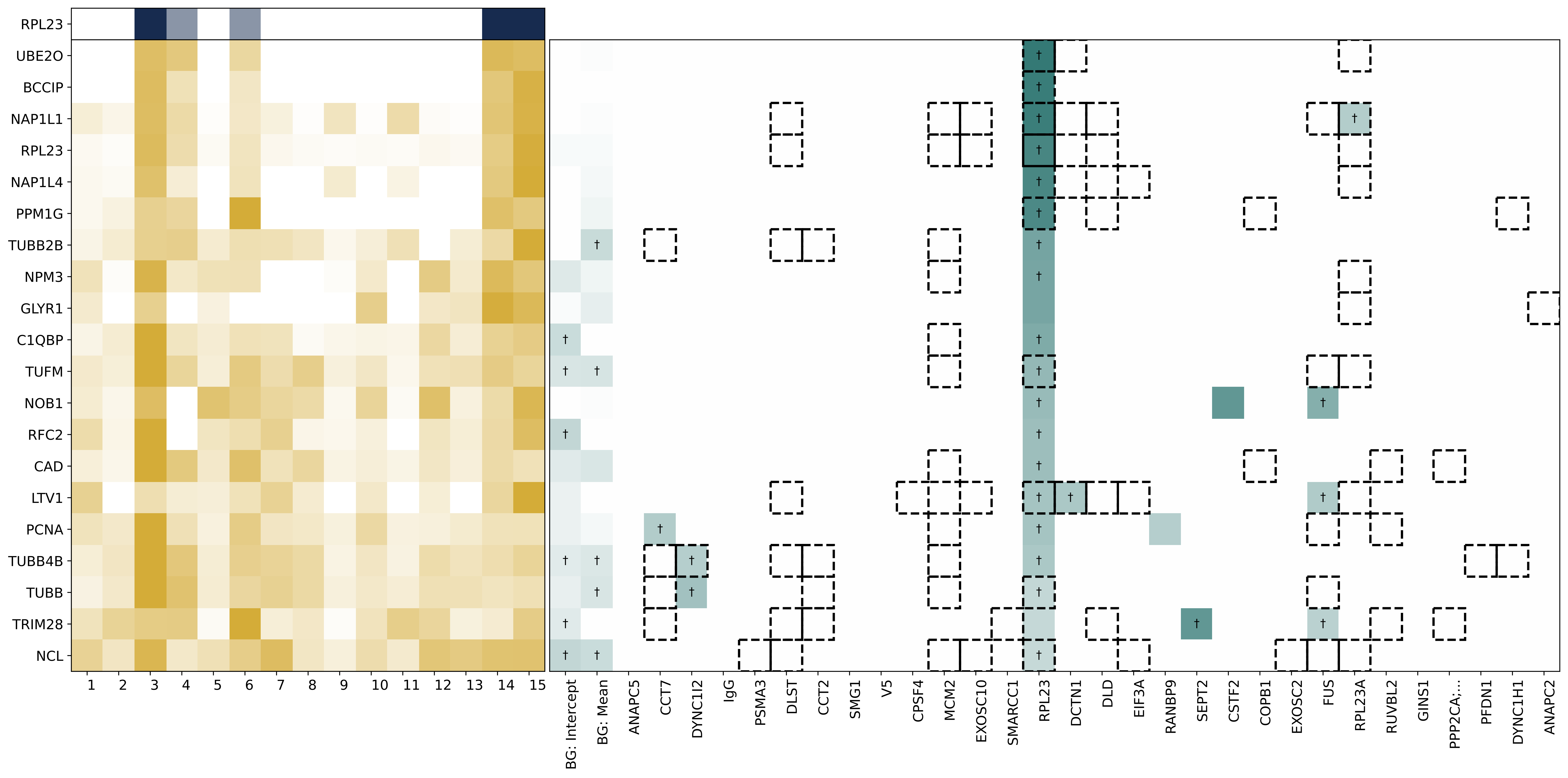
RPL23
UBE2O
BCCIP
NAP1L1
RPL23
NAP1L4
PPM1G
TUBB2B
NPM3
GLYR1
C1QBP
TUFM
NOB1
RFC2
CAD
LTV1
PCNA
TUBB4B
TUBB
TRIM28
NCL
1 2 3 4 5 6 7 8 9 10 11 12 13 14 15
BG: Intercept
BG: Mean
ANAPC5
CCT7
DYNC1I2
IgG
PSMA3
DLST
CCT2
SMG1
V5
CPSF4
MCM2
EXOSC10
SMARCC1
RPL23
DCTN1
DLD
EIF3A
RANBP9
SEPT2
CSTF2
COPB1
EXOSC2
FUS
RPL23A
RUVBL2
GINS1
PPP2CA;...
PFDN1
DYNC1H1
ANAPC2

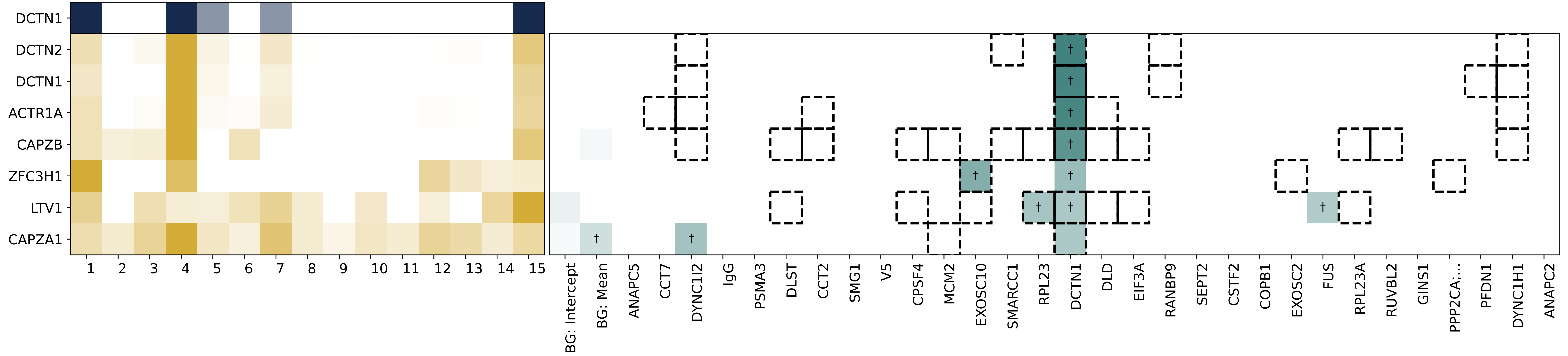

DCTN1
DCTN2
DCTN1
ACTR1A
CAPZB
ZFC3H1
LTV1
CAPZA1
†
BG: Intercept
BG: Mean
ANAPC5
CCT7
DYNC1I2
IgG
PSMA3
DLST
CCT2
SMG1
V5
CPSF4
MCM2
EXOSC10
SMARCC1
RPL23
DCTN1
DLD
EIF3A
RANBP9
SEPT2
CSTF2
COPB1
EXOSC2
FUS
RPL23A
RUVBL2
GINS1
PPP2CA;...
PFDN1
DYNC1H1
ANAPC2

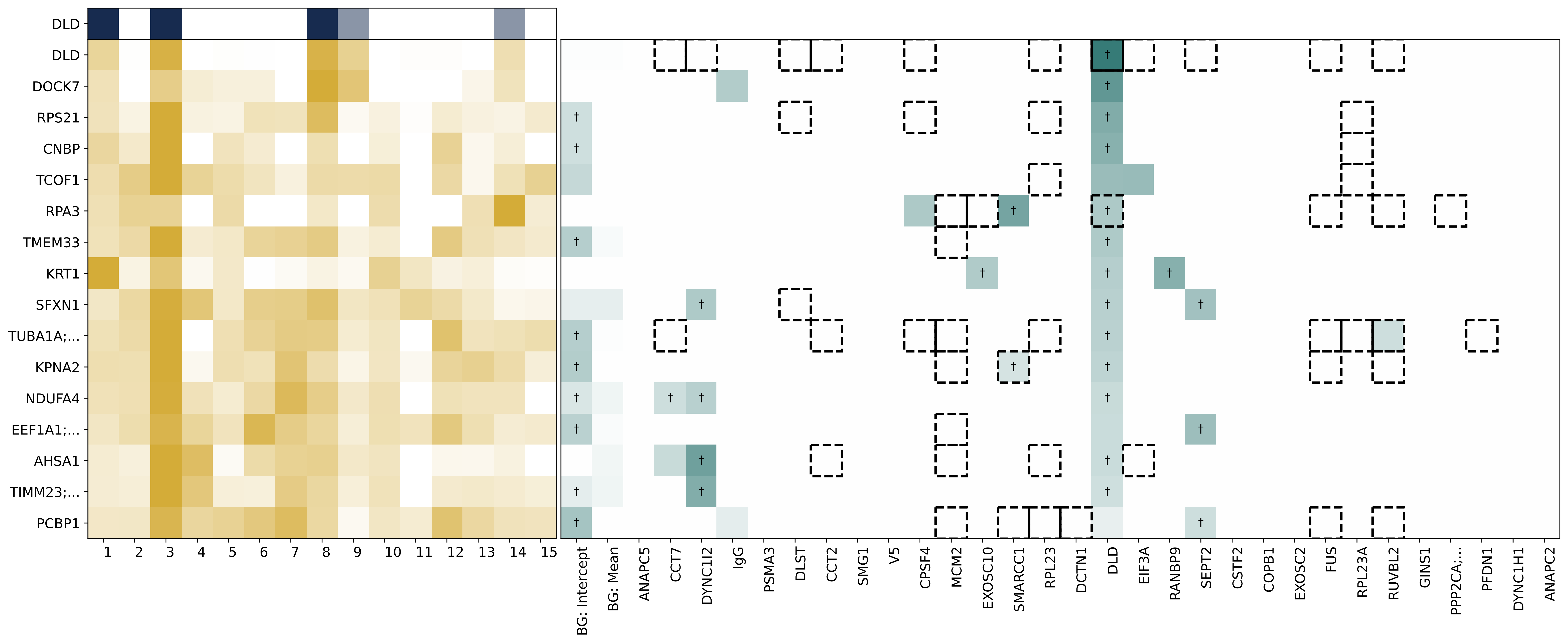

DLD
DLD
DOCK7
RPS21
CNBP
TCOF1
RPA3
TMEM33
KRT1
SFXN1
TUBA1A;...
KPNA2
NDUFA4
EEF1A1;...
AHSA1
TIMM23;...
PCBP1
1 2 3 4 5 6 7 8 9 10 11 12 13 14 15
BG: Intercept
BG: Mean
ANAPC5
CCT7
DYNC1I2
IgG
PSMA3
DLST
CCT2
SMG1
V5
CPSF4
MCM2
EXOSC10
SMARCC1
RPL23
DCTN1
DLD
EIF3A
RANBP9
SEPT2
CSTF2
COPB1
EXOSC2
FUS
RPL23A
RUVBL2
GINS1
PPP2CA;...
PFDN1
DYNC1H1
ANAPC2

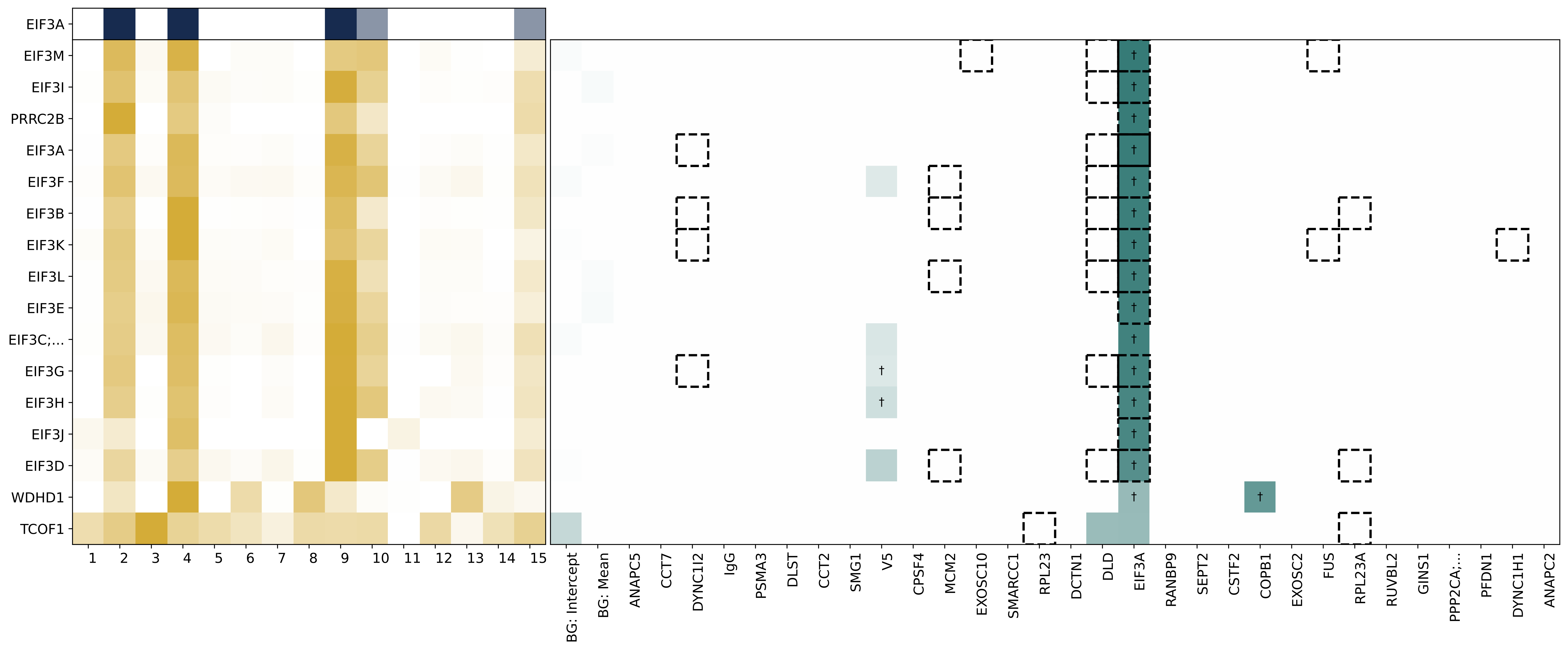
EIF3A
EIF3M
EIF3I
PRRC2B
EIF3A
EIF3F
EIF3B
EIF3K
EIF3L
EIF3E
EIF3C;...
EIF3G
EIF3H
EIF3J
EIF3D
WDHD1
TCOF1
1 2 3 4 5 6 7 8 9 10 11 12 13 14 15
BG: Intercept
BG: Mean
ANAPC5
CCT7
DYNC1I2
IgG
PSMA3
DLST
CCT2
SMG1
V5
CPSF4
MCM2
EXOSC10
SMARCC1
RPL23
DCTN1
DLD
EIF3A
RANBP9
SEPT2
CSTF2
COPB1
EXOSC2
FUS
RPL23A
RUVBL2
GINS1
PPP2CA;...
PFDN1
DYNC1H1
ANAPC2

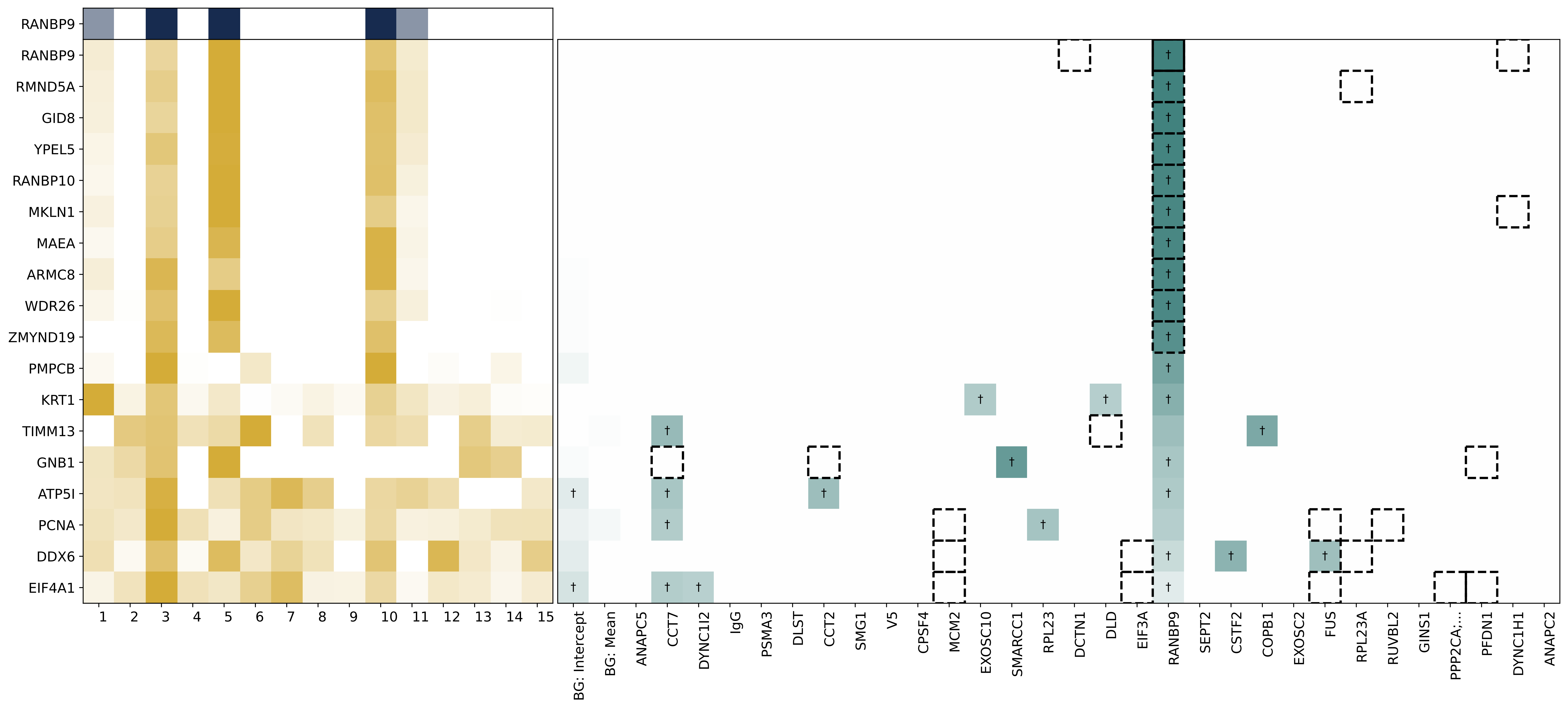
RANBP9
RANBP9
RMND5A
GID8
YPEL5
RANBP10
MKLN1
MAEA
ARMC8
WDR26
ZMYND19
PMPCB
KRT1
TIMM13
GNB1
ATP5I
PCNA
DDX6
EIF4A1
1 2 3 4 5 6 7 8 9 10 11 12 13 14 15
BG: Intercept
BG: Mean
ANAPC5
CCT7
DYNC1I2
IgG
PSMA3
DLST
CCT2
SMG1
V5
CPSF4
MCM2
EXOSC10
SMARCC1
RPL23
DCTN1
DLD
EIF3A
RANBP9
SEPT2
CSTF2
COPB1
EXOSC2
FUS
RPL23A
RUVBL2
GINS1
PPP2CA;...
PFDN1
DYNC1H1
ANAPC2

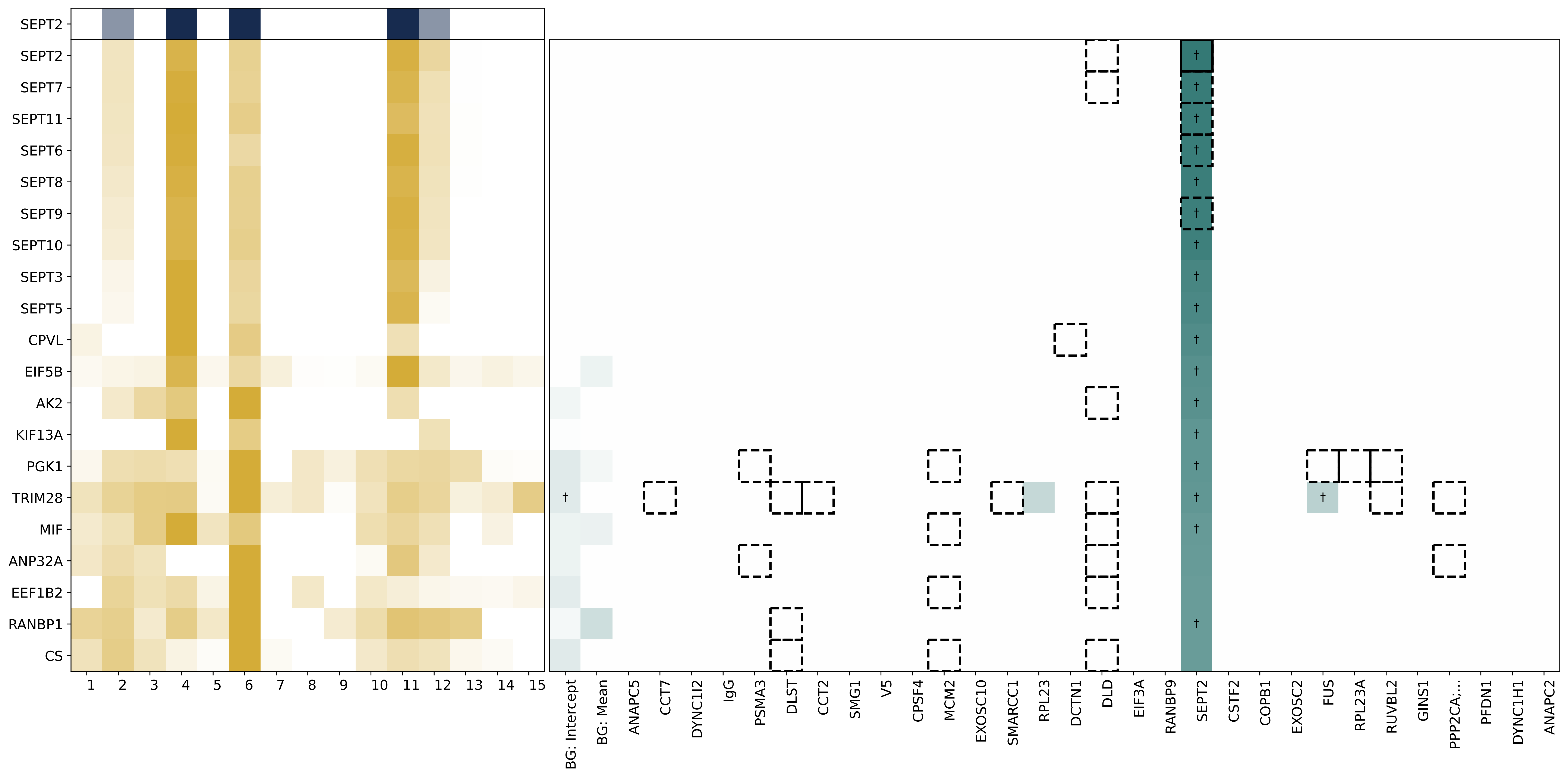

SEPT2
SEPT2
SEPT7
SEPT11
SEPT6
SEPT8
SEPT9
SEPT10
SEPT3
SEPT5
CPVL
EIF5B
AK2
KIF13A
PGK1
TRIM28
MIF
ANP32A
EEF1B2
RANBP1
CS
BG: Intercept
BG: Mean
ANAPC5
CCT7
DYNC1I2
IgG
PSMA3
DLST
CCT2
SMG1
V5
CPSF4
MCM2
EXOSC10
SMARCC1
RPL23
DCTN1
DLD
EIF3A
RANBP9
SEPT2
CSTF2
COPB1
EXOSC2
FUS
RPL23A
RUVBL2
GINS1
PPP2CA;...
PFDN1
DYNC1H1
ANAPC2

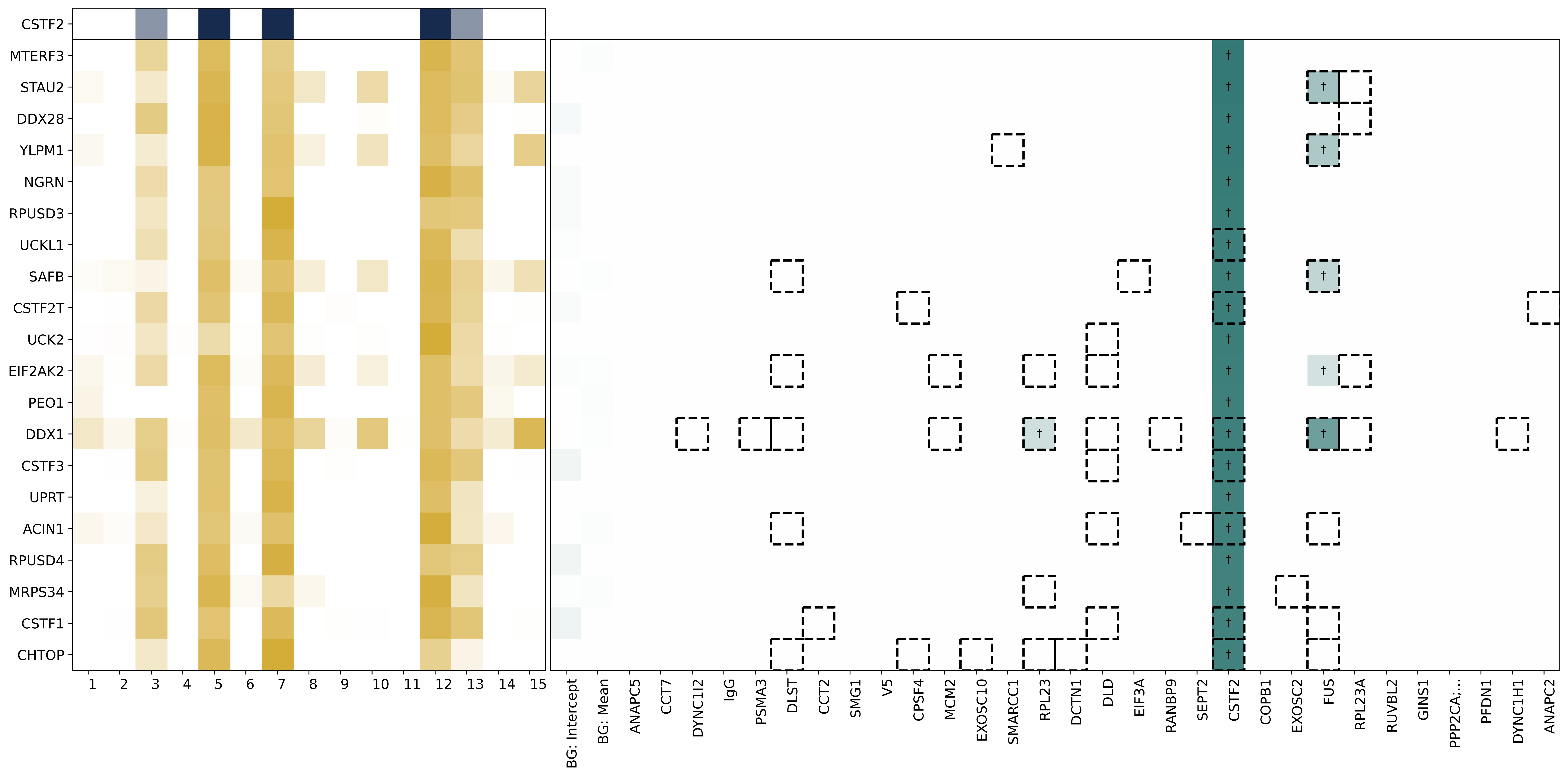

CSTF2
MTERF3
STAU2
DDX28
YLPM1
NGRN
RPUSD3
UCKL1
SAFB
CSTF2T
UCK2
EIF2AK2
PEO1
DDX1
CSTF3
UPRT
ACIN1
RPUSD4
MRPS34
CSTF1
CHTOP
BG: Intercept
BG: Mean
ANAPC5
CCT7
DYNC1I2
IgG
PSMA3
DLST
CCT2
SMG1
V5
CPSF4
MCM2
EXOSC10
SMARCC1
RPL23
DCTN1
DLD
EIF3A
RANBP9
SEPT2
CSTF2
COPB1
EXOSC2
FUS
RPL23A
RUVBL2
GINS1
PPP2CA;...
PFDN1
DYNC1H1
ANAPC2

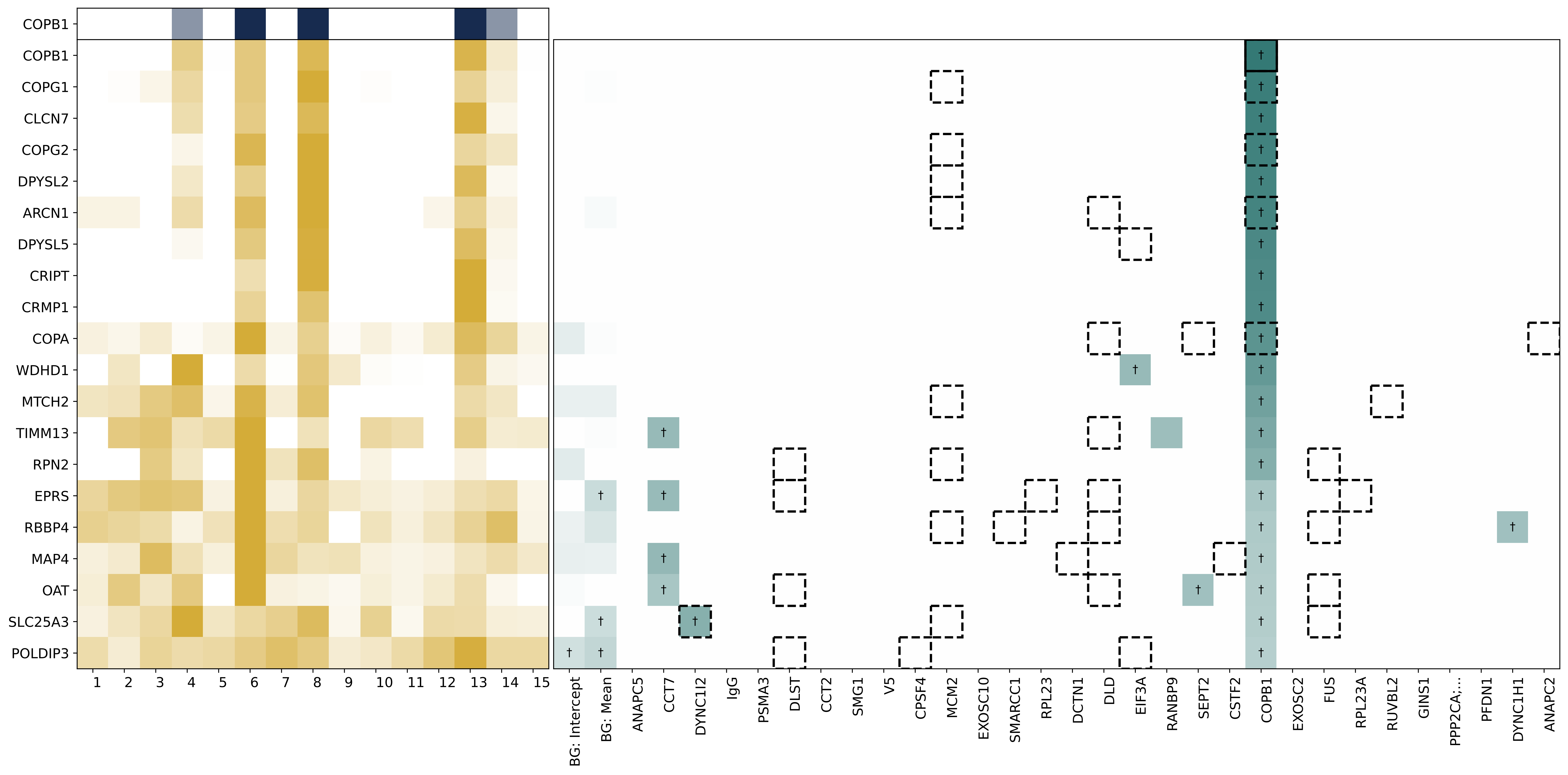

COPB1
COPB1
COPG1
CLCN7
COPG2
DPYSL2
ARCN1
DPYSL5
CRIPT
CRMP1
COPA
WDHD1
MTCH2
TIMM13
RPN2
EPRS
RBBP4
MAP4
OAT
SLC25A3
POLDIP3
BG: Intercept
BG: Mean
ANAPC5
CCT7
DYNC1I2
IgG
PSMA3
DLST
CCT2
SMG1
V5
CPSF4
MCM2
EXOSC10
SMARCC1
RPL23
DCTN1
DLD
EIF3A
RANBP9
SEPT2
CSTF2
COPB1
EXOSC2
FUS
RPL23A
RUVBL2
GINS1
PPP2CA;...
PFDN1
DYNC1H1
ANAPC2

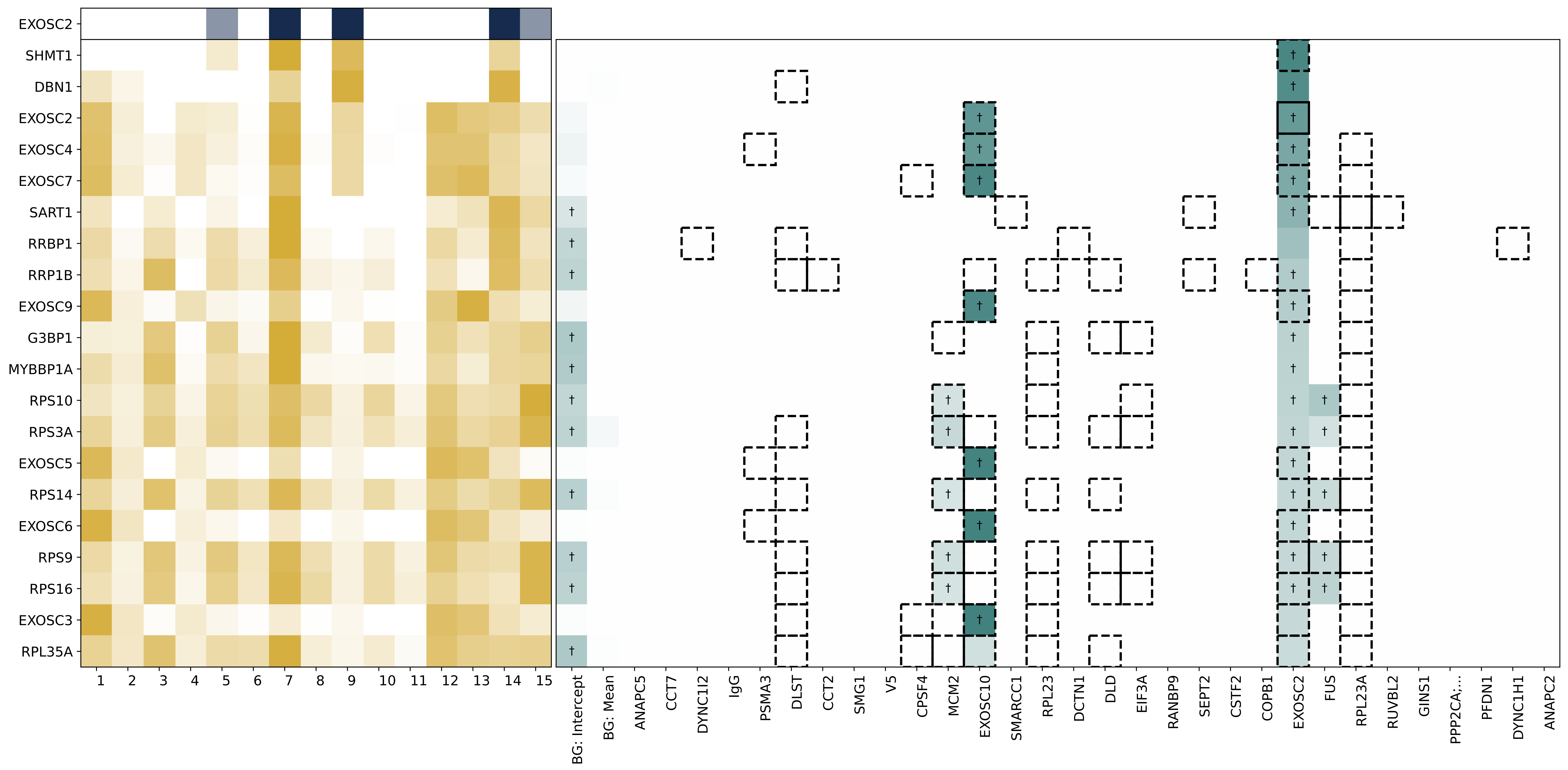

EXOSC2
SHMT1
DBN1
EXOSC2
EXOSC4
EXOSC7
SART1
RRBP1
RRP1B
EXOSC9
G3BP1
MYBBP1A
RPS10
RPS3A
EXOSC5
RPS14
EXOSC6
RPS9
RPS16
EXOSC3
RPL35A
1 2 3 4 5 6 7 8 9 10 11 12 13 14 15
BG: Intercept
BG: Mean
ANAPC5
CCT7
DYNC1I2
IgG
PSMA3
DLST
CCT2
SMG1
V5
CPSF4
MCM2
EXOSC10
SMARCC1
RPL23
DCTN1
DLD
EIF3A
RANBP9
SEPT2
CSTF2
COPB1
EXOSC2
FUS
RPL23A
RUVBL2
GINS1
PPP2CA;...
PFDN1
DYNC1H1
ANAPC2
†

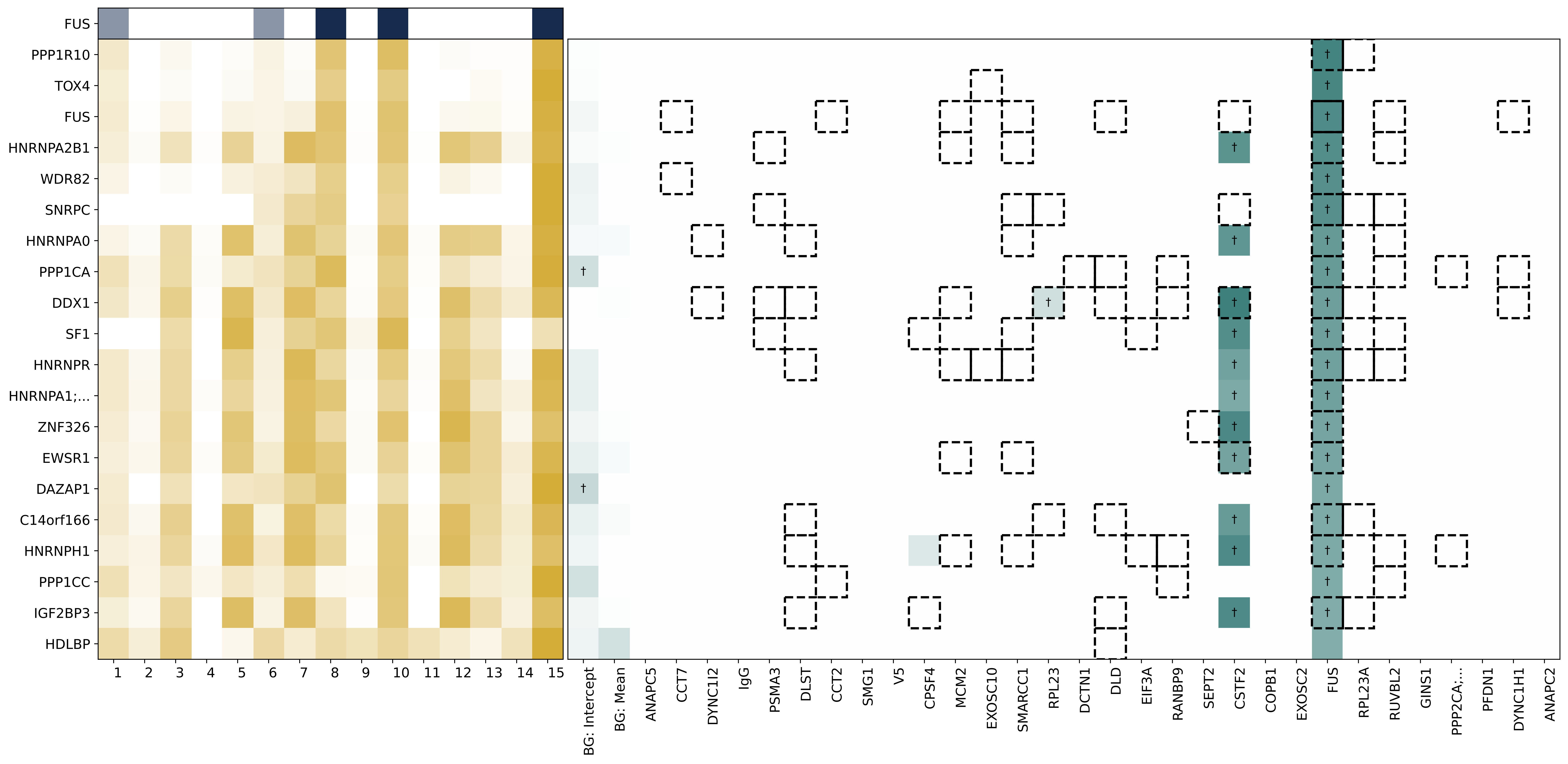
FUS
PPP1R10
TOX4
FUS
HNRNPA2B1
WDR82
SNRPC
HNRNPA0
PPP1CA
DDX1
SF1
HNRNPR
HNRNPA1;...
ZNF326
EWSR1
DAZAP1
C14orf166
HNRNPH1
PPP1CC
IGF2BP3
HDLBP
1 2 3 4 5 6 7 8 9 10 11 12 13 14 15
BG: Intercept
BG: Mean
ANAPC5
CCT7
DYNC1I2
IgG
PSMA3
DLST
CCT2
SMG1
V5
CPSF4
MCM2
EXOSC10
SMARCC1
RPL23
DCTN1
DLD
EIF3A
RANBP9
SEPT2
CSTF2
COPB1
EXOSC2
FUS
RPL23A
RUVBL2
GINS1
PPP2CA;...
PFDN1
DYNC1H1
ANAPC2

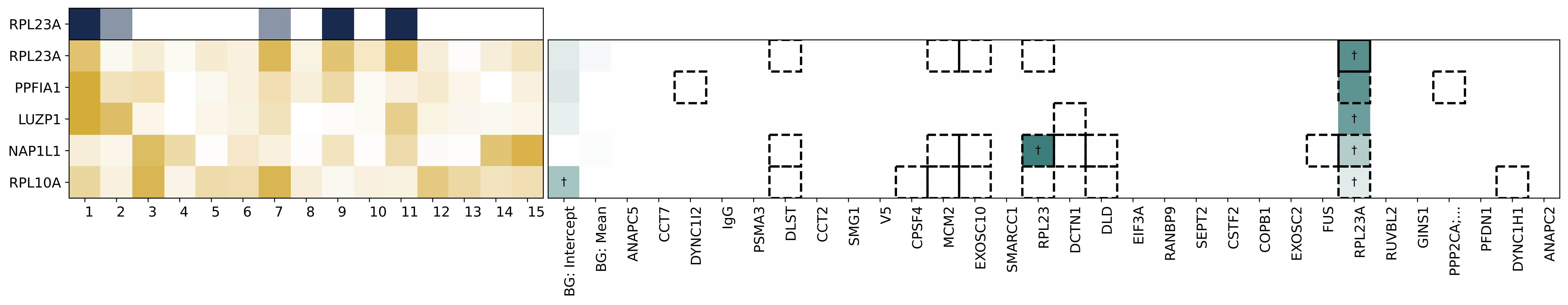
RPL23A
RPL23A
PPFIA1
LUZP1
NAP1L1
RPL10A
1 2 3 4 5 6 7 8 9 10 11 12 13 14 15
BG: Intercept
BG: Mean
ANAPC5
CCT7
DYNC1I2
IgG
PSMA3
DLST
CCT2
SMG1
V5
CPSF4
MCM2
EXOSC10
SMARCC1
RPL23
DCTN1
DLD
EIF3A
RANBP9
SEPT2
CSTF2
COPB1
EXOSC2
FUS
RPL23A
RUVBL2
GINS1
PPP2CA;...
PFDN1
DYNC1H1
ANAPC2
†

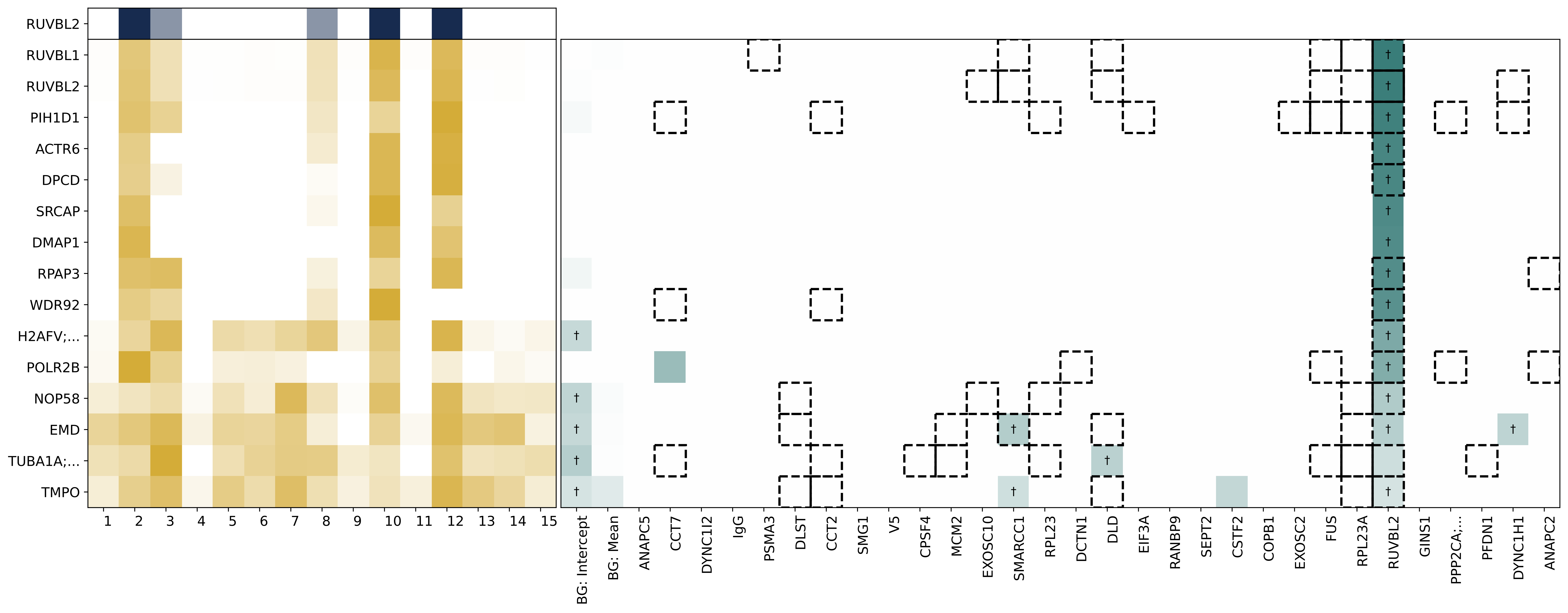

RUVBL2
RUVBL1
RUVBL2
PIH1D1
ACTR6
DPCD
SRCAP
DMAP1
RPAP3
WDR92
H2AFV;...
POLR2B
NOP58
EMD
TUBA1A;...
TMPO
1 2 3 4 5 6 7 8 9 10 11 12 13 14 15
BG: Intercept
BG: Mean
ANAPC5
CCT7
DYNC1I2
IgG
PSMA3
DLST
CCT2
SMG1
V5
CPSF4
MCM2
EXOSC10
SMARCC1
RPL23
DCTN1
DLD
EIF3A
RANBP9
SEPT2
CSTF2
COPB1
EXOSC2
FUS
RPL23A
RUVBL2
GINS1
PPP2CA;...
PFDN1
DYNC1H1
ANAPC2

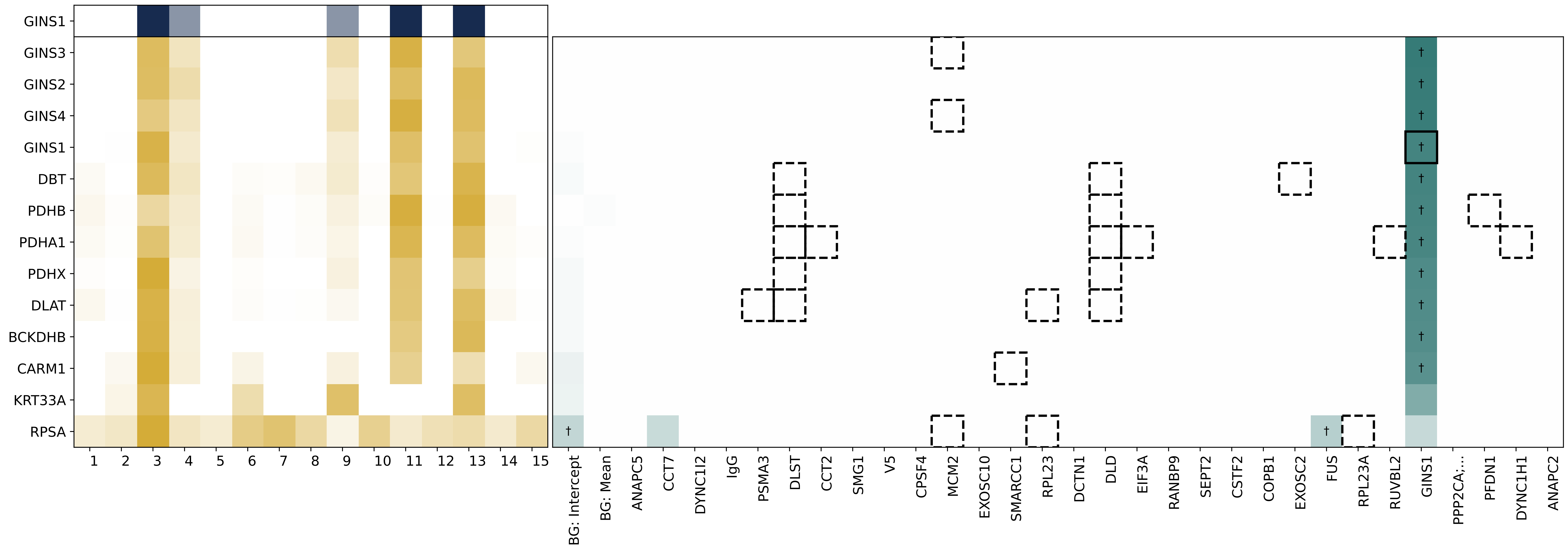
GINS1
GINS3
GINS2
GINS4
GINS1
DBT
PDHB
PDHA1
PDHX
DLAT
BCKDHB
CARM1
KRT33A
RPSA
1 2 3 4 5 6 7 8 9 10 11 12 13 14 15
BG: Intercept
BG: Mean
ANAPC5
CCT7
DYNC1I2
IgG
PSMA3
DLST
CCT2
SMG1
V5
CPSF4
MCM2
EXOSC10
SMARCC1
RPL23
DCTN1
DLD
EIF3A
RANBP9
SEPT2
CSTF2
COPB1
EXOSC2
FUS
RPL23A
RUVBL2
GINS1
PPP2CA;...
PFDN1
DYNC1H1
ANAPC2

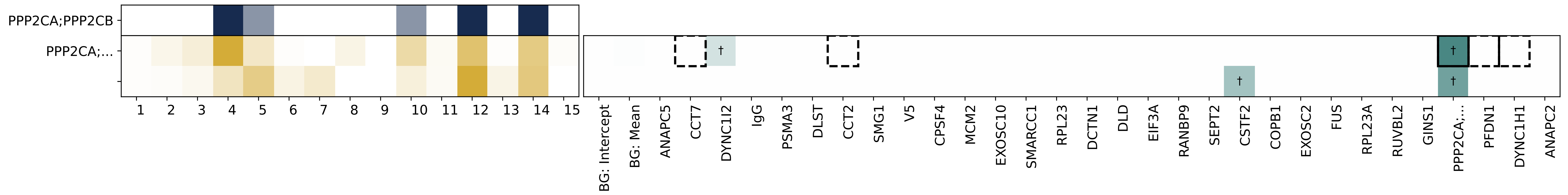

PPP2CA;PPP2CB
PPP2CA;...
†
BG: Intercept
BG: Mean
ANAPC5
CCT7
DYNC1I2
IgG
PSMA3
DLST
CCT2
SMG1
V5
CPSF4
MCM2
EXOSC10
SMARCC1
RPL23
DCTN1
DLD
EIF3A
RANBP9
SEPT2
CSTF2
COPB1
EXOSC2
FUS
RPL23A
RUVBL2
GINS1
PPP2CA;...
PFDN1
DYNC1H1
ANAPC2

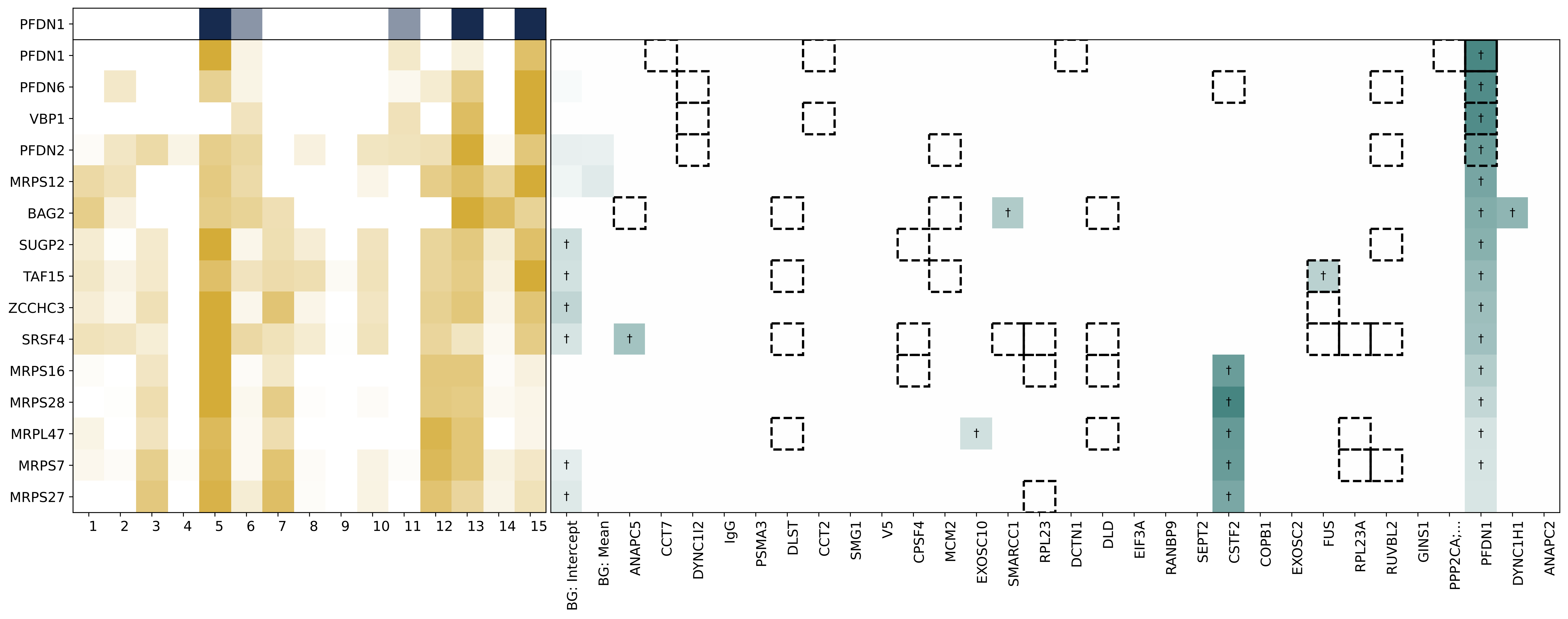
PFDN1
PFDN1
PFDN6
VBP1
PFDN2
MRPS12
BAG2
SUGP2
TAF15
ZCCHC3
SRSF4
MRPS16
MRPS28
MRPL47
MRPS7
MRPS27
1 2 3 4 5 6 7 8 9 10 11 12 13 14 15
BG: Intercept
BG: Mean
ANAPC5
CCT7
DYNC1I2
IgG
PSMA3
DLST
CCT2
SMG1
V5
CPSF4
MCM2
EXOSC10
SMARCC1
RPL23
DCTN1
DLD
EIF3A
RANBP9
SEPT2
CSTF2
COPB1
EXOSC2
FUS
RPL23A
RUVBL2
GINS1
PPP2CA;...
PFDN1
DYNC1H1
ANAPC2

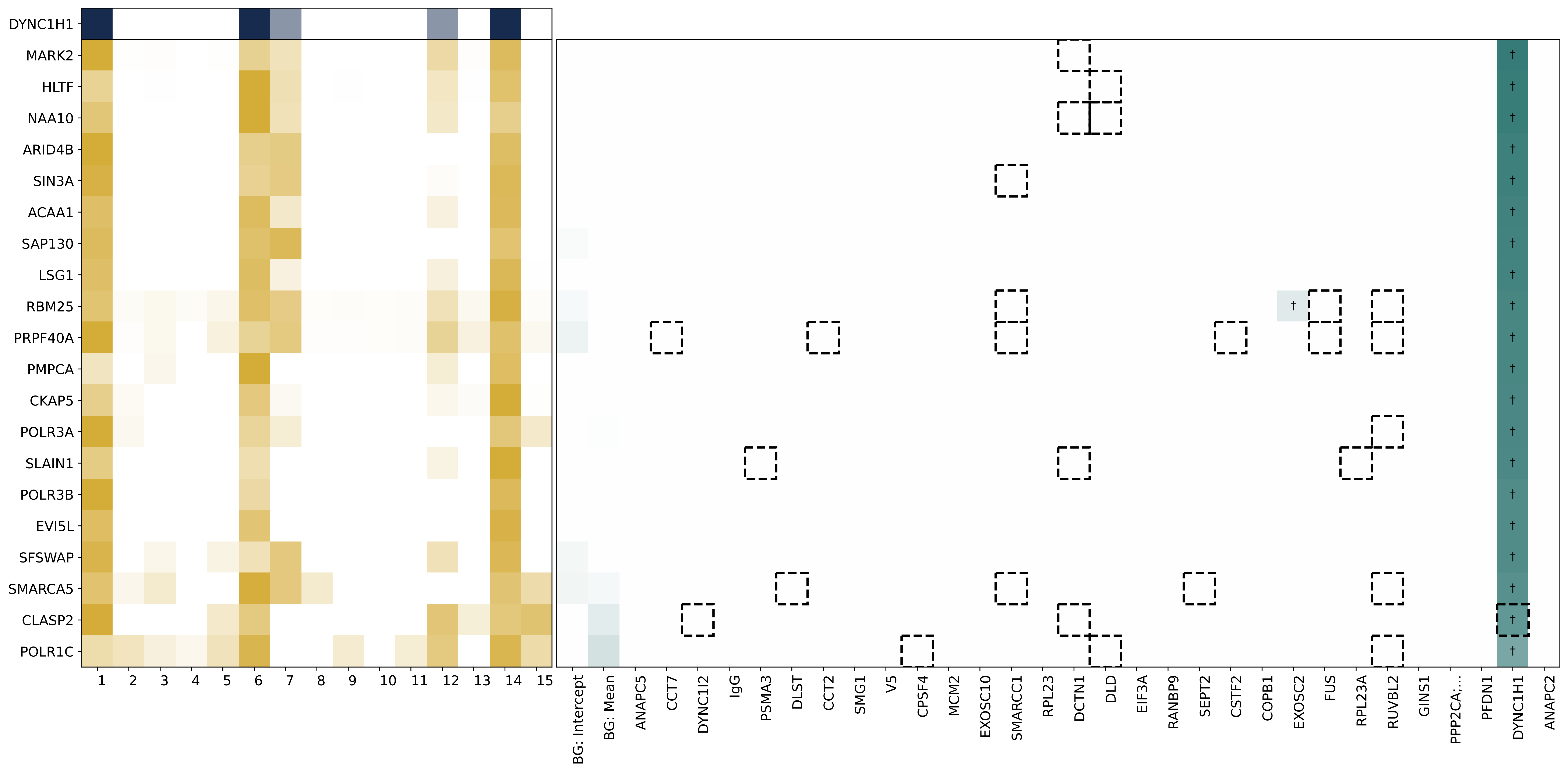
DYNC1H1
MARK2
HLTF
NAA10
ARID4B
SIN3A
ACAA1
SAP130
LSG1
RBM25
PRPF40A
PMPCA
CKAP5
POLR3A
SLAIN1
POLR3B
EVI5L
SFSWAP
SMARCA5
CLASP2
POLR1C
BG: Intercept
BG: Mean
ANAPC5
CCT7
DYNC1I2
IgG
PSMA3
DLST
CCT2
SMG1
V5
CPSF4
MCM2
EXOSC10
SMARCC1
RPL23
DCTN1
DLD
EIF3A
RANBP9
SEPT2
CSTF2
COPB1
EXOSC2
FUS
RPL23A
RUVBL2
GINS1
PPP2CA;...
PFDN1
DYNC1H1
ANAPC2

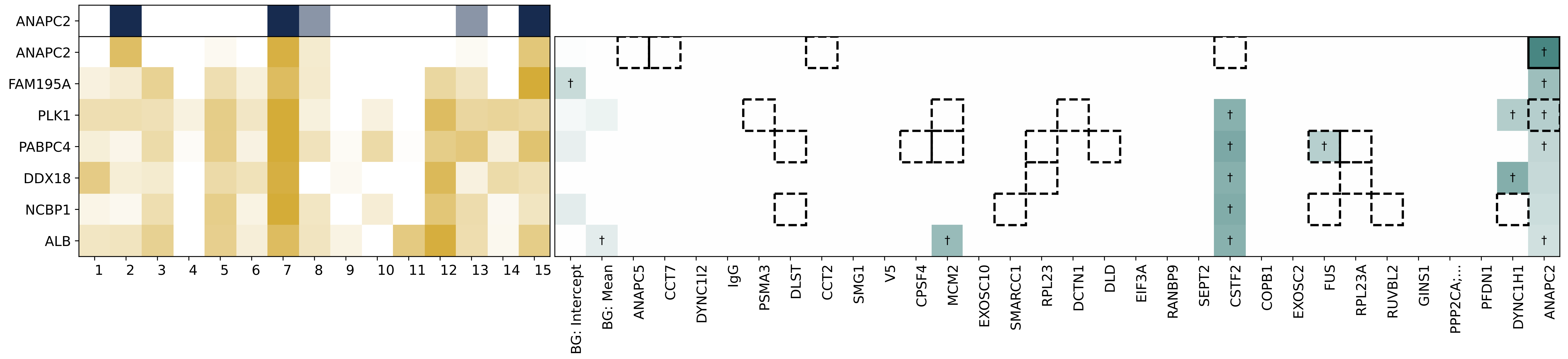
ANAPC2
ANAPC2
FAM195A
PLK1
PABPC4
DDX18
NCBP1
ALB
1 2 3 4 5 6 7 8 9 10 11 12 13 14 15
BG: Intercept
BG: Mean
ANAPC5
CCT7
DYNC1I2
IgG
PSMA3
DLST
CCT2
SMG1
V5
CPSF4
MCM2
EXOSC10
SMARCC1
RPL23
DCTN1
DLD
EIF3A
RANBP9
SEPT2
CSTF2
COPB1
EXOSC2
FUS
RPL23A
RUVBL2
GINS1
PPP2CA;...
PFDN1
DYNC1H1
ANAPC2
