## Supplemental Table 5 for "SPRINT-MS: A high-throughput platform for identifying protein-protein interactions using pooled IP-MS and sparse signal recovery"

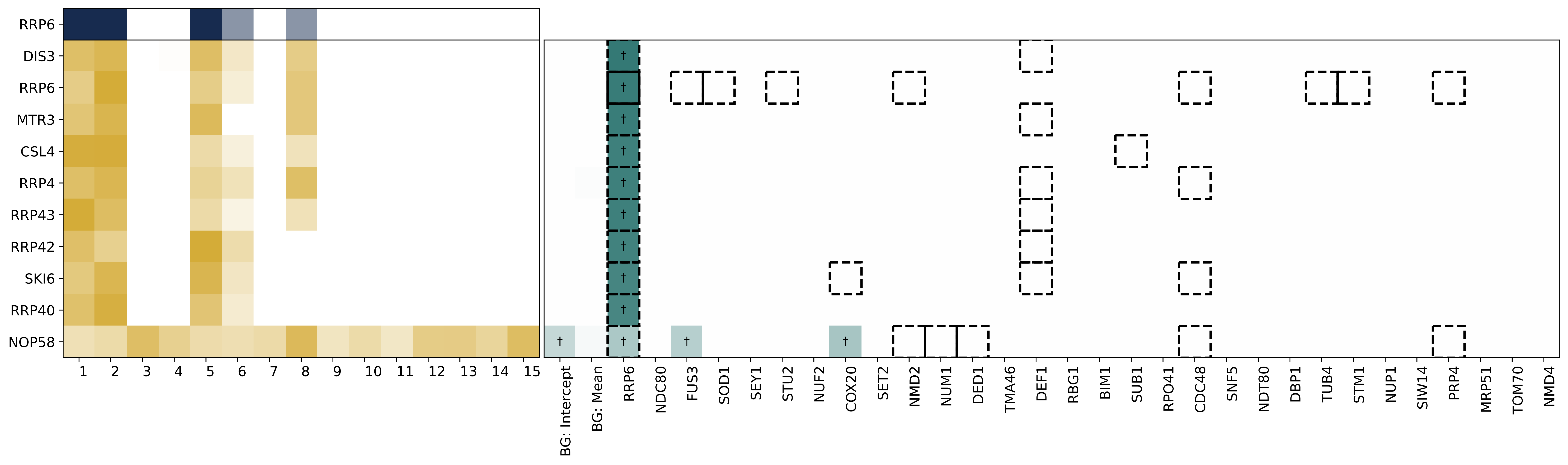

RRP6
DIS3
RRP6
MTR3
CSL4
RRP4
RRP43
RRP42
SKI6
RRP40
NOP58
BG: Intercept
BG: Mean
RRP6
NDC80
FUS3
SOD1
SEY1
STU2
NUF2
COX20
SET2
NMD2
NUM1
DED1
TMA46
DEF1
RBG1
BIM1
SUB1
RPO41
CDC48
SNF5
NDT80
DBP1
TUB4
STM1
NUP1
SIW14
PRP4
MRP51
TOM70
NMD4

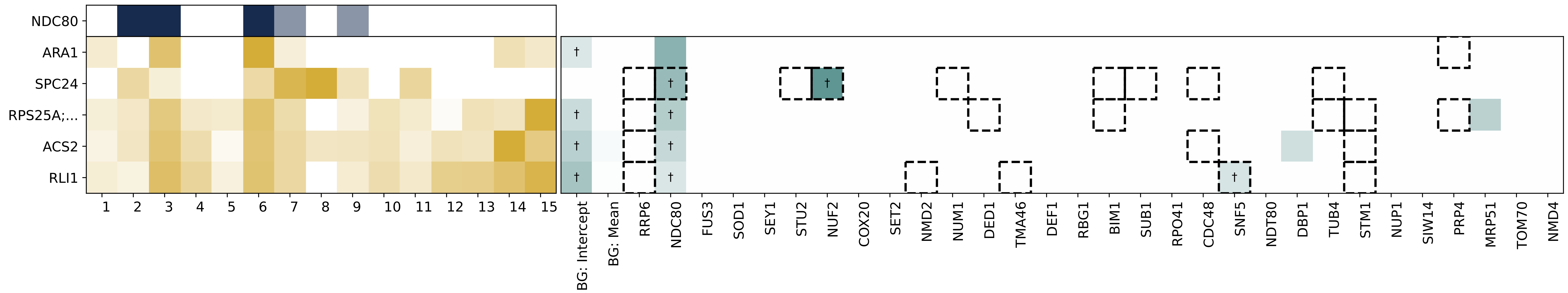

NDC80
ARA1
SPC24
RPS25A;...
ACS2
RLI1
BG: Intercept
BG: Mean
RRP6
NDC80
FUS3
SOD1
SEY1
STU2
NUF2
COX20
SET2
NMD2
NUM1
DED1
TMA46
DEF1
RBG1
BIM1
SUB1
RPO41
CDC48
SNF5
NDT80
DBP1
TUB4
STM1
NUP1
SIW14
PRP4
MRP51
TOM70
NMD4

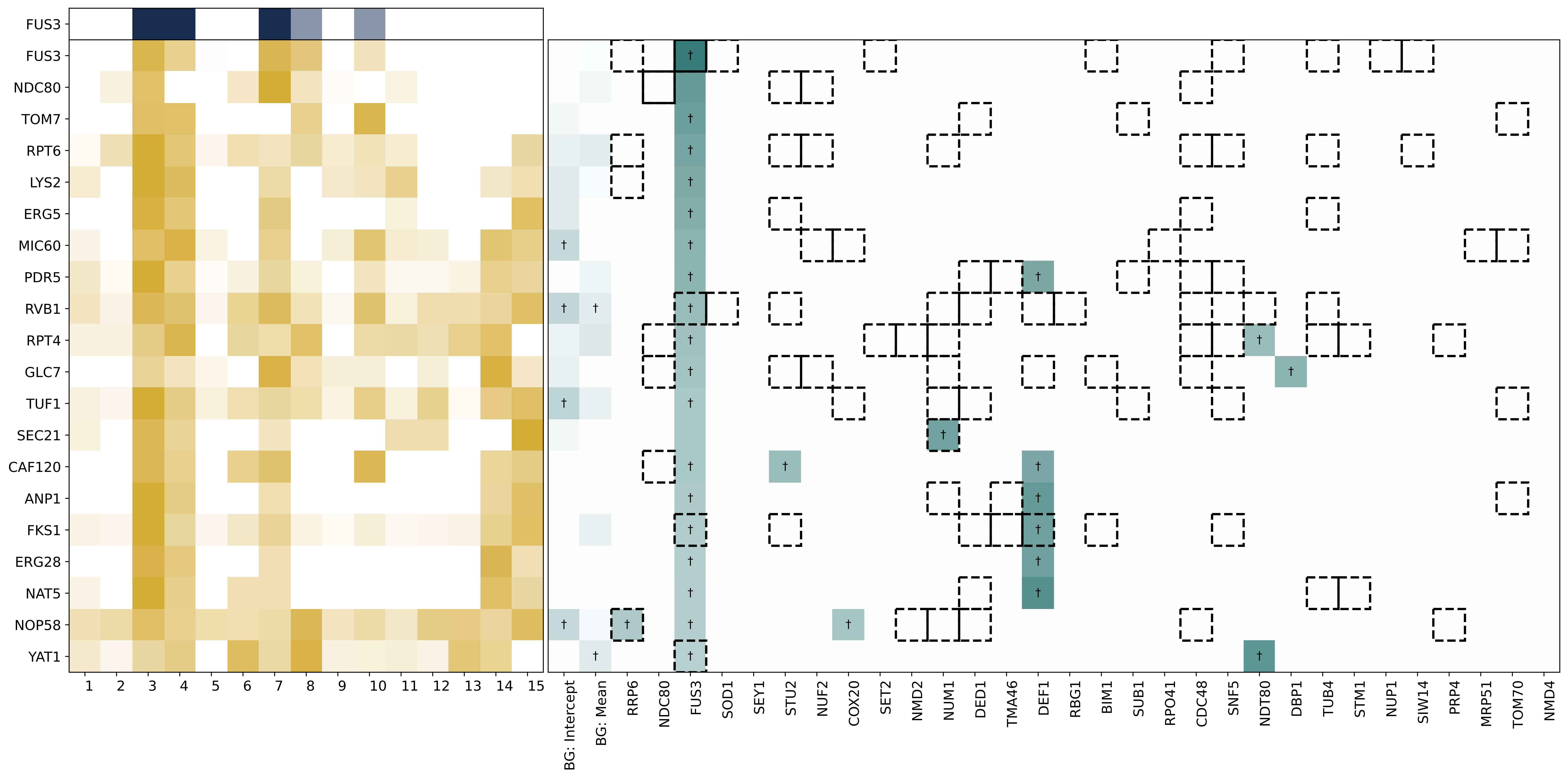

FUS3
FUS3
NDC80
TOM7
RPT6
LYS2
ERG5
MIC60
PDR5
RVB1
RPT4
GLC7
TUF1
SEC21
CAF120
ANP1
FKS1
ERG28
NAT5
NOP58
YAT1
1 2 3 4 5 6 7 8 9 10 11 12 13 14 15
BG: Intercept
BG: Mean
RRP6
NDC80
FUS3
SOD1
SEY1
STU2
NUF2
COX20
SET2
NMD2
NUM1
DED1
TMA46
DEF1
RBG1
BIM1
SUB1
RPO41
CDC48
SNF5
NDT80
DBP1
TUB4
STM1
NUP1
SIW14
PRP4
MRP51
TOM70
NMD4

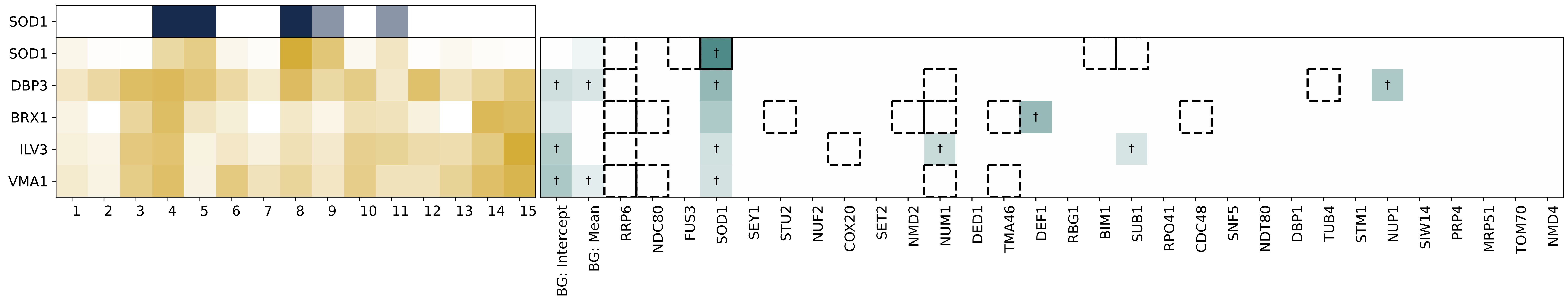

SOD1
SOD1
DBP3
BRX1
ILV3
VMA1
1 2 3 4 5 6 7 8 9 10 11 12 13 14 15
BG: Intercept
BG: Mean
RRP6
NDC80
FUS3
SOD1
SEY1
STU2
NUF2
COX20
SET2
NMD2
NUM1
DED1
TMA46
DEF1
RBG1
BIM1
SUB1
RPO41
CDC48
SNF5
NDT80
DBP1
TUB4
STM1
NUP1
SIW14
PRP4
MRP51
TOM70
NMD4
†

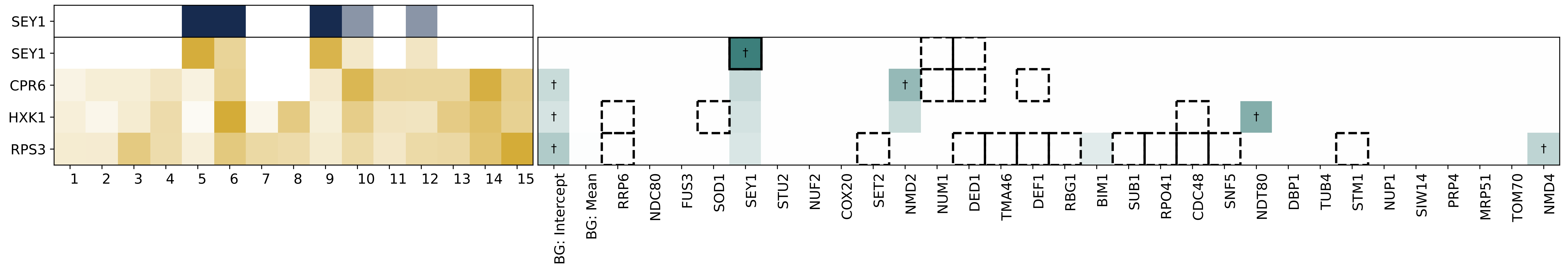

SEY1
SEY1
CPR6
HXK1
RPS3
BG: Intercept
BG: Mean
RRP6
NDC80
FUS3
SOD1
SEY1
STU2
NUF2
COX20
SET2
NMD2
NUM1
DED1
TMA46
DEF1
RBG1
BIM1
SUB1
RPO41
CDC48
SNF5
NDT80
DBP1
TUB4
STM1
NUP1
SIW14
PRP4
MRP51
TOM70
NMD4

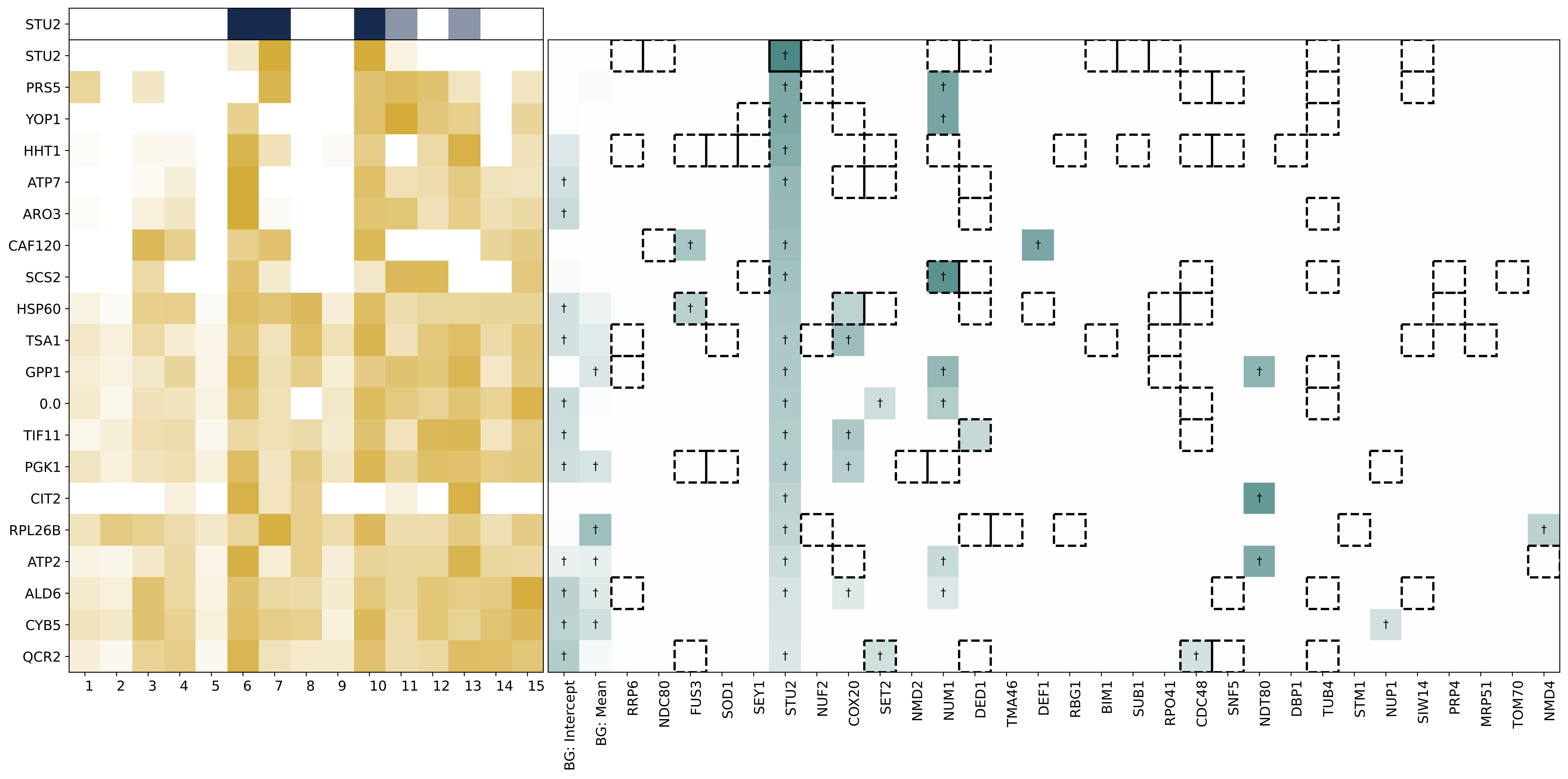

STU2
STU2
PRS5
YOP1
HHT1
ATP7
ARO3
CAF120
SCS2
HSP60
TSA1
GPP1
0.0
TIF11
PGK1
CIT2
RPL26B
ATP2
ALD6
CYB5
QCR2
1 2 3 4 5 6 7 8 9 10 11 12 13 14 15
BG: Intercept
BG: Mean
RRP6
NDC80
FUS3
SOD1
SEY1
STU2
NUF2
COX20
SET2
NMD2
NUM1
DED1
TMA46
DEF1
RBG1
BIM1
SUB1
RPO41
CDC48
SNF5
NDT80
DBP1
TUB4
STM1
NUP1
SIW14
PRP4
MRP51
TOM70
NMD4

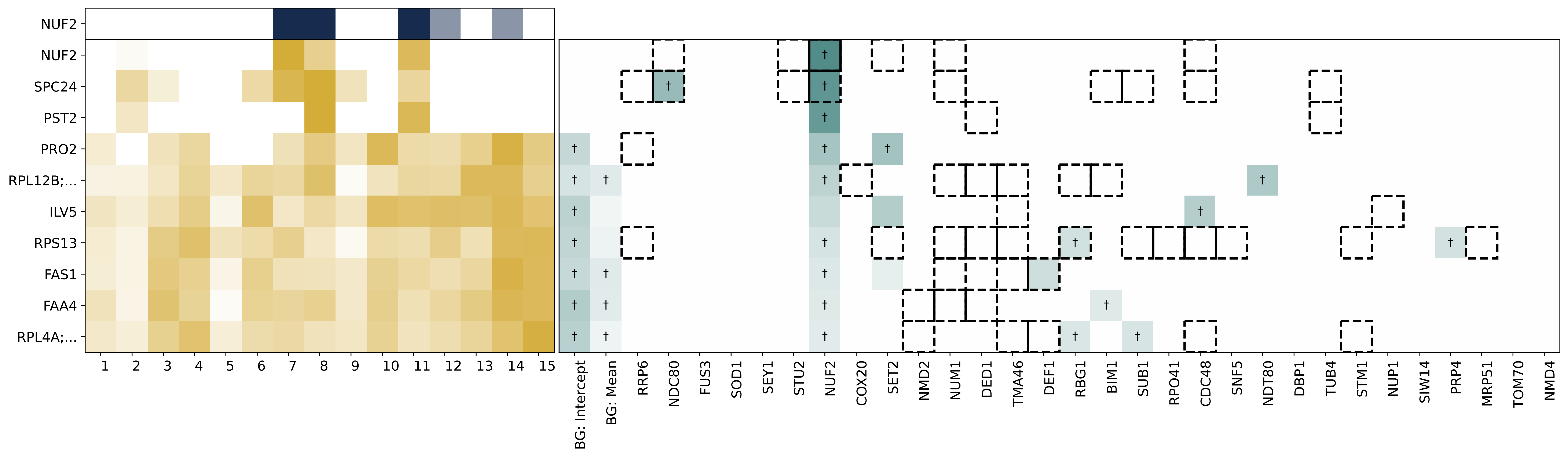
NUF2
NUF2
SPC24
PST2
PRO2
RPL12B;...
ILV5
RPS13
FAS1
FAA4
RPL4A;...
1 2 3 4 5 6 7 8 9 10 11 12 13 14 15
BG: Intercept
BG: Mean
RRP6
NDC80
FUS3
SOD1
SEY1
STU2
NUF2
COX20
SET2
NMD2
NUM1
DED1
TMA46
DEF1
RBG1
BIM1
SUB1
RPO41
CDC48
SNF5
NDT80
DBP1
TUB4
STM1
NUP1
SIW14
PRP4
MRP51
TOM70
NMD4

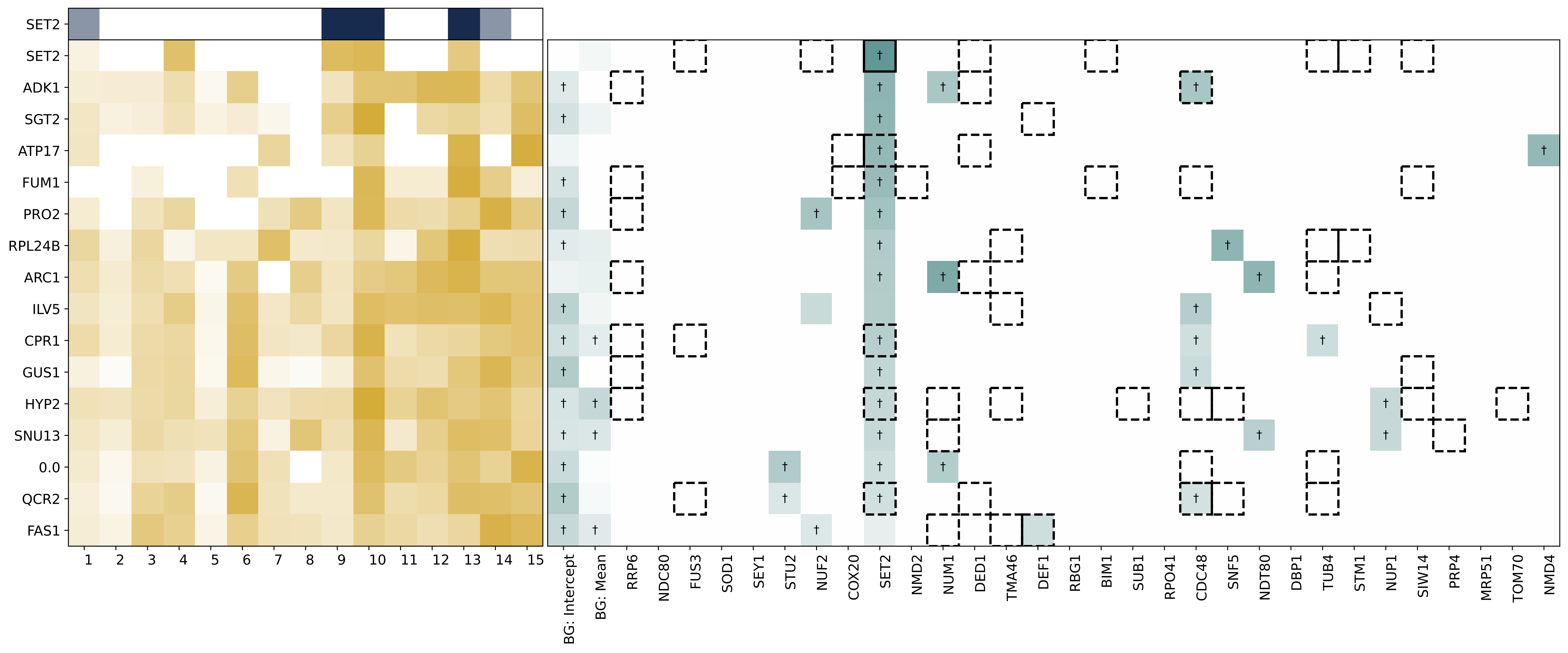
SET2
SET2
ADK1
SGT2
ATP17
FUM1
PRO2
RPL24B
ARC1
ILV5
CPR1
GUS1
HYP2
SNU13
0.0
QCR2
FAS1
BG: Intercept
BG: Mean
RRP6
NDC80
FUS3
SOD1
SEY1
STU2
NUF2
COX20
SET2
NMD2
NUM1
DED1
TMA46
DEF1
RBG1
BIM1
SUB1
RPO41
CDC48
SNF5
NDT80
DBP1
TUB4
STM1
NUP1
SIW14
PRP4
MRP51
TOM70
NMD4

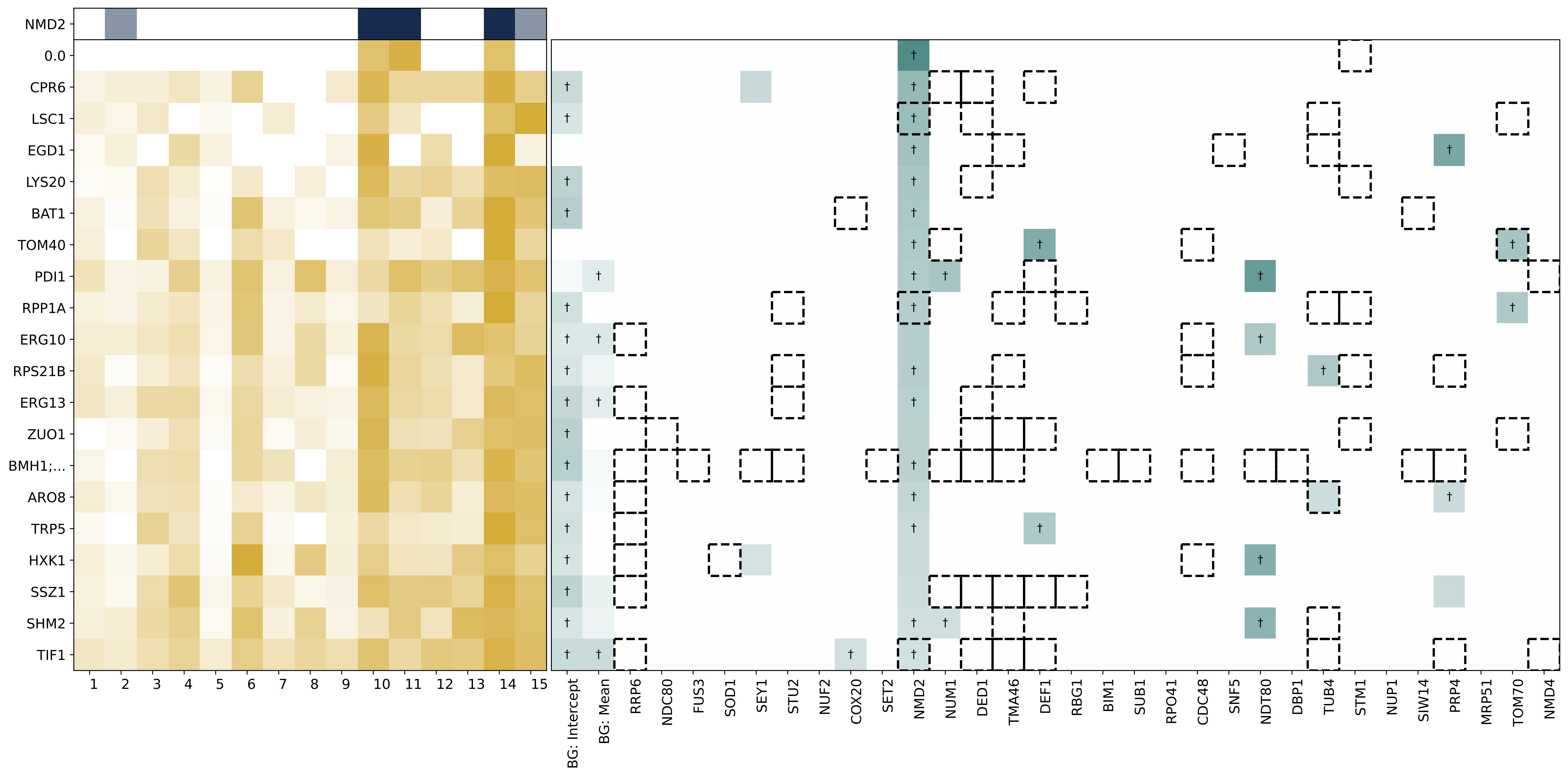

NMD2
0.0
CPR6
LSC1
EGD1
LYS20
BAT1
TOM40
PDI1
RPP1A
ERG10
RPS21B
ERG13
ZUO1
BMH1;...
ARO8
TRP5
HXK1
SSZ1
SHM2
TIF1
1 2 3 4 5 6 7 8 9 10 11 12 13 14 15
BG: Intercept
BG: Mean
RRP6
NDC80
FUS3
SOD1
SEY1
STU2
NUF2
COX20
SET2
NMD2
NUM1
DED1
TMA46
DEF1
RBG1
BIM1
SUB1
RPO41
CDC48
SNF5
NDT80
DBP1
TUB4
STM1
NUP1
SIW14
PRP4
MRP51
TOM70
NMD4

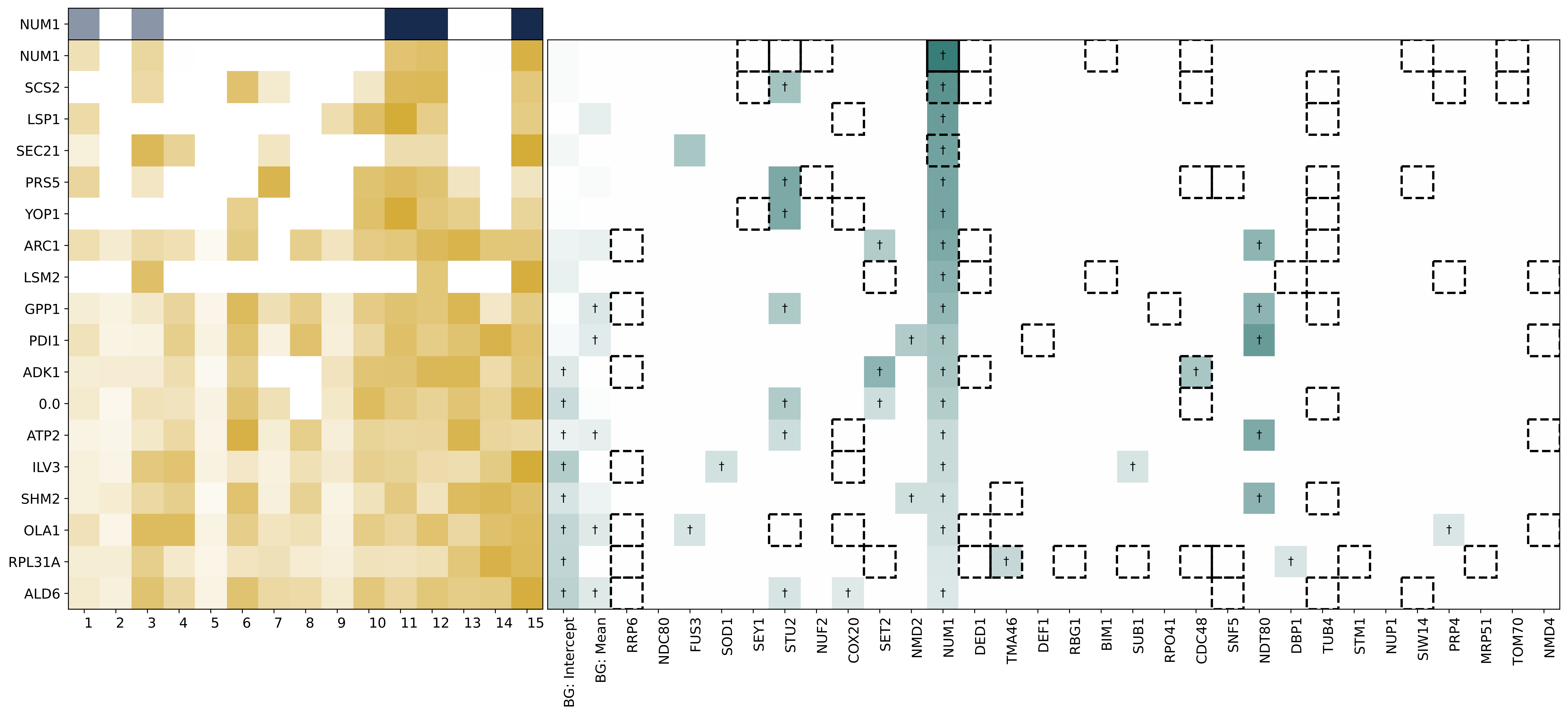
NUM1
NUM1
SCS2
LSP1
SEC21
PRS5
YOP1
ARC1
LSM2
GPP1
PDI1
ADK1
0.0
ATP2
ILV3
SHM2
OLA1
RPL31A
ALD6
1 2 3 4 5 6 7 8 9 10 11 12 13 14 15
BG: Intercept
BG: Mean
RRP6
NDC80
FUS3
SOD1
SEY1
STU2
NUF2
COX20
SET2
NMD2
NUM1
DED1
TMA46
DEF1
RBG1
BIM1
SUB1
RPO41
CDC48
SNF5
NDT80
DBP1
TUB4
STM1
NUP1
SIW14
PRP4
MRP51
TOM70
NMD4

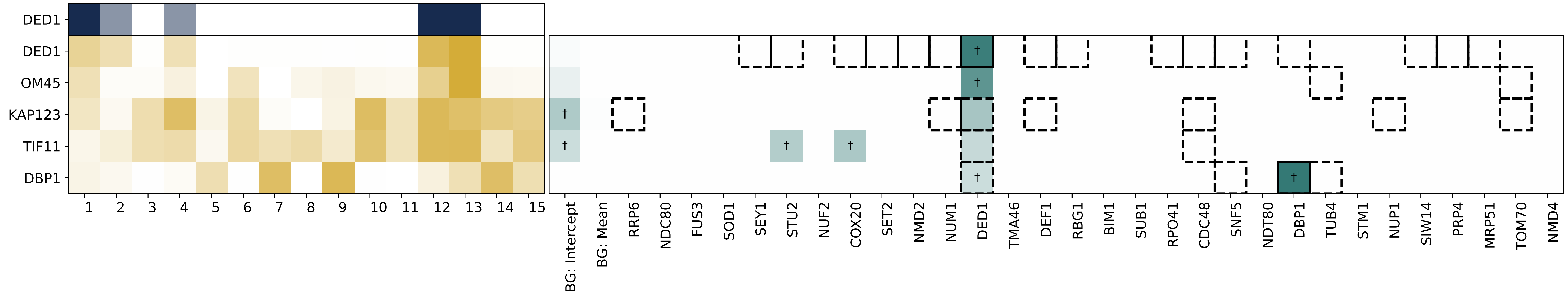

DED1
DED1
OM45
KAP123
TIF11
DBP1
1 2 3 4 5 6 7 8 9 10 11 12 13 14 15
BG: Intercept
BG: Mean
RRP6
NDC80
FUS3
SOD1
SEY1
STU2
NUF2
COX20
SET2
NMD2
NUM1
DED1
TMA46
DEF1
RBG1
BIM1
SUB1
RPO41
CDC48
SNF5
NDT80
DBP1
TUB4
STM1
NUP1
SIW14
PRP4
MRP51
TOM70
NMD4

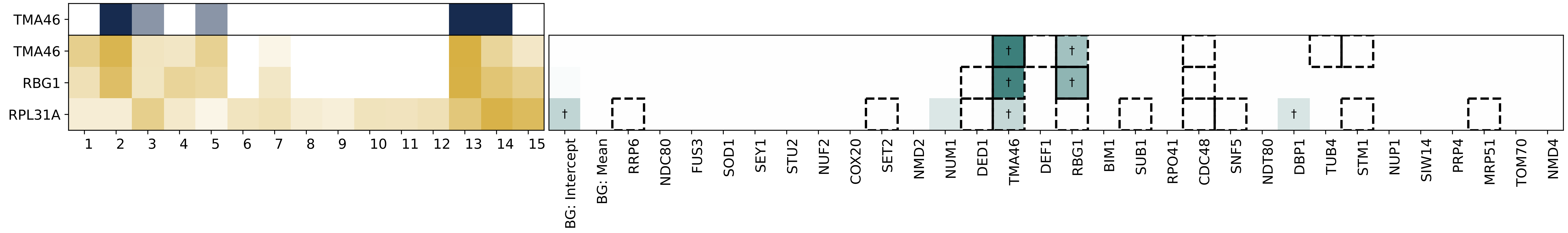

TMA46
TMA46
RBG1
RPL31A
†
BG: Intercept
BG: Mean
RRP6
NDC80
FUS3
SOD1
SEY1
STU2
NUF2
COX20
SET2
NMD2
NUM1
DED1
TMA46
DEF1
RBG1
BIM1
SUB1
RPO41
CDC48
SNF5
NDT80
DBP1
TUB4
STM1
NUP1
SIW14
PRP4
MRP51
TOM70
NMD4

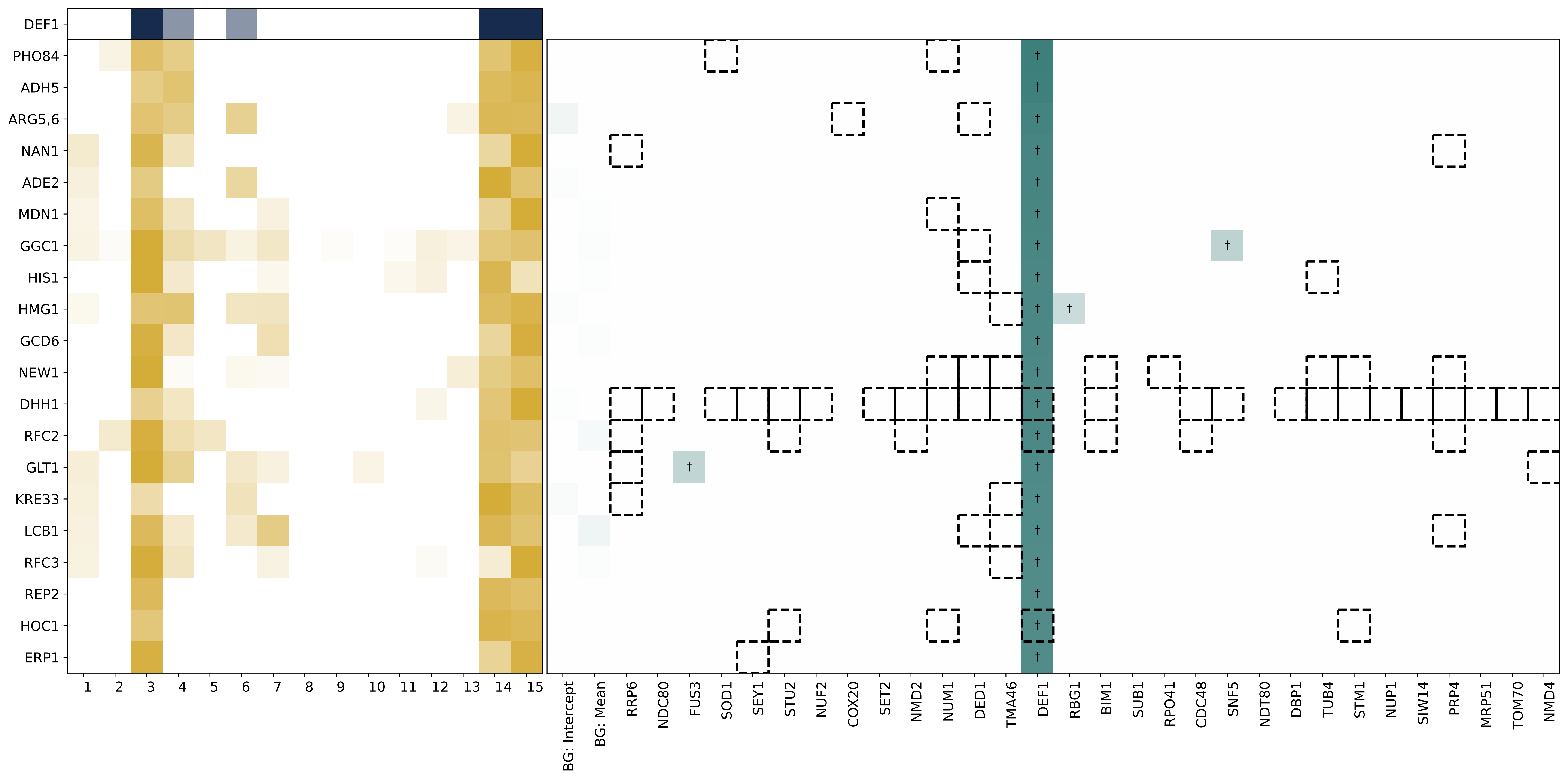

DEF1
PHO84
ADH5
ARG5,6
NAN1
ADE2
MDN1
GGC1
HIS1
HMG1
GCD6
NEW1
DHH1
RFC2
GLT1
KRE33
LCB1
RFC3
REP2
HOC1
ERP1
BG: Intercept
BG: Mean
RRP6
NDC80
FUS3
SOD1
SEY1
STU2
NUF2
COX20
SET2
NMD2
NUM1
DED1
TMA46
DEF1
RBG1
BIM1
SUB1
RPO41
CDC48
SNF5
NDT80
DBP1
TUB4
STM1
NUP1
SIW14
PRP4
MRP51
TOM70
NMD4

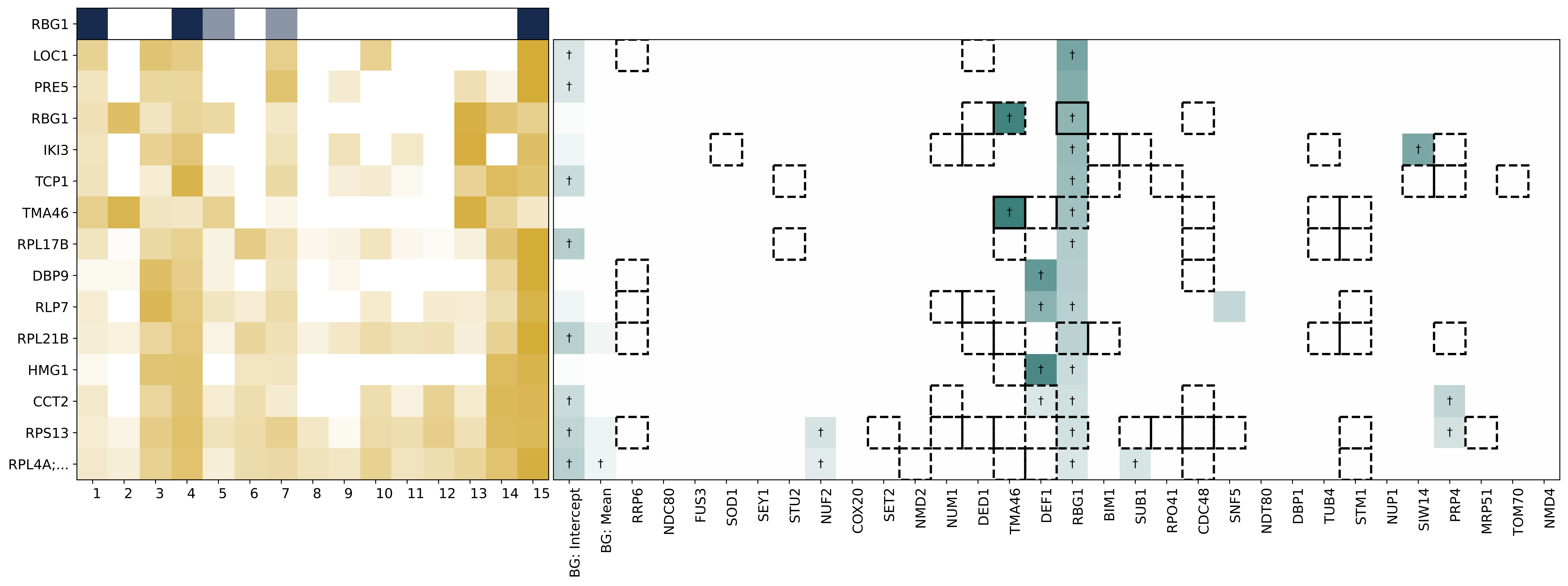

RBG1
LOC1
PRE5
RBG1
IKI3
TCP1
TMA46
RPL17B
DBP9
RLP7
RPL21B
HMG1
CCT2
RPS13
RPL4A;...
1 2 3 4 5 6 7 8 9 10 11 12 13 14 15
BG: Intercept
BG: Mean
RRP6
NDC80
FUS3
SOD1
SEY1
STU2
NUF2
COX20
SET2
NMD2
NUM1
DED1
TMA46
DEF1
RBG1
BIM1
SUB1
RPO41
CDC48
SNF5
NDT80
DBP1
TUB4
STM1
NUP1
SIW14
PRP4
MRP51
TOM70
NMD4

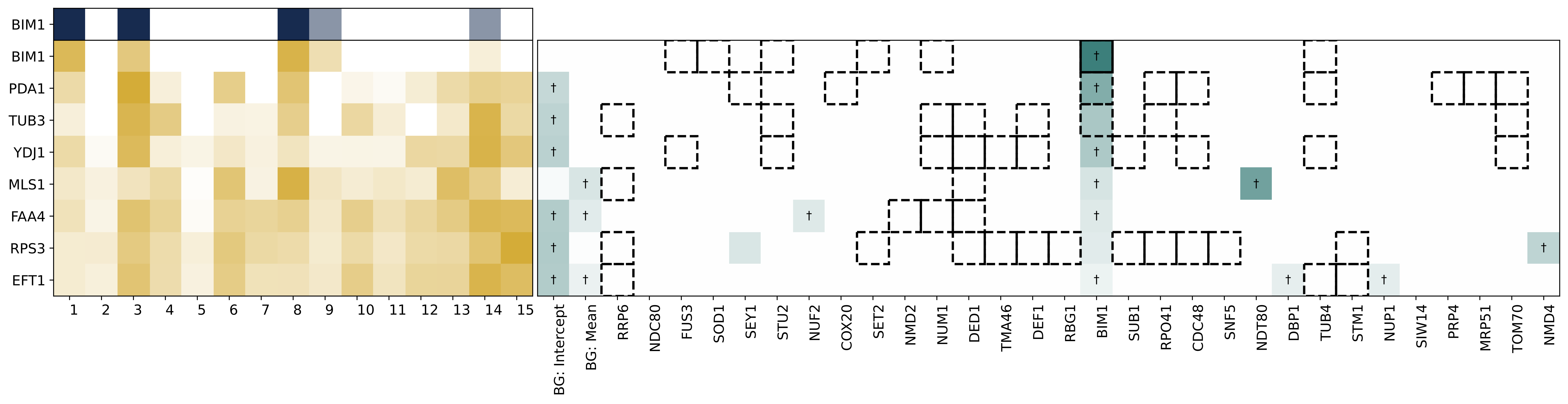

BIM1
BIM1
PDA1
TUB3
YDJ1
MLS1
FAA4
RPS3
EFT1
BG: Intercept
BG: Mean
RRP6
NDC80
FUS3
SOD1
SEY1
STU2
NUF2
COX20
SET2
NMD2
NUM1
DED1
TMA46
DEF1
RBG1
BIM1
SUB1
RPO41
CDC48
SNF5
NDT80
DBP1
TUB4
STM1
NUP1
SIW14
PRP4
MRP51
TOM70
NMD4

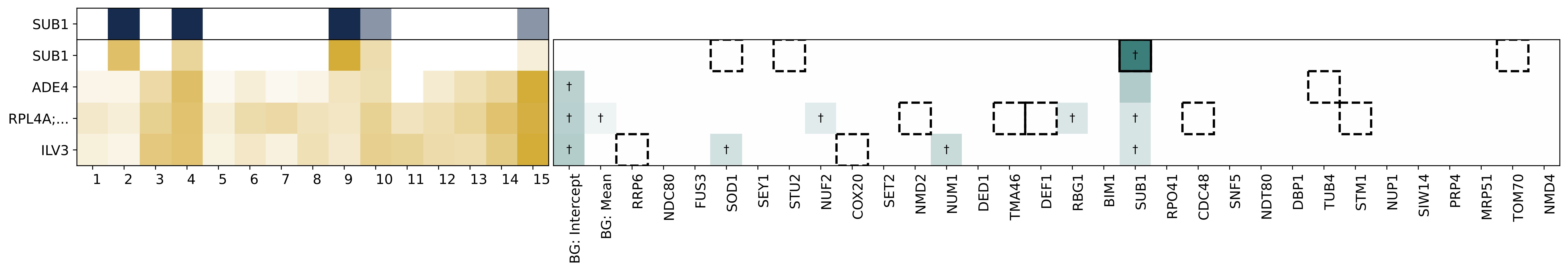
SUB1
SUB1
ADE4
RPL4A;...
ILV3
1 2 3 4 5 6 7 8 9 10 11 12 13 14 15
BG: Intercept
BG: Mean
RRP6
NDC80
FUS3
SOD1
SEY1
STU2
NUF2
COX20
SET2
NMD2
NUM1
DED1
TMA46
DEF1
RBG1
BIM1
SUB1
RPO41
CDC48
SNF5
NDT80
DBP1
TUB4
STM1
NUP1
SIW14
PRP4
MRP51
TOM70
NMD4

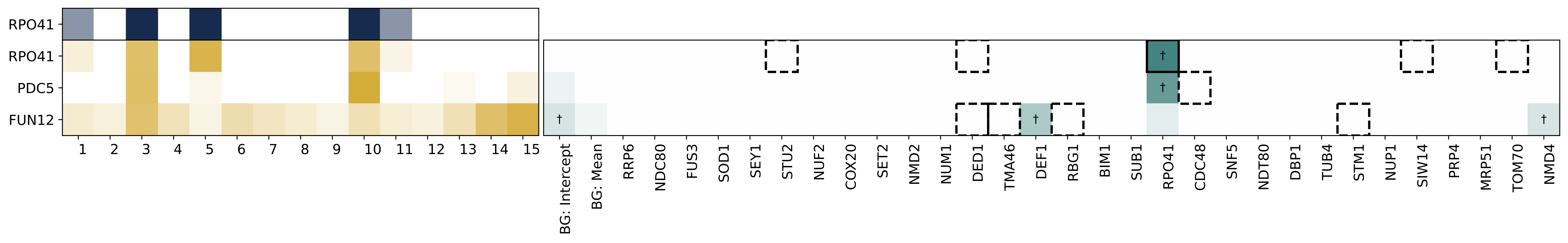

RPO41
RPO41
PDC5
FUN12
BG: Intercept
BG: Mean
RRP6
NDC80
FUS3
SOD1
SEY1
STU2
NUF2
COX20
SET2
NMD2
NUM1
DED1
TMA46
DEF1
RBG1
BIM1
SUB1
RPO41
CDC48
SNF5
NDT80
DBP1
TUB4
STM1
NUP1
SIW14
PRP4
MRP51
TOM70
NMD4

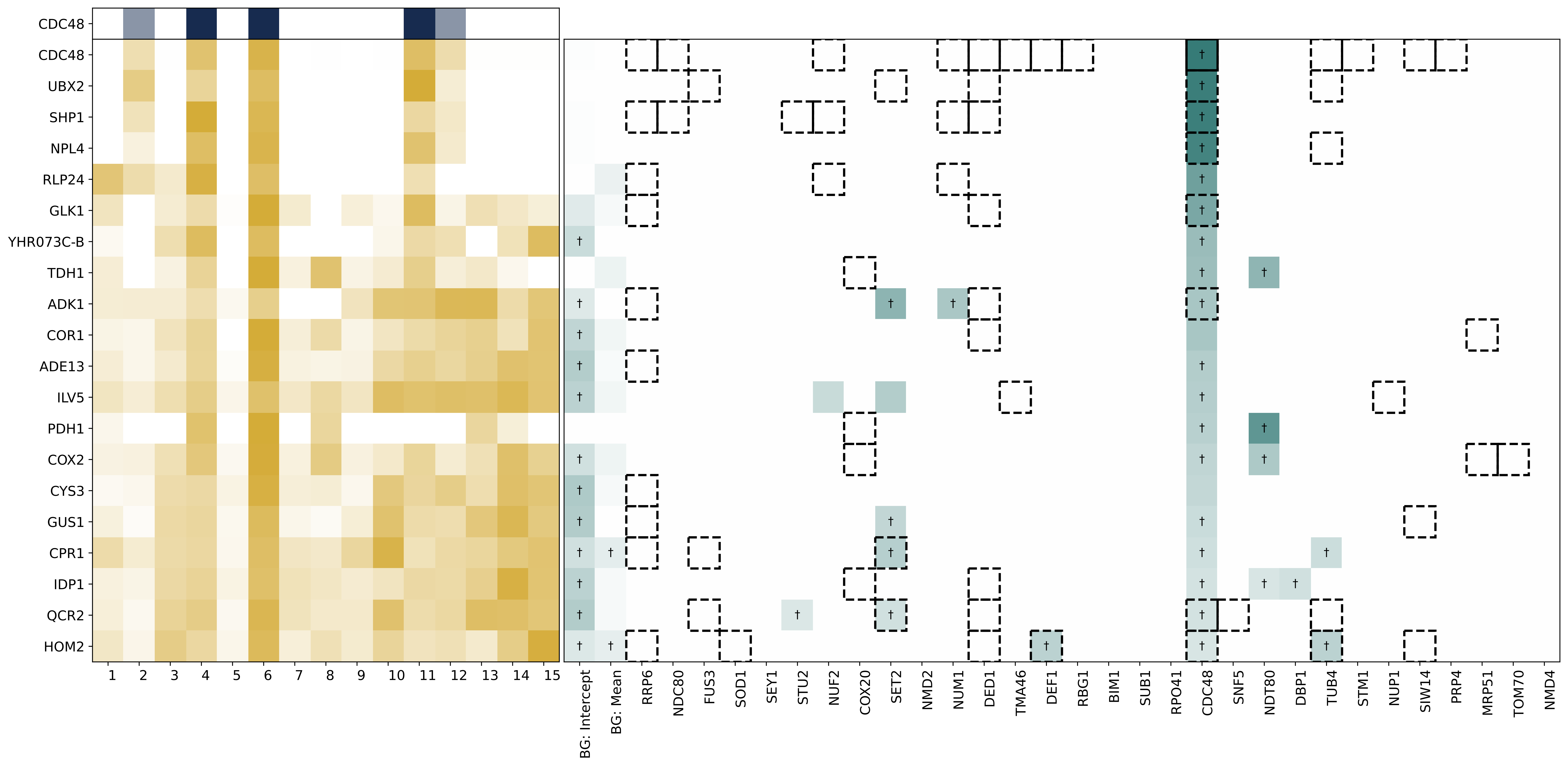

CDC48
CDC48
UBX2
SHP1
NPL4
RLP24
GLK1
YHR073C-B
TDH1
ADK1
COR1
ADE13
ILV5
PDH1
COX2
CYS3
GUS1
CPR1
IDP1
QCR2
HOM2
BG: Intercept
BG: Mean
RRP6
NDC80
FUS3
SOD1
SEY1
STU2
NUF2
COX20
SET2
NMD2
NUM1
DED1
TMA46
DEF1
RBG1
BIM1
SUB1
RPO41
CDC48
SNF5
NDT80
DBP1
TUB4
STM1
NUP1
SIW14
PRP4
MRP51
TOM70
NMD4

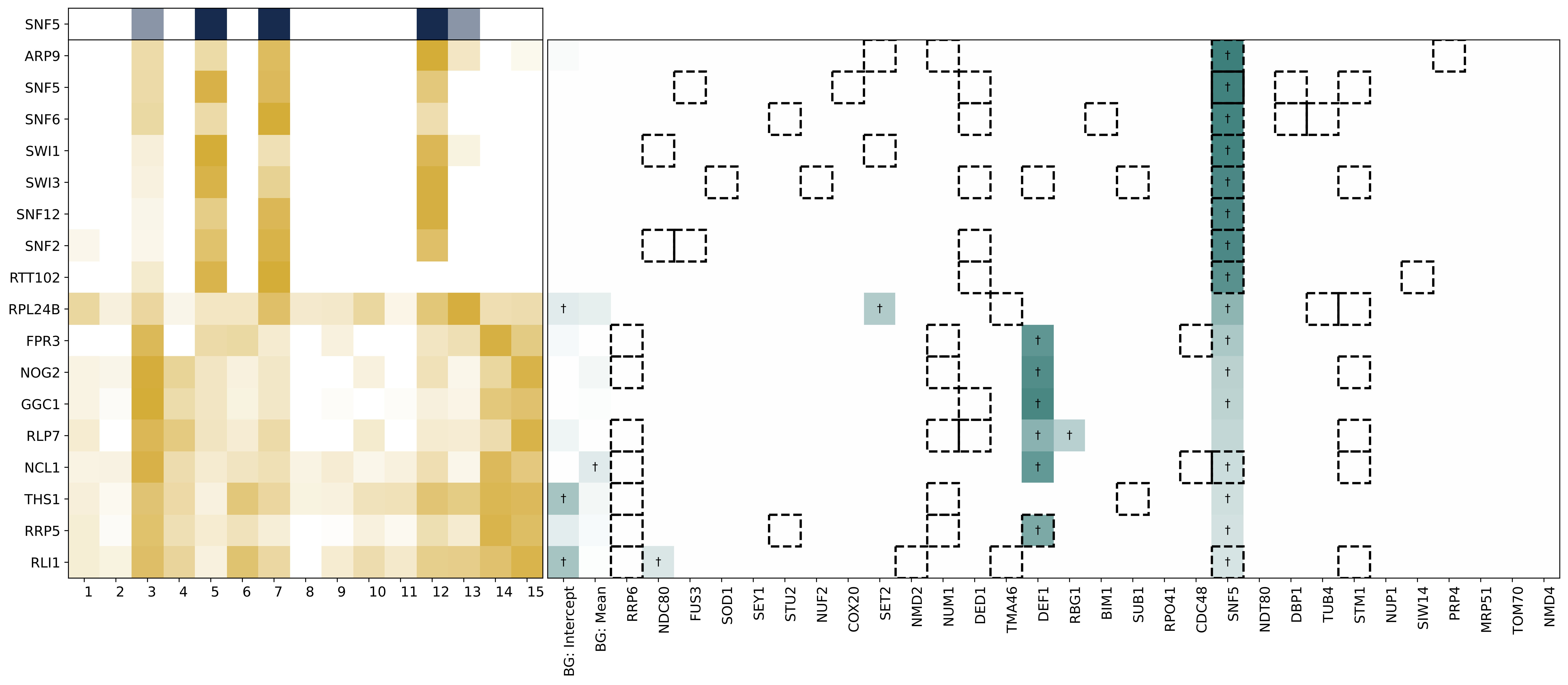
SNF5
ARP9
SNF5
SNF6
SWI1
SWI3
SNF12
SNF2
RTT102
RPL24B
FPR3
NOG2
GGC1
RLP7
NCL1
THS1
RRP5
RLI1
1 2 3 4 5 6 7 8 9 10 11 12 13 14 15
BG: Intercept
BG: Mean
RRP6
NDC80
FUS3
SOD1
SEY1
STU2
NUF2
COX20
SET2
NMD2
NUM1
DED1
TMA46
DEF1
RBG1
BIM1
SUB1
RPO41
CDC48
SNF5
NDT80
DBP1
TUB4
STM1
NUP1
SIW14
PRP4
MRP51
TOM70
NMD4

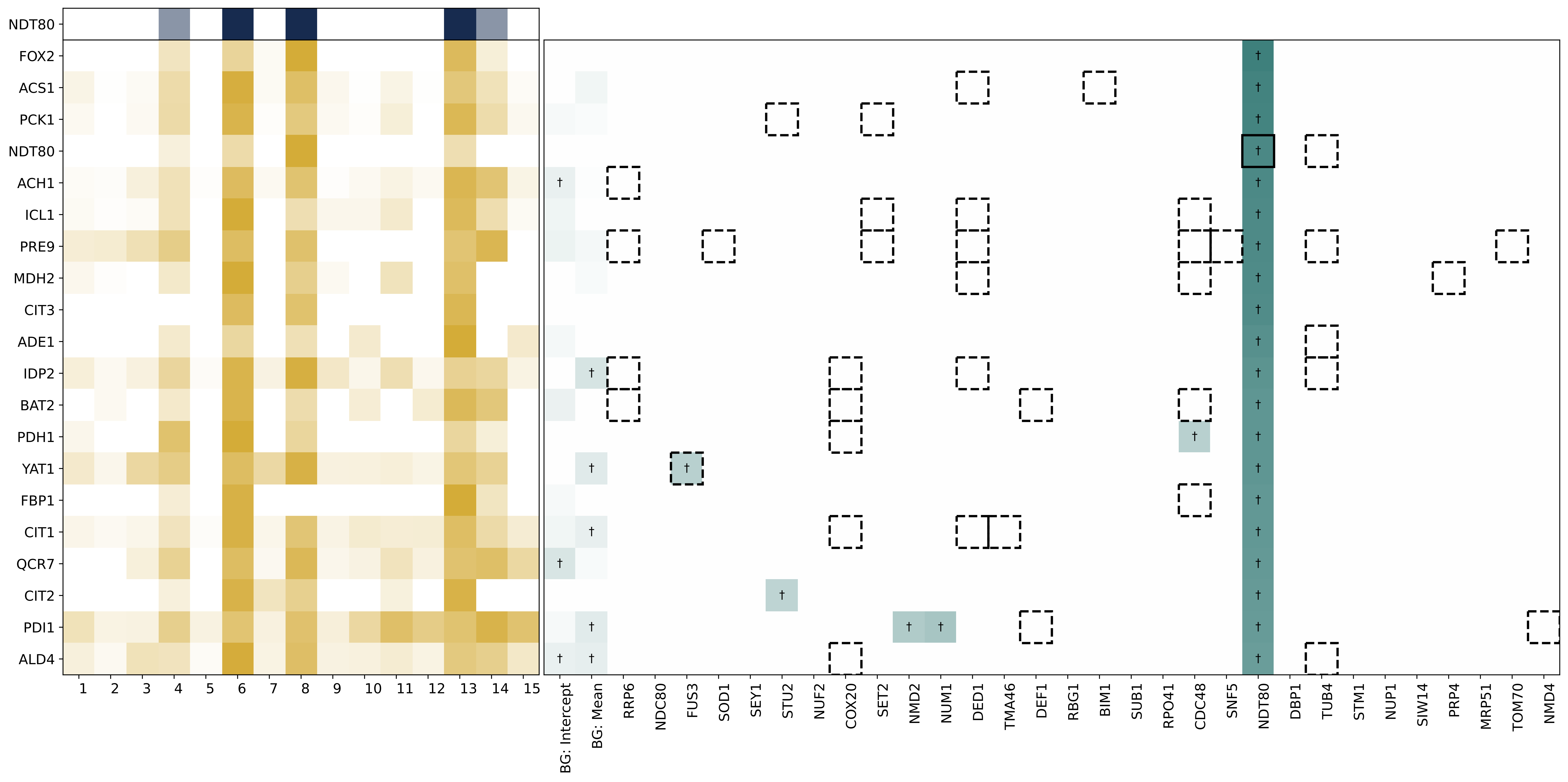

NDT80
FOX2
ACS1
PCK1
NDT80
ACH1
ICL1
PRE9
MDH2
CIT3
ADE1
IDP2
BAT2
PDH1
YAT1
FBP1
CIT1
QCR7
CIT2
PDI1
ALD4
1 2 3 4 5 6 7 8 9 10 11 12 13 14 15
BG: Intercept
BG: Mean
RRP6
NDC80
FUS3
SOD1
SEY1
STU2
NUF2
COX20
SET2
NMD2
NUM1
DED1
TMA46
DEF1
RBG1
BIM1
SUB1
RPO41
CDC48
SNF5
NDT80
DBP1
TUB4
STM1
NUP1
SIW14
PRP4
MRP51
TOM70
NMD4

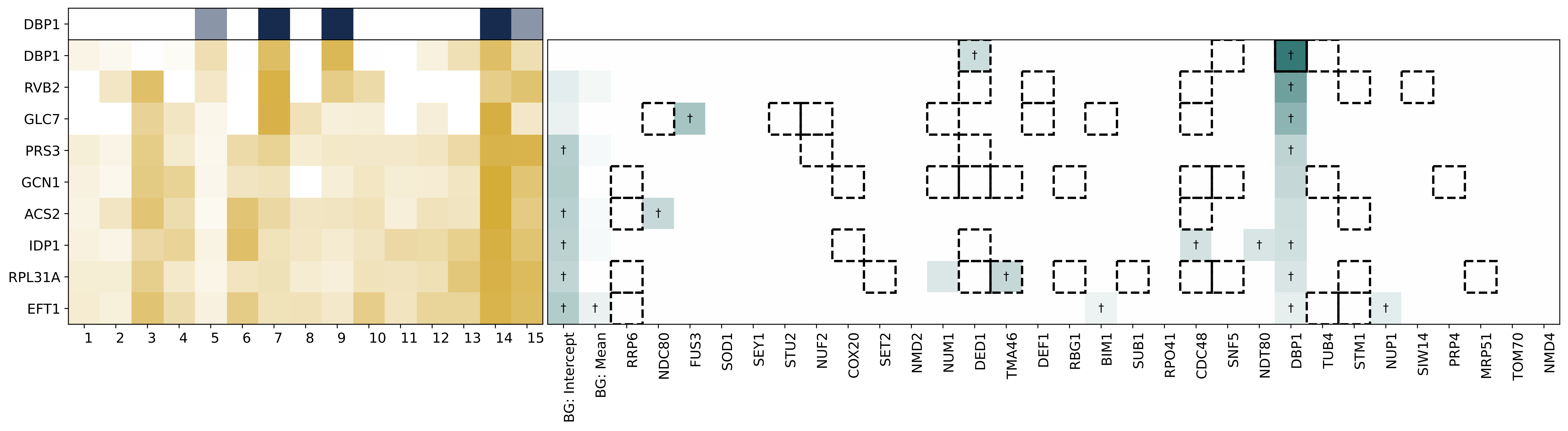

DBP1
DBP1
RVB2
GLC7
PRS3
GCN1
ACS2
IDP1
RPL31A
EFT1
1 2 3 4 5 6 7 8 9 10 11 12 13 14 15
BG: Intercept
BG: Mean
RRP6
NDC80
FUS3
SOD1
SEY1
STU2
NUF2
COX20
SET2
NMD2
NUM1
DED1
TMA46
DEF1
RBG1
BIM1
SUB1
RPO41
CDC48
SNF5
NDT80
DBP1
TUB4
STM1
NUP1
SIW14
PRP4
MRP51
TOM70
NMD4

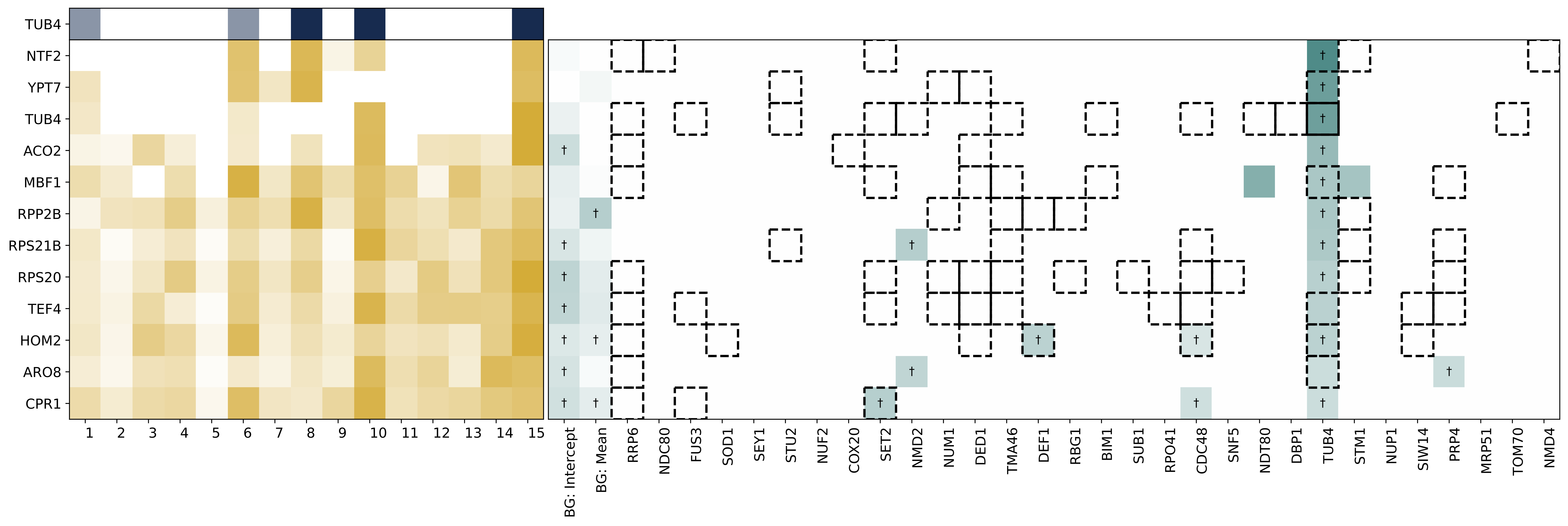

TUB4
NTF2
YPT7
TUB4
ACO2
MBF1
RPP2B
RPS21B
RPS20
TEF4
HOM2
ARO8
CPR1
BG: Intercept
BG: Mean
RRP6
NDC80
FUS3
SOD1
SEY1
STU2
NUF2
COX20
SET2
NMD2
NUM1
DED1
TMA46
DEF1
RBG1
BIM1
SUB1
RPO41
CDC48
SNF5
NDT80
DBP1
TUB4
STM1
NUP1
SIW14
PRP4
MRP51
TOM70
NMD4

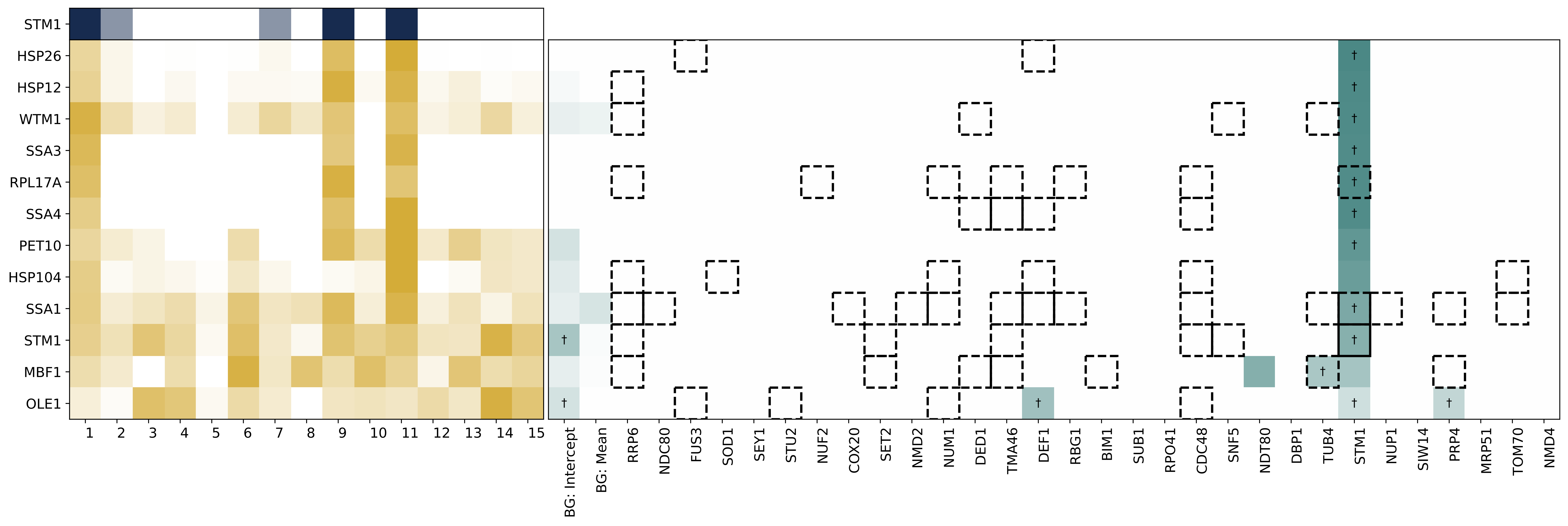
STM1
HSP26
HSP12
WTM1
SSA3
RPL17A
SSA4
PET10
HSP104
SSA1
STM1
MBF1
OLE1
1 2 3 4 5 6 7 8 9 10 11 12 13 14 15
BG: Intercept
BG: Mean
RRP6
NDC80
FUS3
SOD1
SEY1
STU2
NUF2
COX20
SET2
NMD2
NUM1
DED1
TMA46
DEF1
RBG1
BIM1
SUB1
RPO41
CDC48
SNF5
NDT80
DBP1
TUB4
STM1
NUP1
SIW14
PRP4
MRP51
TOM70
NMD4

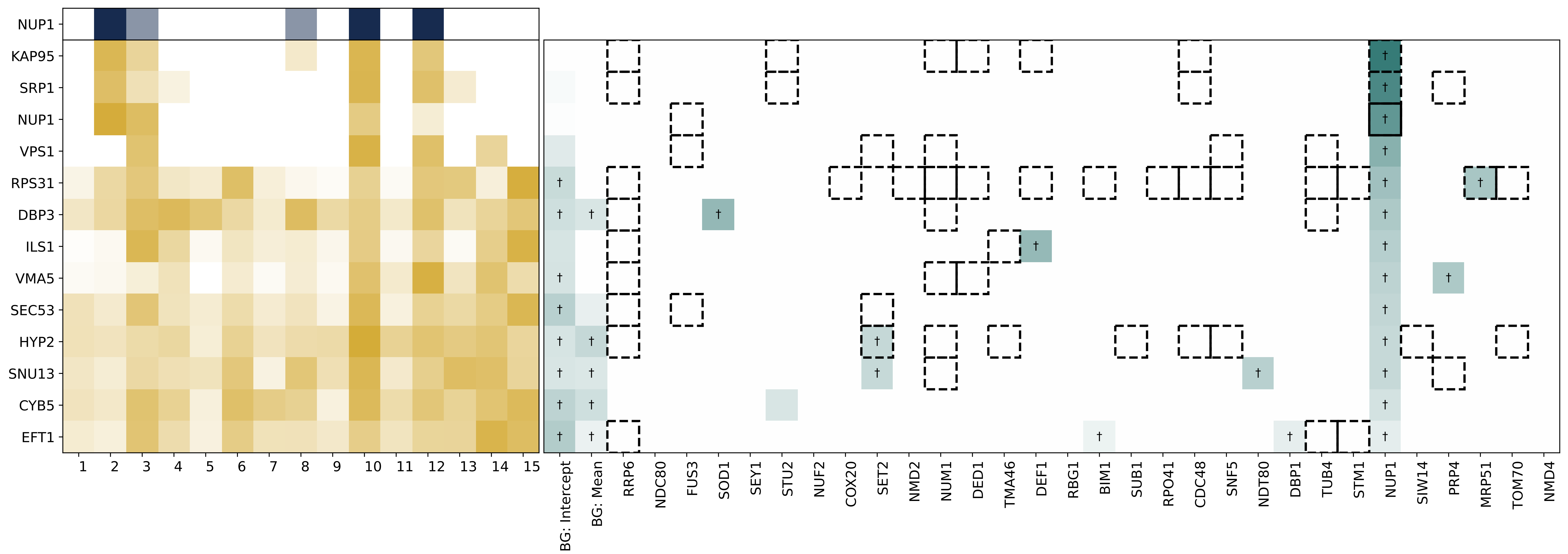

NUP1
KAP95
SRP1
NUP1
VPS1
RPS31
DBP3
ILS1
VMA5
SEC53
HYP2
SNU13
CYB5
EFT1
BG: Intercept
BG: Mean
RRP6
NDC80
FUS3
SOD1
SEY1
STU2
NUF2
COX20
SET2
NMD2
NUM1
DED1
TMA46
DEF1
RBG1
BIM1
SUB1
RPO41
CDC48
SNF5
NDT80
DBP1
TUB4
STM1
NUP1
SIW14
PRP4
MRP51
TOM70
NMD4

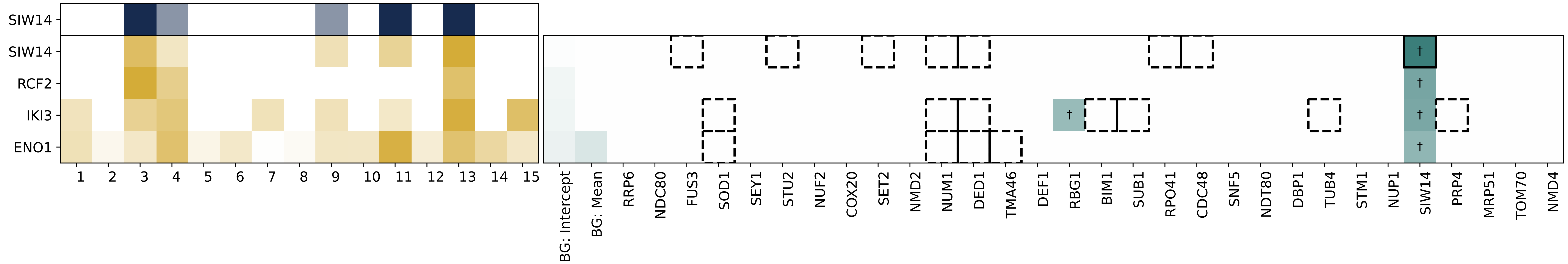
SIW14
SIW14
RCF2
IKI3
ENO1
1 2 3 4 5 6 7 8 9 10 11 12 13 14 15
BG: Intercept
BG: Mean
RRP6
NDC80
FUS3
SOD1
SEY1
STU2
NUF2
COX20
SET2
NMD2
NUM1
DED1
TMA46
DEF1
RBG1
BIM1
SUB1
RPO41
CDC48
SNF5
NDT80
DBP1
TUB4
STM1
NUP1
SIW14
PRP4
MRP51
TOM70
NMD4

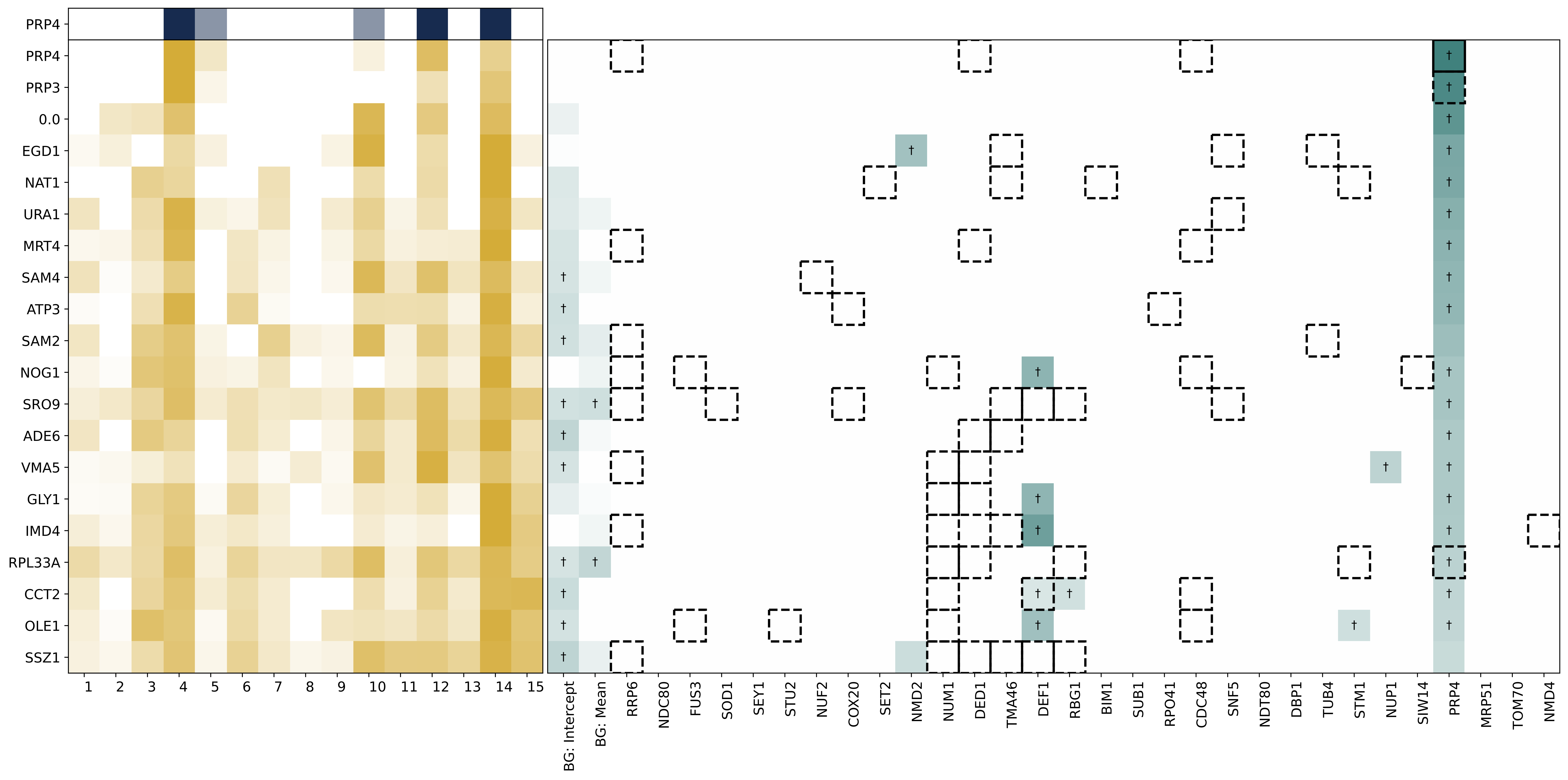
PRP4
PRP4
PRP3
0.0
EGD1
NAT1
URA1
MRT4
SAM4
ATP3
SAM2
NOG1
SRO9
ADE6
VMA5
GLY1
IMD4
RPL33A
CCT2
OLE1
SSZ1
BG: Intercept
BG: Mean
RRP6
NDC80
FUS3
SOD1
SEY1
STU2
NUF2
COX20
SET2
NMD2
NUM1
DED1
TMA46
DEF1
RBG1
BIM1
SUB1
RPO41
CDC48
SNF5
NDT80
DBP1
TUB4
STM1
NUP1
SIW14
PRP4
MRP51
TOM70
NMD4

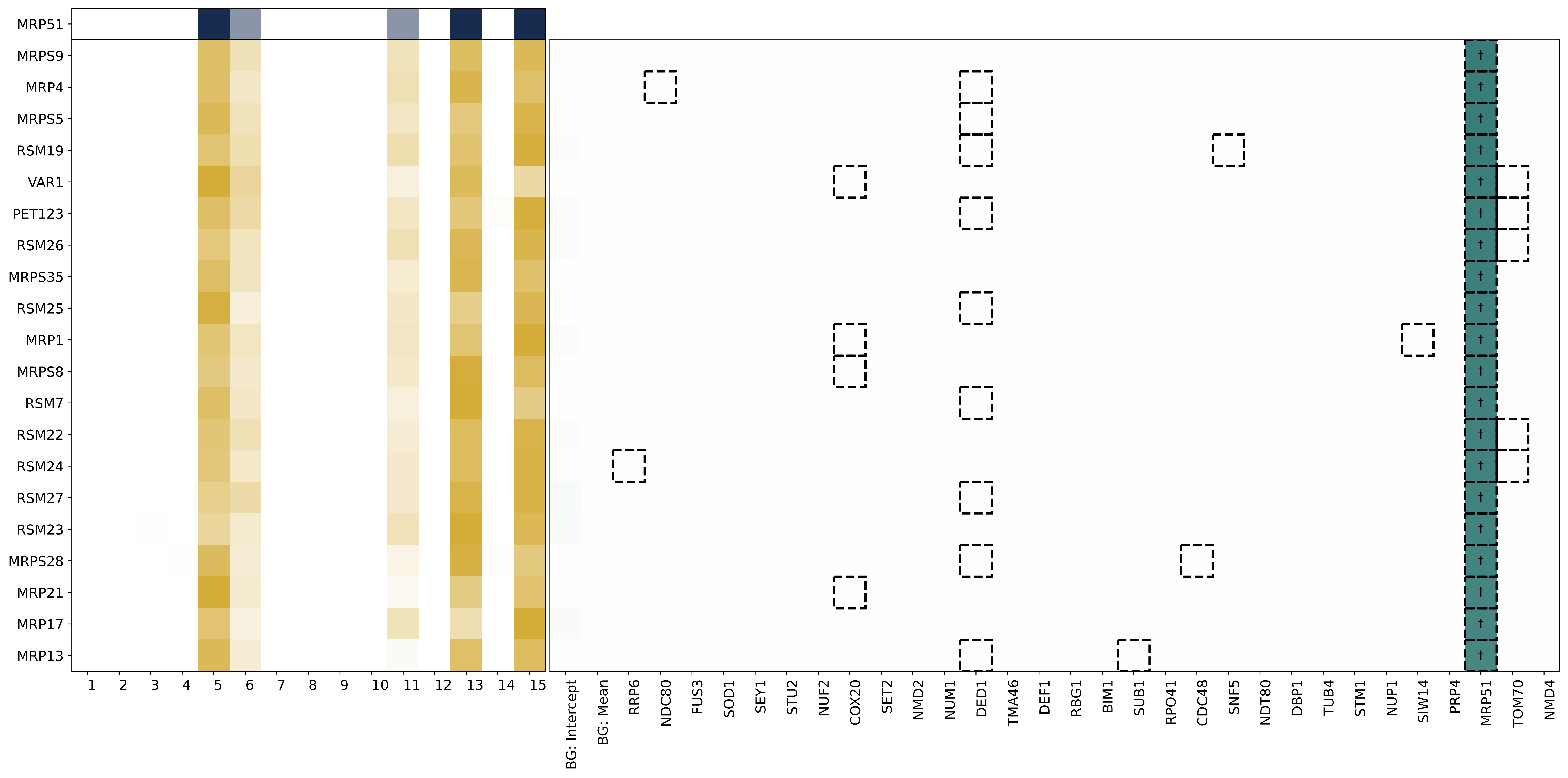

MRP51
MRPS9
MRP4
MRPS5
RSM19
VAR1
PET123
RSM26
MRPS35
RSM25
MRP1
MRPS8
RSM7
RSM22
RSM24
RSM27
RSM23
MRPS28
MRP21
MRP17
MRP13
1 2 3 4 5 6 7 8 9 10 11 12 13 14 15
BG: Intercept
BG: Mean
RRP6
NDC80
FUS3
SOD1
SEY1
STU2
NUF2
COX20
SET2
NMD2
NUM1
DED1
TMA46
DEF1
RBG1
BIM1
SUB1
RPO41
CDC48
SNF5
NDT80
DBP1
TUB4
STM1
NUP1
SIW14
PRP4
MRP51
TOM70
NMD4

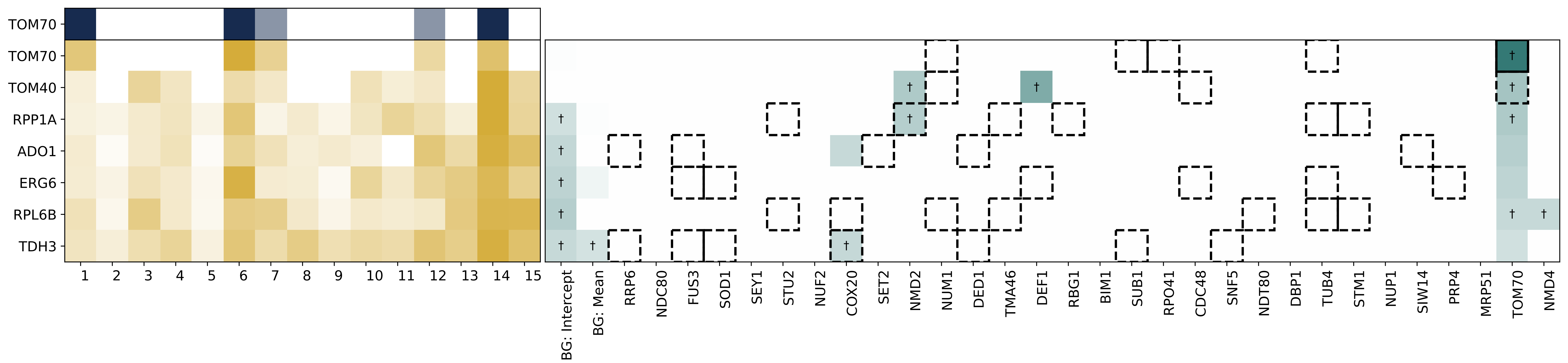
TOM70
TOM70
TOM40
RPP1A
ADO1
ERG6
RPL6B
TDH3
1 2 3 4 5 6 7 8 9 10 11 12 13 14 15
BG: Intercept
BG: Mean
RRP6
NDC80
FUS3
SOD1
SEY1
STU2
NUF2
COX20
SET2
NMD2
NUM1
DED1
TMA46
DEF1
RBG1
BIM1
SUB1
RPO41
CDC48
SNF5
NDT80
DBP1
TUB4
STM1
NUP1
SIW14
PRP4
MRP51
TOM70
NMD4

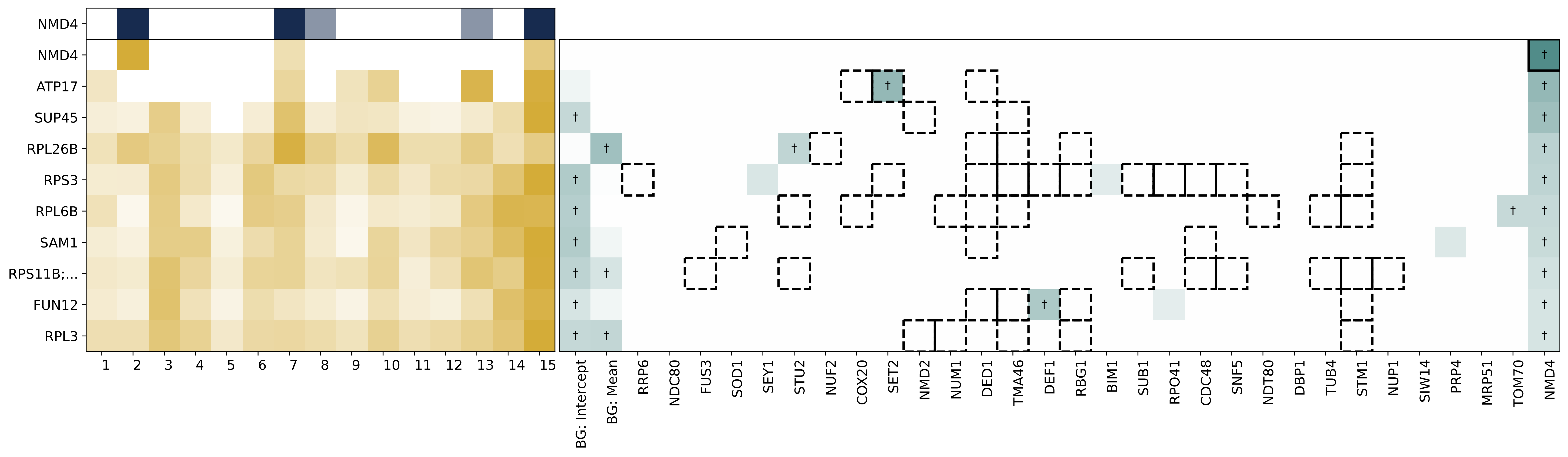
NMD4
NMD4
ATP17
SUP45
RPL26B
RPS3
RPL6B
SAM1
RPS11B;...
FUN12
RPL3
1 2 3 4 5 6 7 8 9 10 11 12 13 14 15
BG: Intercept
BG: Mean
RRP6
NDC80
FUS3
SOD1
SEY1
STU2
NUF2
COX20
SET2
NMD2
NUM1
DED1
TMA46
DEF1
RBG1
BIM1
SUB1
RPO41
CDC48
SNF5
NDT80
DBP1
TUB4
STM1
NUP1
SIW14
PRP4
MRP51
TOM70
NMD4
